# High-quality genome and methylomes illustrate features underlying evolutionary success of oaks

**DOI:** 10.1101/2021.04.09.439191

**Authors:** Victoria L. Sork, Shawn J. Cokus, Sorel T. Fitz-Gibbon, Aleksey V. Zimin, Daniela Puiu, Jesse A. Garcia, Paul F. Gugger, Claudia L. Henriquez, Ying Zhen, Kirk E. Lohmueller, Matteo Pellegrini, Steven L. Salzberg

**Affiliations:** Department of Ecology and Evolutionary Biology, University of California, Los Angeles, CA 90095-1438; Institute of the Environment and Sustainability, University of California, Los Angeles, CA 90095; Department of Molecular, Cell, and Developmental Biology, University of California, Los Angeles, CA 90095-7239; Center for Computational Biology, Whiting School of Engineering, Johns Hopkins University, Baltimore, Maryland 21218; Department of Biomedical Engineering, Johns Hopkins University, Baltimore, Maryland 21218; University of Maryland Center for Environmental Science, Appalachian Laboratory, Frostburg, MD 21532; Department of Human Genetics, David Geffen School of Medicine, University of California, Los Angeles, CA 90095; Departments of Biomedical Engineering, Computer Science, and Biostatistics, Johns Hopkins University, Baltimore, Maryland 21218

## Abstract

The genus *Quercus*, which emerged ~55 million years ago during globally warm temperatures, diversified into ~450 species. We present a high-quality *de novo* genome assembly of a California endemic oak, *Quercus lobata*, revealing features consistent with oak evolutionary success. Effective population size remained large throughout history despite declining since the early Miocene. Analysis of 39,373 mapped protein-coding genes outlined copious duplications consistent with genetic and phenotypic diversity, both by retention of genes created during the ancient *γ* whole genome hexaploid duplication event and by tandem duplication within families, including the numerous resistance genes and also unexpected candidate genes for an incompatibility system involving multiple non-self-recognition genes. An additional surprising finding is that subcontext-specific patterns of DNA methylation associated with transposable elements reveal broadly-distributed heterochromatin in intergenic regions, similar to grasses (another highly successful taxon). Collectively, these features promote genetic and phenotypic variation that would facilitate adaptability to changing environments.

## Introduction

Oaks are a speciose tree genus of the temperate forests of the northern hemisphere (from Canada to Mexico in North America, Norway to Spain in Europe, and China to Borneo in Asia) ^1,2^. The genus evolved in the palearctic during a time when the earth experienced a warmer climate ^3^. Fossil records indicate that sections within the genus — *Quercus, Lobatae*, and *Protobalanus* — were already present in the arctic during the middle Eocene 47.8–38 Mya ^3^. As the planet cooled, oaks disappeared from the arctic and migrated southward, speciating as they spread over Asia, North America, and Europe. Throughout these regions, the resultant species were the foundational constituents of their plant communities ^3^. This genus, which has diversified into two subgenera, eight sections, and more than 400 species ^4^, is an “evolutionary success story” ^1^. In North America, oaks have more biomass than any other woody plant genus, including pines ^5^, making this genus an ecological success story as well. As dominant species, oaks play pivotal roles in shaping biodiversity, creating healthy ecosystems, and sequestering carbon needed to mitigate climate warming. Throughout human history, they have provided valuable food, housing, materials, and cultural resources across multiple continents. Here we seek insights from the oak genome to uncover mechanisms that underlie the success of oaks.

We report details of a high-quality annotated chromosome-level genome assembly for *Quercus lobata* Née (valley oak; tree SW786) and associated tissue-specific methylomes. We analyze sequence trends of heterozygosity in valley oak and the European pedunculate oak (*Q. robur*) to show that effective population size (*N_e_*) has declined over time, but remained sufficiently large since divergence from a common ancestor to retain high levels of genetic variation. Large *N_e_* could help response to selection as the environment has changed over the last 50 million years. Further, our analysis of tandemly duplicated genes identifies large numbers of duplicated families, which, as Plomion et al. ^6^ also report, are particularly enriched for resistance genes and are likely associated with longevity and the eternal “arms race” with pests. We discover a large tandemly duplicated gene family that may be part of a previously undescribed non-self-recognition system that could prevent self-fertilization and promote outcrossing, or selectively allow occasional hybridizations. We also find many genes retained from the ancient *γ* paleohexaploid duplication event of the core eudicots. These are enriched for transcription factors and housekeeping genes, which may be more subject to strong (hard) selective sweeps than the tandemly duplicated genes^7^. Finally, we find some surprising similarities with the genomes of Poaceae (grasses — also highly successful plants). DNA methylation (BS-Seq) patterns indicate heterochromatin-rich chromosome arms, and additionally show CHH methylation peaks upstream of transcription start sites. Such prominent “mCHH islands” are known in maize ^8^ and a few other plants. These features could both affect gene expression and also facilitate tandem duplication events creating phenotypic variation and opportunities for selection.

## Results

### Genome assembly

An initial draft genome (version 1.0)^9^ was assembled from small (≈150x coverage) and large insert (≈50x) Illumina paired end reads. The final assembly (version 3.0) was constructed with the addition of Pacific Biosciences long reads (≈80x) and Hi-C long-range links produced by Dovetail Genomics and the HiRise re-scaffolder^10^, dramatically increasing NG50 scaffold size from 2 kbp to 75 Mbp (see **Methods**). The twelve longest scaffolds (“chromosomes”) were named and oriented to agree with pedunculate oak *Q. robur* ^6^, and correspond (in order, but not generally orientation) with the twelve linkage groups (LGs) of an existing moderate-density physical map of *Q. robur* × *Q. petraea* ^11^ that we did not use during sequence construction. This physical map consists of 4,217 sequence context-defined single nucleotide polymorphism (SNP) markers (after dropping 22 SNPs associated to two physical locations each). The LGs and our chromosomes show a predominantly monotonic one-to-one correspondence (e.g., chr. 1 in Figure 1A; see details in SI Section II: Validation and orientation of chromosomes, Figures S1 and S2, and Table S2). 99% of SNPs had at least one BLASTN alignment to our genome, and 98% of these had at least one alignment to the same chromosome as its LG. (95% of SNPs had all alignments to the same chromosome, and 86% had a unique alignment.) A small stretch of our chromosome 1 was found to be a mis-assembled mitochondrial sequence and was replaced by a gap of the same length (Figure S3).

**Figure 1:**
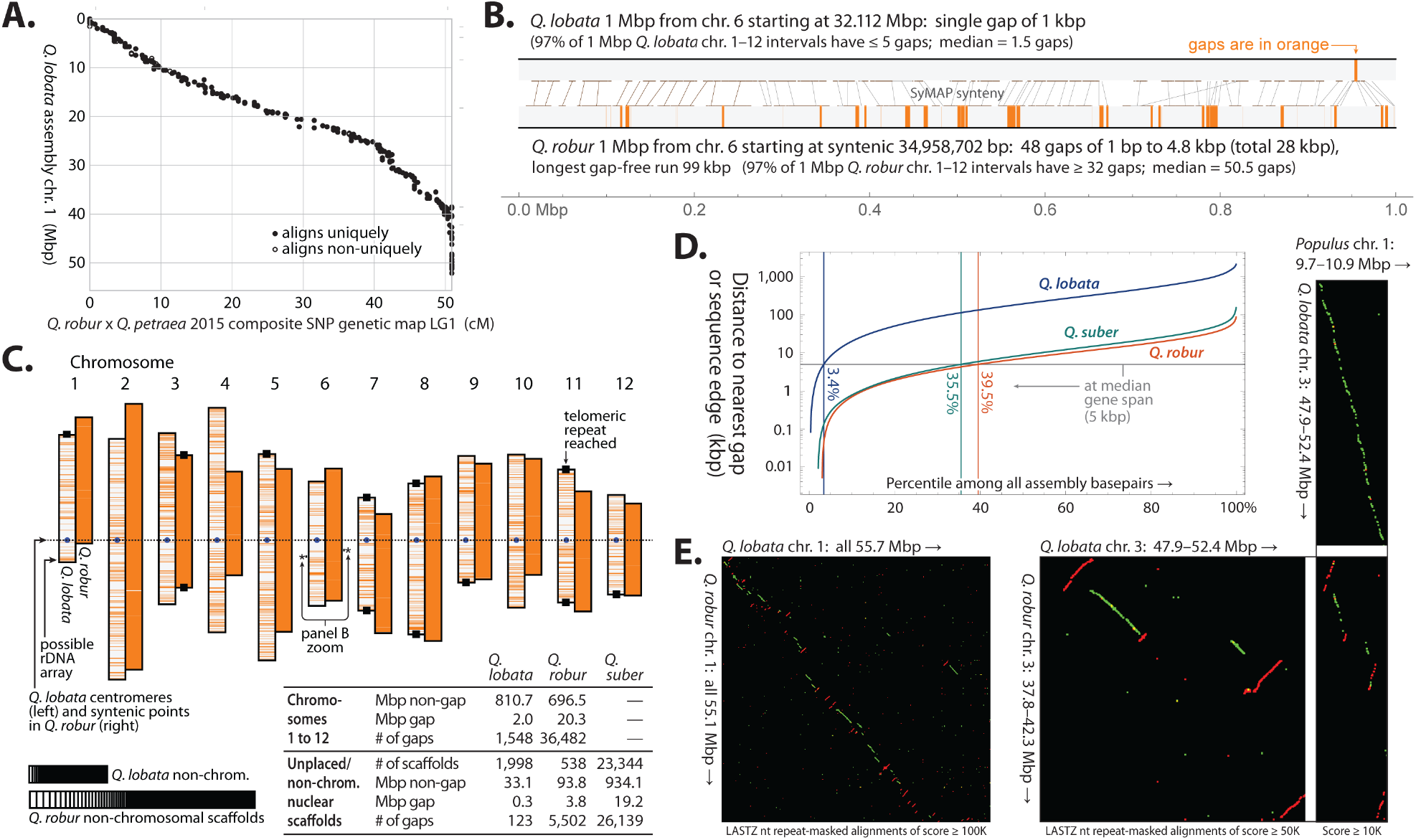
Overview of assemblies of *Q. lobata* tree SW786 (version 3.0), *Q. robur* (version PM1N) ^6^, and *Q. suber* (version 1.0) ^12^. (**A)** Alignment of a physical map linkage group 1 to *Q. lobata* chr. 1, exhibiting high concordance and overall monotonicity. (**B)** A representative 1 Mbp region from the *Q. lobata* assembly (top) and the syntenic 1 Mbp region from the *Q. robur* assembly (bottom), showing gaps in orange. (**C)** Overview of the chromosome-level assemblies (*Q. lobata* left member of each pair, *Q. robur* right) with orange lines indicating gaps, and basic statistics for all three assemblies. (**D)** Distributions of distance from a random base pair to the nearest gap or sequence edge. **(E)** Nucleotide alignments of entire chr. 1 of *Q. lobata* and *Q. robur*, showing numerous apparent rearrangements and inversions, in contrast to a more detailed illustrative region between chr. 3 of the two *Quercus* with chr. 1 of more distant *Populus trichocarpa* ^14^, in which the *Q. lobata* assembly is straight-line syntenic with *Populus* but that of *Q. robur* is not. Alignments between nominal same/opposite strands are colored green/red.

A comparison of our assembly with the two others available for *Quercus* — a chromosomal-level one for *Q. robur* ^6^ and a short-scaffold one for cork oak *Q. suber* ^12^ — revealed high similarity, despite ≈35M years since a common ancestor ^13^. This similarity is both at the level of repeats (see **Repetitive sequences**), as well as non-repetitive non-gap sequence where LASTZ aligns 88% of *Q. lobata* to *Q. robur* with average nucleotide identity 96%, and 86% of *Q. lobata* to *Q. suber* with average 93% identity. The larger contributing alignments tend to have even higher identity; e.g., the longest alignments capturing half of *Q. lobata* have average identity 98% for *Q. robur* and 95% for *Q. suber*.

Our assembly is characterized by much higher contiguity than the other two. For example, comparing *Q. lobata* vs. *Q. robur* and *Q. suber*, the number of gapless runs (“contigs”) is more than an order of magnitude smaller at 3.7k vs. 43k and 49k, respectively; N50 for gapless runs is more than 20-fold larger at 966 kbp vs. 37 kbp and 45 kbp; and N90 is also more than 20-fold larger at 205 kbp vs. 10 kbp and 9 kbp. Comparing a representative 1 Mbp from *Q. lobata* and the syntenic 1 Mbp from *Q. robur* (Figure 1B), the former has a single gap of 1 kbp while the latter has 28 kbp in 48 gaps of 1 bp to 5 kbp each. This pattern is typical: over all 1 Mbp regions from chr. 1–12, *Q. lobata* has median 1–2 gaps (97% of regions have ≤ 5), but *Q. robur* has 50–51 (97% having ≥ 32). Our assembly reaches telomeric repeats on both ends of four chromosomes, and on one end of four more. (Telomeric repeats, centromeres, and rDNA are discussed in SI Section V: Repetitive sequences.) Visualizing the entire *Q. lobata* and *Q. robur* assemblies (Figure 1C), *Q. robur* gaps appear nearly solid. The percent of non-gap sequence placed in a chromosome is 96% in valley oak vs. 88% in pedunculate oak (and 0% in cork oak). The three assemblies differ considerably in total Mbp of non-gap sequence: 845 Mbp for valley oak vs. 791 Mbp and 934 Mbp for pedunculate and cork oak, respectively. However, there are three *Q. suber* scaffold populations by length, and the longest — those ≥ ≈50 Kbp — total a more comparable 837 Mbp. More than a third of *Q. robur* and *Q. suber* base pairs are closer than a median gene span (5 kbp) to an assembly gap or sequence edge, while 96% of *Q. lobata* base pairs are further away (Figure 1D).

Apparent segmental rearrangements and inversions between *Q. lobata* and *Q. robur* were unexpectedly prevalent (e.g., Figure 1E left shows chr. 1 vs. chr. 1 as typical). Most of these, however, are likely scaffolding errors in *Q. robur*. Pedunculate oak has much smaller contigs, and its scaffolding was constructed using linkage maps (which are low in resolution compared to Hi-C) as well as synteny to *Prunus*, which may lead to mistakes in order and orientation of contigs (especially for small contigs). By contrast, alignments of *Q. lobata* with more distant species (*Populus, Eucalyptus, Theobroma*, and *Coffea*) showed numerous and widespread regions in continuous syntenies where *Q. robur* was not as continuous; to illustrate, Figure 1E right shows Mbp-scale regions of chr. 3 of the two *Quercus* vs. chr. 1 of *Populus*. Further, comparison of the formerly mentioned LGs from *Q. robur* × *Q. petraea* to both the *Q. lobata* and *Q. robur* assemblies shows *Q. robur* with more disagreements (Figures S1, S2). Thus, with the currently available *Q. robur* assembly, we conclude that alignments of *Q. robur* versus, e.g., *Q. lobata* are not reflective of true rearrangements and inversions.

#### Demographic histories of Q. lobata and Q. robur

Ancient oaks evolved over 50 Mya, initially in the subtropical climate of the palearctic of the Northern Hemisphere and, as the planet cooled, shifting southward to their contemporary distribution throughout the Northern Hemisphere (Figure 2A). Consistent with the large range, we found heterozygosity (average 0.50%–0.66%; see SI Section III: Analysis of heterozygosity, and Figures S4, S5) across the genome to be similar to but slightly less than the 0.73% computed for *Q. robur*, possibly due to the much larger species range of pedunculate oak and/or lower representation in the *Q. robur* assembly of highly homologous sequence loci resulting in increased post-alignment pileup of multiple actual loci at single assembly loci. To gain insight into the population history of oaks, we inferred the effective population size (*N_e_*) of *Q. lobata* and *Q. robur* over time. The Pairwise Sequentially Markovian Coalescent (PSMC’) method ^15^ applied to the individuals used to build the genomes mapped to their own assemblies (Figure 2B) enabled examination of the last ≈25M years of evolution (Figure S7). To verify accurate inference on this timespan, we generated simulated datasets using the inferred demographic history. We selected ancestral population sizes matching empirical genome-wide heterozygosities (see **Methods** and SI Section IV: Demographic analysis, and Figures S8, S9, and S10), and display these in Figure 2C.

**Figure 2:**
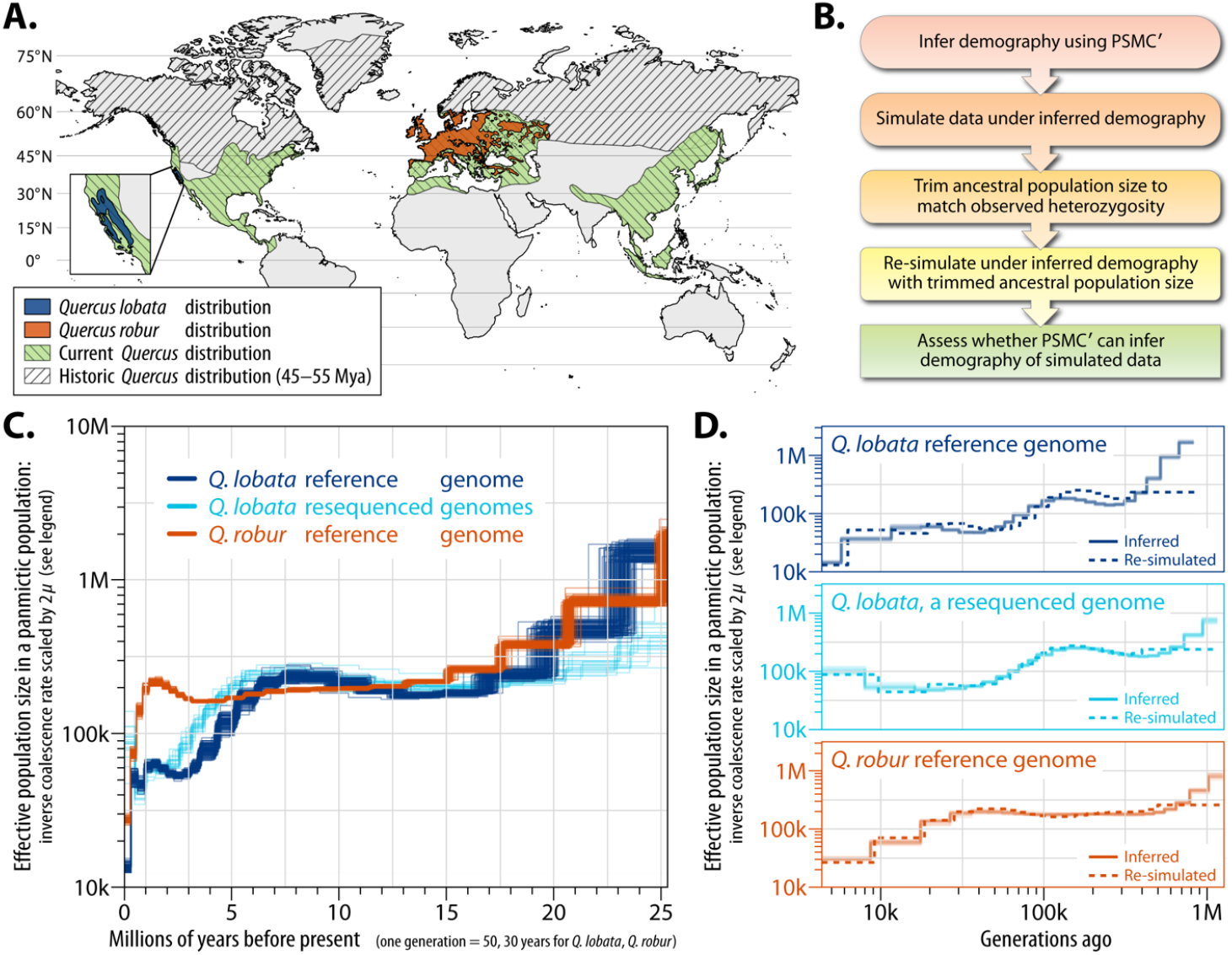
Demographic evolutionary history analysis of *Q. lobata* and *Q. robur*. (**A)** Historic and contemporary species ranges based on Barrón, et al. ^3^ and fossil occurrence records from the Global Biodiversity Information Facility website (GBIF.org 19th January 2019, https://doi.org/10.15468/dl.kotc15). Contemporary distribution of *Q. robur* is from the European Forest Genetic Resources Programme (http://www.euforgen.org/species/quercus-robur/); *Q. lobata* is based on Griffin and Critchfield ^21^. **(B)** Stages of the analysis. **(C)** Inferred effective population sizes over time via PSMC’ (100 bootstraps shown per condition), using a mutation rate of 1.01 × 10^−8^ bp per generation (see SI Section IV: Demographic analysis and Figure S6 for other parameters). **(D)** PSMC’ accurately infers demography (solid) on data simulated (dashed) under models fit to the empirical data.

We ran PSMC’ on data simulated under trimmed demographic models and found accurate inference of population size over time, except for the single oldest time step where population sizes were often over-estimated (Figure 2D). The PSMC’ analysis indicates ancestral populations of both *Q. lobata* and *Q. robur* had high (>500k) effective population sizes that then showed initially similar declines, perhaps as populations were shifting southward (Figure 2C). *Q. lobata* shows an additional decline in *N_e_* at ≈5 Mya, which would have occurred after the shift from a period of subtropical climate with year-round rainfall to a Mediterranean climate with summer drought ^16^. By contrast, for *Q. robur* (being more widely distributed throughout Europe), *N_e_* remained relatively flat until the last ≈1M years. At this point, *Q. robur* declines to *N_e_* < 50k (and below *Q. lobata*) during the “Ice Ages” when the region was experiencing a series of warm and cold periods creating genetic bottlenecks and expansions (Figure 2C and D). Both species have retained sufficiently large effective population sizes to facilitate natural selection ^17^.

#### Repetitive sequences

As with many plant species, the valley oak genome contains substantial repeats, with 54% of non-gap base pairs marked as repetitive by RepeatMasker in combination with a species-specific database constructed by RepeatModeler+Classifier (Figure 3A and B; the modeling step was essential, as RepBase only marked 13%). The largest identified portion is transposable elements (TEs), primarily Copia and Gypsy elements of the long terminal repeat (LTR) type. The level of repetitiveness is similar to the 54% (disregarding gaps) found by application of the same process to *Q. robur* (for which Plomion et al. ^6^ reported 52% via REPET and other annotation, including manual curation). RepeatModeler+Classifier also detects 51% in *Q. suber* ^12^, 55% in *Eucalyptus* ^18^, 55% in *Theobroma* ^19^, 51% in *Coffea* ^20^ and 43% in *Populus* ^14^. Centromeric, telomeric, and rDNA repeats for valley oak were identified (see SI Section V: Repetitive sequences), and specific sequence-defined repeat superfamilies are correlated or anticorrelated to various levels with centromeric proximity, forming (as do protein-coding gene exons) density gradients that are the main chromosome-scale repeat-associated features, presumably reflecting overall chromatin structure (Figures S11, S12, and Figure 3C–D).

**Figure 3:**
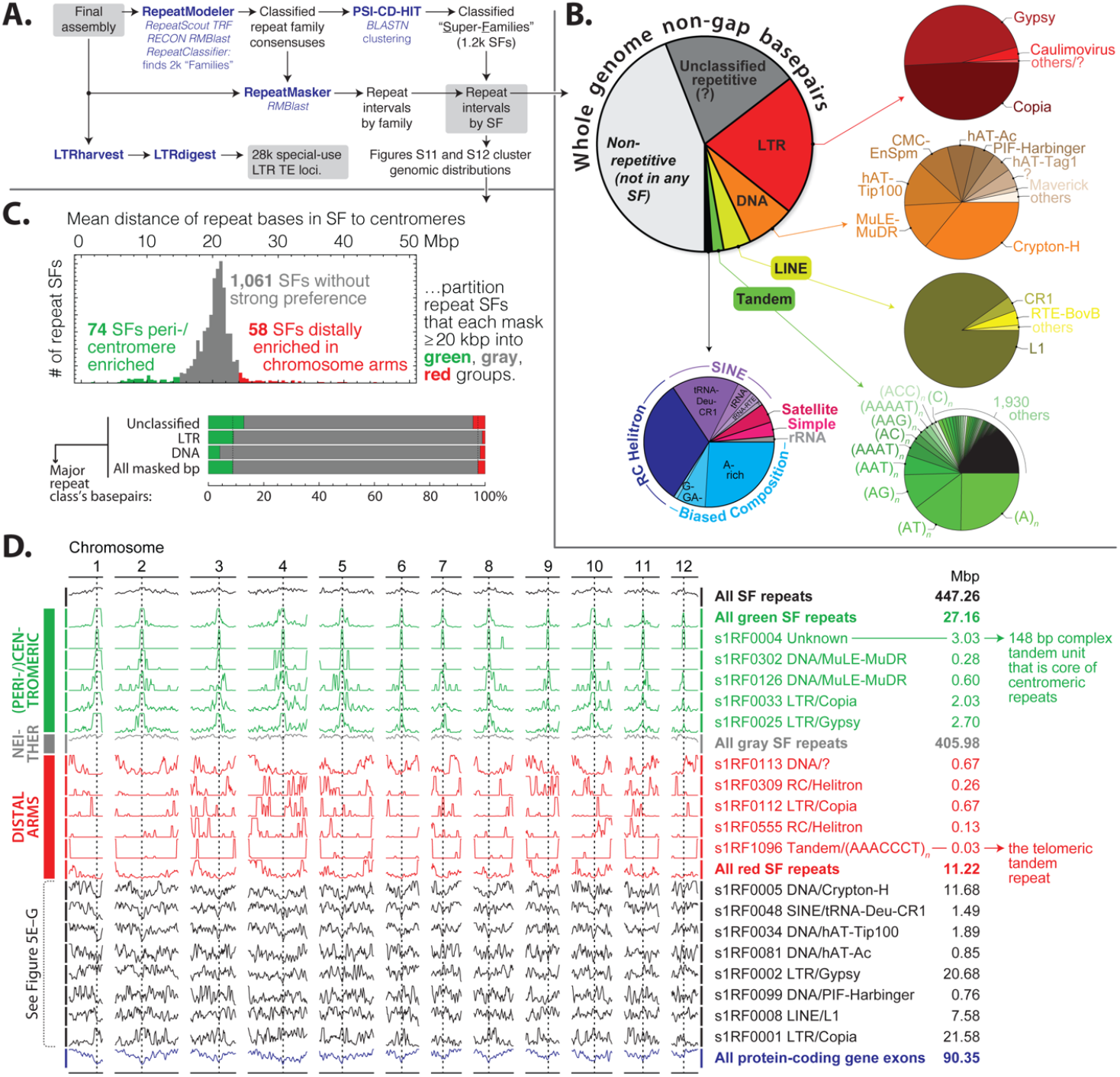
Dispersed and local (tandem/satellite, simple/biased composition) repetitive sequence in *Q. lobata*. **(A)** Primary analysis outline. **(B)** Assembly partitioned into RepeatClassifier/RepeatMasker major and minor classes; 54% of non-gap base pairs are covered by repeat superfamilies (SFs), and transposable elements (TEs) are prevalent. **(C)** Unsupervised comparisons of how the 1,193 individual SFs each with ≥ 20 kbp distribute across chromosomes (Figures S11 and S12) suggest the primary distributional diversity at chromosome scale is proximity to centromeres (green, 74 SFs totaling 27 Mbp) vs. telomeres (red, 58 SFs totaling 11 Mbp) vs. more-or-less uniformity (gray, 1,061 SFs totaling 406 Mbp). **(D)** Chromosomal distribution of selected SFs and sets of SFs, illustrating the diversity across and within the trichotomy of (C). The *y*-axis in each row is linear number of member base pairs in 3 Mbp bins every 1 Mbp, with zero at the lower edge and 95th percentile (or row maximum if the percentile is zero) at the upper edge. Black rows near the bottom are the representative SFs of Figure 5E–G.

The repeat content of *Q. lobata, Q. robur*, and *Q. suber* are very similar at the sequence level. There are six combinations in which RepeatModeler can be used to build a species-specific repeat consensus database from one of the three *Quercus* assemblies, which can then be applied by RepeatMasker to one of the two other assemblies. In all six combinations, 89% to 92% of non-gap base pairs are marked the same way (repetitive or not repetitive) as when the native consensus database for the species being masked is used.

#### Gene prediction and annotation

Using the AUGUSTUS gene modeler ^22^ and a diverse set of experimental data (Iso-Seq, RNA-Seq, DNA methylation) and *in silico* data (known proteins, repeats), we modeled 68k putative protein-coding genes (PCGs) (see **Methods**, Figure 7A and Table S3). As many corresponded to transposons with little expression or appeared hypothetical for other reasons, we removed 29k to obtain the primary set of 39,373 PCGs we report, of which 35k have at least one intron and all of which have UTRs annotated and are ostensibly complete. *Q. robur* reports only 29k PCG models, of which just 20k have introns, and about half UTRs; in the other direction, *Q. suber*’s annotation by NCBI (thinned to one isoform per locus) reports more 49k PCG loci (about half with UTRs), but a more comparable 36k with introns and 38k ostensibly complete, and with a much higher number containing transposon domains by comparison.

**Figure 4:**
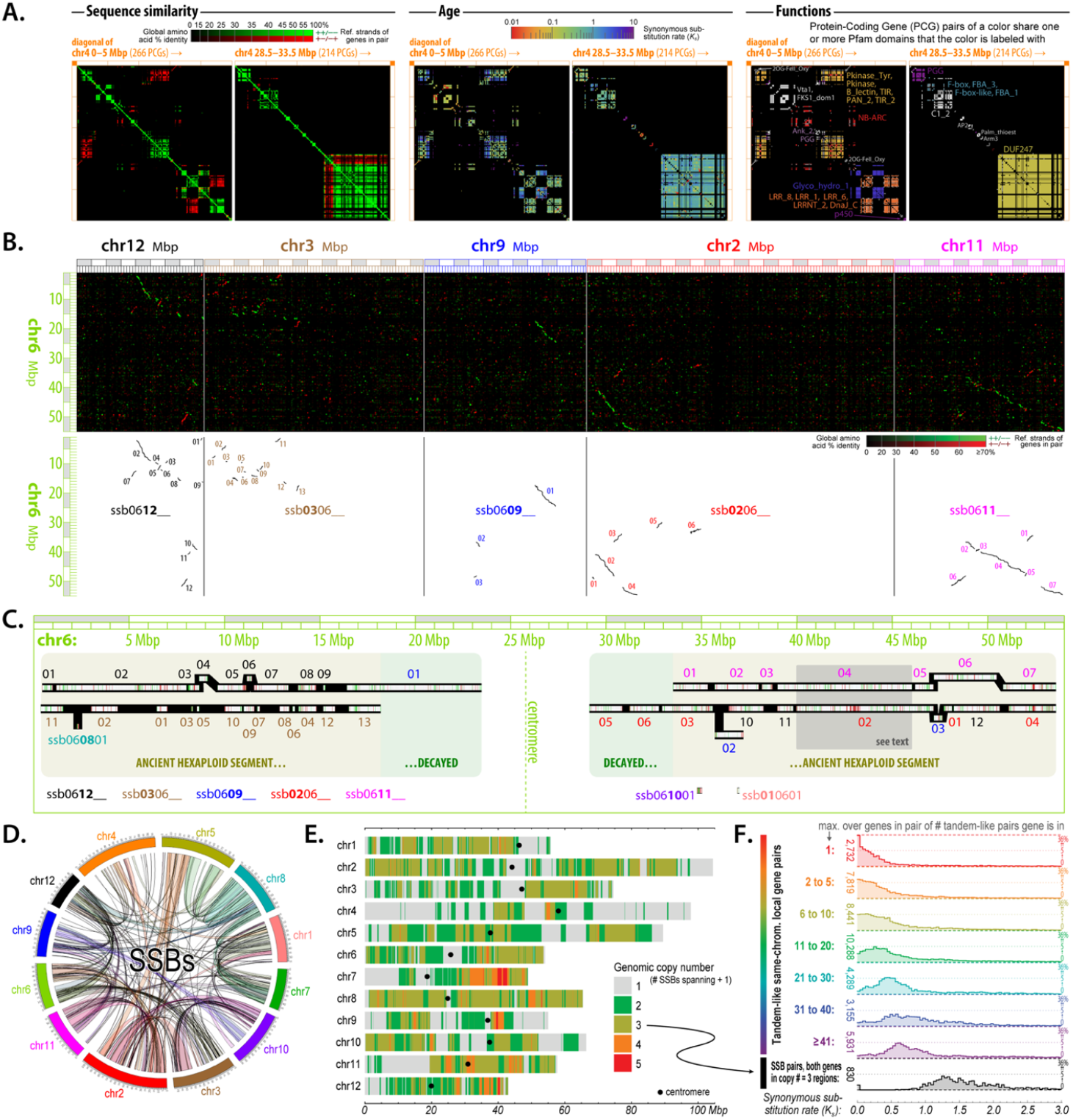
Duplicated protein-coding genes. **(A)** Sequence similarity (amino acid identity), age (synonymous substitution rate *K_s_*), and functions (shared Pfam domains) for all pairs of proteins within two illustrative 5 Mbp regions of chr. 4. Nearly half of *Q. lobata* PCGs are involved in tandem-like blocks of varying sizes (up to Mbp scales and dozens of genes at a time), often locally rearranged, and originating and growing at a variety of ages. Genes involved are diverse, but enriched in certain functions. **(B, C)** With no recent whole-genome polyploidization, most of the detected PCG syntenies of *Q. lobata* to itself (“SSBs”) are small and diffuse and reflect the core eudicot triplication event *γ* over 100 Mya. Despite its age, this event remains quite evident — albeit highly fragmented, dispersed, and partially decayed. The whole of chr. 6 vs. the whole of chr. 12/3/9/2/11 are shown as exemplary. **(D, E)** SSBs [even without chaining as in (C)] cover much of the chromosomes. The highest fraction (34% of base pairs) is spanned by manifest triplication, 27% by duplication (while some duplication is recent, most appears to be decayed triplication), and 34% by no detected extant synteny. **(F)** The pairwise synonymous substitution rate (*K_s_*) tends to be very low for-genes tandemly duplicated just once (red) and increases as tandem-like block size increases (orange to violet), suggesting larger blocks are older. *K_s_* is essentially always extremely high (≥ ~1.0) for SSB gene pairs where both pair-genes lie in chromosomal regions spanned by exactly two SSBs (black), supporting the syntenic triplications to be of ancient origin.

**Figure 5:**
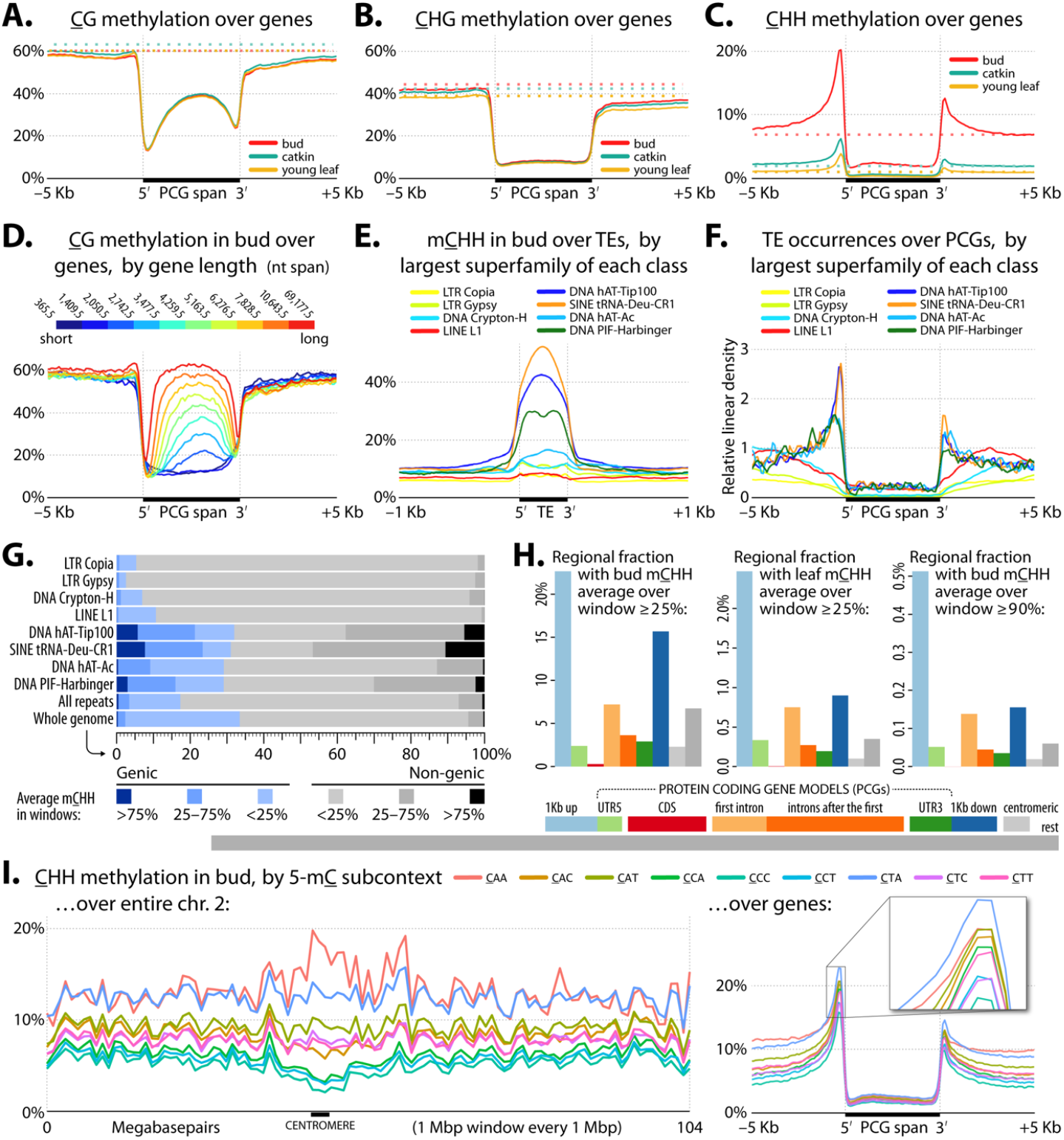
*Q. lobata* DNA methylation in protein-coding genes and repeats. **(A–C)** Average methylation levels (100 bp windows) with respect to PCGs (normalized to 5 kbp long) for the three sampled tissues (bud, catkin, and young leaf) by methylation context: **(A)** CG, **(B)** CHG, and **(C)** CHH. Dotted lines show genome-wide backgrounds, and TSS/TES = Transcription Start/End Site. **(D)** mCG for-genes in deciles by gene length. **(E)** Average bud mCHH (20 bp windows) across representative repeat SFs (normalized to 400 bp long) in selected RepeatClassifier minor classes. **(F)** Relative density of representative repeat SFs around genes (100 bp windows). **(G)** Distribution of mCHH for representative repeat SFs (100 bp sliding disjoint windows). ‘Genic’ = gene spans enlarged by 1 kbp on each end. **(H)** Partitioning of whole genome base pairs into nine types of regions vs. mCHH coverage. Lower horizontal bars reflect relative size. Vertical bars show percent of each genomic context covered by 100 bp windows with mCHH > 25% or > 90% in bud or young leaf. **(I)** mCHH by 3 nt subcontext (queried cytosine is underlined, and is the first nt of the three); l**eft side:** 1 Mbp windows across all of chr. 2, **right side:** across genes (normalized to 5 kbp length) in bud tissue.

**Figure 6:**
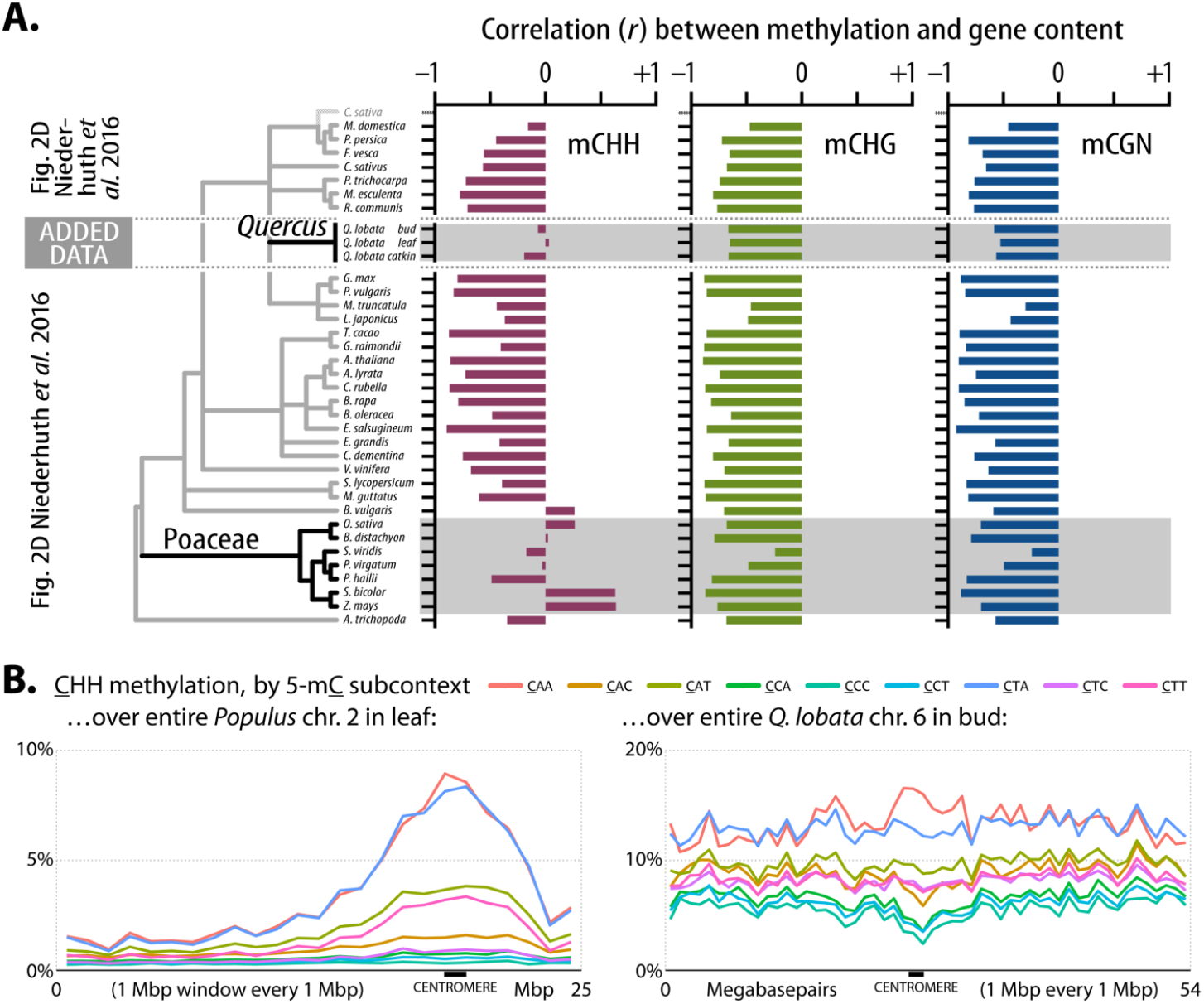
(**A)** Pearson correlation (*r*) between methylation level and number of genes (100 kbp windows) for mCHH (left), mCHG (middle), and mCG (right) context levels from leaf tissue-based Figure 2D of Niederhuth et al. ^43^, inserting our values for three oak tissues (bud, leaf, and catkin from tree SW786, having matched analysis details as closely as possible). **(B**) Comparison of all nine DNA subcontext methylation levels within the CHH context over an illustrative chromosome of *Popular trichocarpa* ^46^ and *Q. lobata*. (See Figure 5 legend.)

**Figure 7:**
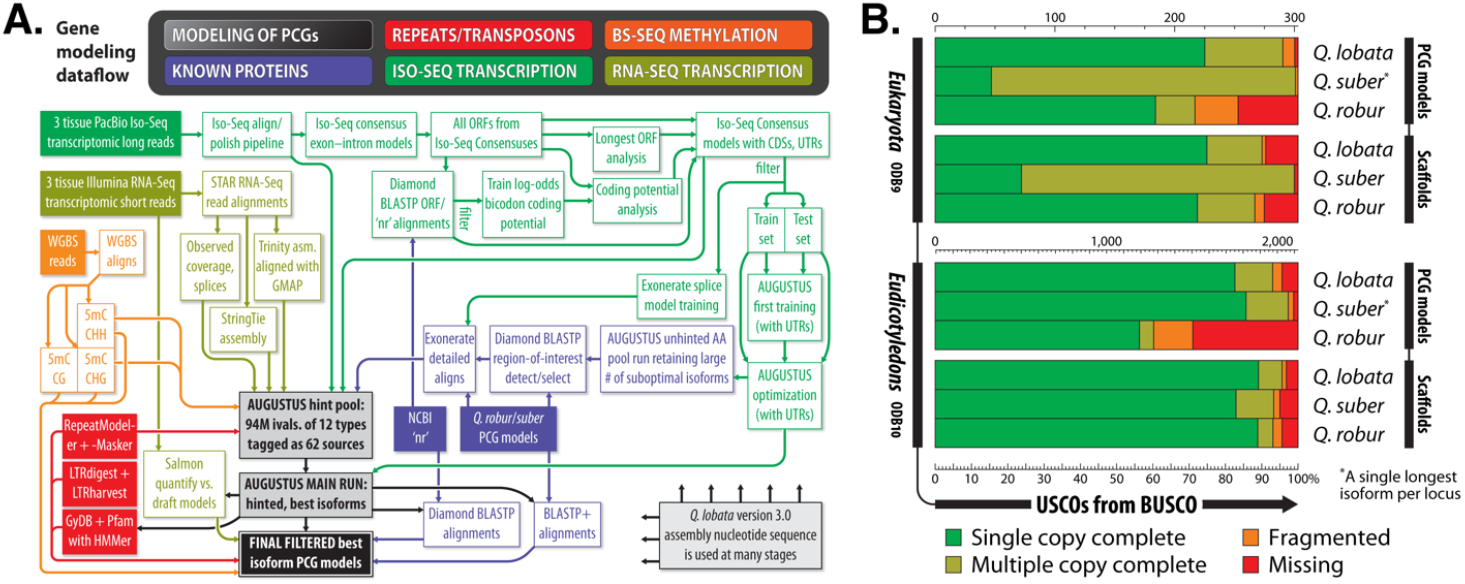
**(A)** Dataflow of protein-coding gene (PCG) modeling. **(B)** BUSCO v3 analysis of PCG models and genomic scaffolds for *Q. lobata, Q. robur*, and *Q. suber* against the ODB9 Eukaryota and ODB10 Eudicotyledons USCO sets.

We assigned gene names, functions, and orthologs via the PANTHER and Pfam components of InterProScan, and OMA ^23^. We evaluated the *Q. lobata, Q. robur*, and *Q. suber* scaffolds and single isoform PCG model sets with BUSCO (Figure 7B). *Q. lobata* compares favorably to the other two, and does not have the high multicopy anomaly of *Q. suber* in the 303-USCO ODB9 Eukaryota set ^24^, or the high missing and fragmented fraction of *Q. robur*’s small model set (especially with the more comprehensive 2,121-USCO set for Eudicotyledons from ODB10).

#### Gene duplications

Protein–protein alignments among the *Q. lobata* PCGs exposed a rich panoply of duplication structure in terms of genomic positions, ages, and functions. Prominent and complex tandem-like blocks of high-similarity genes can be seen via visualizations of all– vs.–all alignments (see **Methods**). These duplications often involve local rearrangements, and can extend into megabases with dozens of genes involved at a time. Figure 4A (left third) exhibits two illustrative 5 Mbp regions of chr. 4. Approximately 40% of PCGs participate in these blocks, which have sizes of two to ≈100 genes each, with larger sizes rarified like a power law (Figure S13). Roughly a third of participating genes are duplicated only once, slightly more than half two to 20 times, and only a tenth more than 20 times. Visualizations (e.g., coordinated Figure 4A middle third) of the synonymous codon substitution rate (*K_s_*) over gene pairs in blocks suggest a wide variety of ages for the majority of retained expansion for individual blocks. Larger blocks tend to be older (Figure 4F colored distributions), but even old blocks tend to have younger points suggestive of ongoing growth. While numerous tandem gene copies are shorter or have reduced or no RNA-Seq evidence of expression, many copies (even within larger blocks) are not particularly short or of lower expression and so do not appear to be pseudogenes. Functions of tandemly duplicated genes are diverse, as evident from the variety of Pfam domains they contain (e.g., coordinated Figure 4A right third). Relatively few distinct domains, however, are strongly enriched over all tandemly duplicated genes, and include NB-ARC, LRR_8, B_lectin, LRR_1, TIR_2, LRRNT_2, p450, TIR, and PGG (associated with resistance/defense); Pkinase_Tyr and Pkinase (signal transduction); UDPGT (the large UDP-glucoronosyl/glucosyl transferase family); S_locus_glycop, PAN_2, and DUF247 (see below); F-box, FBA_3, and FBA_1 (protein–protein interactions/degradation, signal transduction and regulation); and GST_N, GST_N_3, and GST_N_2 (glutathione S-transferases, with functions including stress tolerance/signaling and detoxification).

Many of the strongly enriched domains are part of the domain architecture of plant disease resistance genes (R-genes) identified by Gururani, et al. ^25^. It is difficult to be sure whether an R-5gene is actually acting as a pathogen defense mechanism in a given plant species, but Gururani, et al.^25^ reviewed the experimenal evidence and identified eight classes of R-genes based on the arrangements of domains and structural motifs. Using their criteria for domain combinations (see Supplementary Information, Section VI. R-gene identification), we analyzed the domain architecture of 39373 *Q. lobata* proteins and found 751 R-genes, which contained the highly likely combination of domains involved in R-genes, and another 2176 genes that are good candidates for R-genes because they include enough qualifying features (Table S4). For the *Q. robur* annotation (25,808 proteins) ^6^, we counted 632 R-genes plus 1645 candidate R-genes and for *Q. suber* (49,388 proteins) ^12^, we found 723 *R-*genes plus 2182 candidate R-genes (Table S4). These numbers are based on the predicted gene models from each genome rather than DNA sequences, and the differences among species are more likely to represent differences across annotations rather than differences in DNA sequences. Collectively they document high levels of R-genes in oaks and illustrate tremendous opportunity for plant defense mechanisms.

### Possible DUF247-based non-self-recognition system

An investigation of PCGs found in blocks containing at least 30 tandemly duplicated genes uncovered DUF247 (PF03140) as the most enriched Pfam domain (Table S5; also see large block in Figure 4A). The only known suggested function for DUF247-containing genes (“DUF247 genes”) is from the Poaceae family, where two DUF247 genes in rye grass segregate with each of two known self-recognition loci and are proposed to be the male determinants of a multi-locus self-incompatibility system^26,27^. Among the evidence is a self-compatible rye grass species with a disrupted DUF247 gene ^26^. If the DUF247 gene family affects self-recognition in oaks, the extensive duplication suggests a non-self rather than a self-recognition system ^28,29^. This type of system has been demonstrated in Solanaceae, e.g., petunia ^30^ and tomato ^31^, Rosaceae, e.g., pear and apple ^32^, and Plantaginaceae, e.g., snapdragon ^33^. In these, the S-locus includes a single female determinant gene (S-RNAse) and commonly seven to 20 linked paralogs of male determinant F-box genes (SLFs). In snapdragon ^33^, up to 37 linked male determinant SLF genes were observed, while (at the other extreme) *Prunus* species have a single F-box gene for the male determinant and appear to have adopted an S-RNAse-based self-recognition system rather than non-self-recognition ^34^, demonstrating the feasibility of transitioning from one to the other. The span of the large DUF247 block (Figure 4A) contains 34 predicted PCGs with a complete DUF247 domain, 22 with partial DUF247 domains, and 17 additional genes. Among the 17 additional genes are two pectinesterase inhibitor-like genes shown to be involved in regulating pollen tube growth in maize ^35^, a pectin depolymerase gene, two E3 ubiquitin ligases that have been shown to confer self-incompatibility when transplanted to *Arabidopsis* ^36^, a DNA helicase, and 11 uncharacterized genes. DUF247 genes are entirely specific to plants and usually carry a single copy of the domain that comprises almost the entire gene. Across the 104 plant genomes in Pfam Release 33.1^37^ with ≥ 17,500 predicted protein entries (to restrict to genomes most likely to be complete), the top five by number of DUF247 domain occurrences are three tree species — *Juglans regia* (English walnut) *n* = 201; *Eucalyptus grandis*, *n* = 188; and *Populus trichocarpa* (black cottonwood), *n* = 161 — and two polyploid cultivars (wheat, *n* = 192, and peanut, *n* = 165). These tree species, like *Q. lobata* (*n* = 186), do not have identified incompatibility systems, are frequently highly outcrossing, and sometimes self-fertilize at low levels.

#### Long-surviving duplicated genes

Also striking in the visualizations of protein alignments were self-syntenic blocks (SSBs): syntenic runs of proteins within *Q. lobata*, generally between different chromosomes, with a variety of lengths and gene pair densities. Figure 4B (top) shows chr. 6 vs. chr. 12/3/9/2/11 as exemplary (although in low resolution per limited space). For further analysis, 236 SSB runs, each with four to hundreds of gene pairs, were extracted (e.g., Figure 4B bottom) and given accessions “ssbXXYYZZ” with XX ≤ YY indicating the chromosomes involved and ZZ as serial number; more than 7,100 PCGs are directly involved. High resolution examination made evident that, on any given chromosome, runs tended to end and begin close by, and for any particular point on a chromosome to be covered by very few runs (typically, zero to two), so that (nearly) disjoint SSBs could often be clearly ordered to form a small number of chromosome-scale chains (Figure 4C black bars). While a few recent segmental duplications appear, most SSBs are likely “ghosts” of the ancient genome triplication polyploidy event ɣ associated with early diversification of the core eudicots, thought to have occurred about 120 Mya ^38–40^. The high age of many SSBs is supported by the synonymous substitution rate (*K_s_*) for gene pairs in SSBs in triplicated regions being very high (almost entirely > 1.0; Figure 4F black distribution), as well as the generally short length and scattered nature of SSBs (which are within *Q. lobata*) compared to syntenies between *Q. lobata* and different species (*Populus*, *Eucalyptus*, *Theobroma*, and *Coffea*).

While general triplication is clear from the gene pair-defined SSBs (e.g., Figure 4C white bars, with green and red showing supporting gene pairs), few syntenic gene triples have been retained, and detection and characterization of the ɣ ghosts would be unrepresentative for an analysis restricted to gene triples. For example, in the gray shaded region of Figure 4C involving chr. 6/2/11 and spanning 320 chr. 6 genes, the 59 chr. 6 genes supporting local one-to-one chr. 6/2 synteny have only eleven chr. 6 genes in common with the 39 supporting local one-to-one chr. 6/11 synteny. Even before chaining as in Figure 4C, two thirds of the genome are in a SSB (Figure 4D and E), with the largest fraction (34%) actually covered by two (consistent with triplication) and 27% by one (decayed triplications and a few recent segmental duplications); a third (34%) is not covered, and only 5% is covered by three or four SSBs (likely duplications post triplication). Relative to all genes, those in one or two gene pairs supporting SSBs tend to be of higher expression with lower repetitive sequence in their immediate vicinity, and are enriched for certain functional classes, including transcription factors and housekeeping genes (Table S6).

### Genome-wide patterns of DNA methylation and strong mCHH islands

Whole-genome bisulfite sequencing for bud, catkin, and leaf tissue revealed mean DNA 5-methylcytosine methylation (BS-Seq) levels in CG (mCG) and CHG (mCHG) nucleotide contexts as relatively stable across tissues (Figure 5A and B), while levels in CHH (mCHH; Figure 5C) were notably higher in bud than catkin and young leaf, likely due to the increased proportion of undifferentiated meristem tissue ^41^. Mean levels for regions surrounding genes are similar to genome-wide means for all tissues in all contexts (mCHH 1–7%, mCHG 39–43%, mCG 60–62%; Figure S14), with the exception of peaks of mCHH near transcription boundaries of genes (Figure 5C). These mCHH peaks are similar in both position and scale above background to the “mCHH islands” of maize ^42,43^ (Auxiliary Spreadsheet 1). We examined mCHH across representative repeat superfamilies (SFs), specifically, those of highest mass within selected RepeatClassifier minor repeat classes, as seen in Figure 5E. Within genic regions, three SFs — s1RF0048 (“SINE tRNA-Deu-CR1”), s1RF0034 (“DNA transposon hAT-Tip”), and s1RF0099 (“DNA transposon PIF-Harbinger”) — were both high in mCHH and preferentially located in the highly methylated gene boundary regions (Figures 5E and F). Members of these SFs are found in both genic and non-genic regions with broadly similar mCHH levels (Figure 5G and Figure S15). However, in view of overall genome-wide mCHH levels (including centromeres and intergenic space), we find regions surrounding genes to be highly enriched for mCHH (Figure 5H). Similar enrichment patterns are seen in bud and leaf, despite different overall mCHH levels (Figure 5C and H), and similar patterns are also seen if mCHH window stringency is varied from 25% to 90%, although at these extremes we observe decreases in the relative amount of downstream and non-genic mCHH (Figure 5H). All methylation is typically low near transcription boundaries (Figure 5A), and remains low for mCHG and mCHH across gene bodies. However, gene body mCG rises for longer-genes, reaching near-background levels in the longest genes (Figure 5D).

### Broad distribution of heterochromatin

*Q. lobata* appears to have heterochromatin dispersed throughout chromosomes more or less equally, with only minor increase of density toward centromeres. This interpretation is based on both the distribution of genes and repeats as well as indications of widespread histone-driven DNA methylation, a pattern more similar to maize and rice methylomes than to the *Arabidopsis* and tomato methylomes in which the methylated repeats are concentrated in pericentromeric heterochromatic regions ^44,45^. As such, a majority of repeat mass does not show strong positional correlation with centromeres (Figure 3D gray). Also, 92% of PCGs have a RepeatMasker-defined repeat within the gene’s upstream 2 kbp, which is high, because among 34 angiosperms, reported numbers range from 29% (*Arabidopsis*) to 94% (*Zea mays*), with an average of 50% ^43^. (See also Auxiliary Spreadsheet 1).

A second indication of heterochromatin-rich chromosome arms is the type of methylation found on intergenic repeats. Different mechanisms of generating plant mCHH, such as RNA-directed DNA Methylation (RdDM) or CMT3-mediated histone-associated methylation, have been shown to have distinct preferences for specific nucleotide subcontexts (finer than <u>C</u>G/<u>C</u>HG/<u>C</u>HH: <u>C</u>AA vs. <u>C</u>AC vs.…). Histone-associated mechanisms are typically responsible for methylation of heterochromatin and have much stronger biases than RdDM ^44^. *Q. lobata* has strong <u>C</u>HH subcontext preferences at chromosome scale (Figure 5I left and Figure S16). Bias patterns around centromeres are likely to indicate the general methylation pattern of heterochromatin in oaks, while chromosome arms represent a mix of genes and intergenic spaces. The peaks of m<u>C</u>HH surrounding gene boundaries (i.e., the m<u>C</u>HH islands) show a distinct pattern, with preference for <u>C</u>AA strongly reduced (Figure 5I right). Moving from gene boundaries toward intergenic space, the subcontext pattern progressively reverts to the likely heterochromatic signal of the centromeres (Figure S17).

An additional measure of the similarity in genome organization between oaks and grasses is the level of correlation between methylation and gene count across chromosomes. When we augment Figure 2D from Niederhuth et al. ^43^ with oak findings, oak is again found comparable to the grasses (Poaceae) and less to the other studied angiosperms (Figure 6A). The low mCHH and gene count correlation reflects a combination of unusually strong gene boundary mCHH islands relative to the background mCHH level (Figure S18 and Auxiliary Spreadsheet 1) and low average gene density in chromosome arms (Figure S19).

## Discussion

Our analysis of a high-contiguity, chromosome-level annotated oak genome reveals previously unreported features of oaks that might contribute to its ability for adaptation to new environments and resulting dominance in North American ecosystems. We find surprising similarities to grasses (Poaceae), another highly successful group of plants. Oaks and grasses both have genomes with large repeat-rich intergenic regions and share methylation features that are somewhat unusual, given the current sampling of methylomes in the literature. Interest has been growing in the adaptive potential provided by large complex intergenic regions often found in plants with larger genomes ^47–49^. For example, a substantially higher percentage of loci associated with phenotypic variation are found in the large intergenic regions of maize versus the smaller intergenic regions of *Arabidopsis* ^49^. Much of this regulatory variation has been found in non-TE stably unmethylated DNA ^50,51^, such that more than 40% of phenotypic variation in maize was associated with open chromatin that makes up less than 1% of the genome ^52^. On the other hand, high density of diverse TEs, which has been connected with local adaptation ^53^, can be a source of both transcription factor binding sites and regulatory non-coding RNAs ^54^, and play a role in three-dimensional genome structure ^51,55,56^. An abundance of intergenic heterochromatin-like structure has been demonstrated in grasses^8,57,58^ and, based on patterns suggestive of histone-driven methylation ^44,45^, are likely also found in oaks (Figure 5I, Figures S16, S17, S19 and Auxiliary Spreadsheet 1). Given the dramatic differences reflected in the chromosome-wide subcontext methylation patterns in the gene-dense arms of *Arabidopsis* and tomato versus the wider spread of genes in maize and rice^44^, and similar differences in poplar versus oak (Figure 6B and Figure S20), oaks and grasses may have some regulatory strategies distinct from those in other angiosperms. Another indicator of similarity between oaks and grasses is the correlation of CHH methylation levels (mCHH) and gene count along chromosomes (Figure 6A). A comprehensive characterization within oaks and across the angiosperms awaits further experimentation and better, more comparable genome sequences, constructed and annotated with consistent methods.

Pronounced mCHH islands are another feature shared between oaks and grasses. In maize, mCHH islands have been proposed to enforce boundaries between heterochromatin and euchromatin, and as such contribute to maintaining suppression of TEs during increases in neighboring gene expression ^8,42,59^. Measured as the ratio of peak mCHH to whole genome average mCHH, we find oaks have unusually strong 5’ mCHH islands (Auxiliary Spreadsheet 1), but it remains to be seen if they also contribute to boundary enforcement. It is possible they are simply the result of the type of TEs found near gene boundaries. In valley oak (Figure 5), maize ^60^, and *Arabidopsis* ^61^, mCHH is influenced by TE family, proximity to genes, and chromosomal location. The strong enrichment of small, highly methylated TE families near-genes (Figure 5E and F) could be due to, for example, selection against large TEs in gene proximal regions.

A potentially exciting discovery is the presence of many Pfam DUF247 domains in one of the largest and densest blocks of tandemly duplicated genes (Figure 4A), as these domains could be part of a non-self-recognition compatibility system ^29^. DUF247 genes have been implicated in a self-recognition system of ryegrass ^26,27^, analogous to S-RNAse-based self- and non-self-recognition systems in petunia ^30,31^, and tomato, apple, snapdragon, and peach ^62^. Oaks have long been thought to possess some kind of self-incompatibility system because of their high outcrossing rates, but the single gene SI systems have not fit observations. However, a non-self-recognition system would be consistent with observed crossing results among self, intra-, and inter-specific pollinations ^63^. Both self- and non-self-recognition systems of co-adapted genes expressed in pollen and pistil and preventing self-fertilization have evolved independently in several lineages of angiosperms ^29,64^. While the roles of DUF247-containing genes need experimental verification, their large numbers and high diversity at the amino acid level are consistent with a non-self-recognition system that could both promote outcrossing while also permitting occasional self and interspecific crosses.

Oaks have a vast reservoir of tandemly duplicated genes of a wide variety of ages (Figure 4F), contributing to their genetic and phenotypic diversity. As reported for pedunculate oak ^6^, resistance genes are a particularly prominent component of the tandemly duplicated gene blocks in valley oak, especially the larger (and older) ones (Auxiliary Spreadsheet 2: see worksheets for tandem pairs >20, 30, 40). The three oak genomes contain hundreds to thousands of potential R-genes: 732 to 2927 for *Q. lobata*, 632 – 2247 for *Q. robur*, and 793 to 2905 for *Q. suber*. In defending oaks from bacteria, viruses, nematodes, oomycetes, and insects, these R-genes may both enable the long lifespan of oaks^6^, and also address the puzzle of how a single or two oak species are able to dominate so many of the ecosystems they occupy. The classic Janzen–Connell ecological hypothesis proposes that pathogens promote tropical forest diversity through conspecific negative density-depending (CNDD) mortality, but CNDD has been shown across all forest types ^65,66^. In oaks, the high number and potential complexity of R-genes could provide a mechanism to reduce CNDD mortality caused by pathogens ^67^. Moreover, the large effective population size could maintain R-genes, especially if not costly ^68^. In fact, other ecosystem-dominant trees, which also contain large numbers of domains associated with resistance genes (such as, NB-ARC and LRRs), include the highly speciose *Eucalyptus* (~600 species) and *Populus* (Table S7). Extensive research demonstrating the importance of R-gene diversity at both the individual and the population level is ongoing in *Arabidopsis*, crop species and other plants ^69,70^. Studying oaks swith large and complex pools of R-genes will provide an important extension of this work.

Inspection of our highly contiguous genome identified numerous syntenic blocks of remnant genes from the g triplication event, which occurred ≈120 Mya ago when the common ancestor of angiosperms underwent two whole genome duplication events ^38–40^. More than 18% of protein-coding genes participate in a gene pair directly supporting a self-syntenic block (SSB), and more than a third of the genome is spanned by a manifest triplication (even without chaining blocks). SSBs (for example, Figure 4 and Auxiliary Spreadsheet 2) provide an extensive single genome resource for documenting remnants associated with the g event. Our annotation finds triplicate families to be enriched for transcription factors, as well as signal transduction and housekeeping genes generally (Table S6 and Auxiliary Spreadsheet 2), as has been found in other studies, e.g., Rensing ^71^. These genes, although maintained over millions of years and highly interconnected ^72^, can respond to selective pressures modifying their existing roles. For example, a recent study of silver birch found selective sweeps around candidate genes enriched among ancient polyploid duplicates that encode developmental timing and physiological cross-talk functions^7^. In oaks, it would be constructive to learn whether these ancient genes have undergone positive selection, allowing adaptation to new environments.

Genomes of high-quality document the deep evolutionary history of species. The oak genome has many features that provide hints of possible reasons for their success. Our exploration has uncovered several surprising similarities to the highly diverse grass genomes that may indicate analogous or even homologous adaptive strategies that would increase functional diversity in addition to the diversity generated by extensive gene duplications. Future oak studies may benefit by looking to the extensive experimental results from both wild and crop grasses for clues to potential mechanisms contributing to their evolutionary success.

## Methods

### Study species, samples, and genomic lab work

*Quercus lobata* Née (Fagaceae) is a widely-distributed endemic California oak species found in oak savannas, oak woodlands, and riparian forests. Oak have a highly outcrossed mating system^73^ with the potential for long distance gene flow occurring through wind-dispersed pollen with long-tailed distributions, despite many near-neighbor pollinations ^74,75^. Acorn dispersal is often restricted except for occasional long-distance colonization by jays ^76^. Occupying an unglaciated region of California, contemporary populations are at least 200k years old with no evidence of severe bottlenecks during cold periods ^77,78^ like those described for the European oaks from glaciation that retreated in the last 10k–20k years, allowing rapid recolonization from refugia in Italy and Spain ^79^. Valley oak and other California oak species have been used as a reliable food source and cultural resource by native peoples of western North America for the last 10k years ^80^. Since the arrival of Europeans, valley oak populations have experienced extensive habitat loss ^81^, and current population recruitment is jeopardized by cattle grazing, rodents, and other factors ^82,83^. Moreover, as its climate niche shifts north and upward ^82,84,85^, extant populations are further challenged by climate warming.

The focal tree for this study is *Q. lobata* adult SW786, which is located at the UC Santa Barbara Sedgwick Nature Reserve, is the same individual that was sequenced for version 1.0 of the genome ^9^. Leaf samples for the initial Illumina sequencing (532M paired-end [PE] 250 nt reads with ≈500 nt inserts giving 133 Gnt and ≈175x coverage, and 318M mate pair [MP] 150 nt reads from ≈3 knt to ≈12 knt fragments giving 48 Gnt and ≈56x coverage) were collected in September 2014, as described in Sork, et al. ^9^. Additional leaves were collected and DNA extracted in April 2016 for Pacific Biosciences whole genome SMRTbell libraries (6M reads of mean ≈9 knt and N50 ≈13 knt giving 58 Gnt and ≈80x coverage), and in March 2017 for Dovetail Chicago Hi-C library preparation. For details of the 19 whole genome resequencing libraries (Illumina PE, mean ≈24x coverage) used for the demographic analysis, three-tissue (bud, leaf, stem) Pacific Biosciences Iso-Seq and Illumina RNA-Seq transcriptome libraries contributing to annotation, and three-tissue (bud, catkin, and young leaf) whole-genome bisulfite libraries (Illumina SE, ≈18x – 19x coverage) for the DNA methylomes, (see SI Section I: Sample collection, library preparation, sequencing, and initial data processing).

### Genome assembly

We constructed the final genome in multiple stages. Stage 1: For the initial “Hybrid Primary” assembly (818 Mbp in 3.6k scaffolds, with longest 6.7 Mbp and NG50 ≈1.2 Mbp assuming at-the-time estimated 730 Mbp for the haploid genome), we applied MaSuRCA 3.2.1 ^86^ to our genomic Illumina PE, Illumina MP, and PacBio SMRT reads. The assembler identified high heterozygosity and selected diploid settings, allowing it to set aside most divergent haplotype variants; the result generally contains a single haplotype, but randomly phased, as we chose the larger scaffold whenever the assembler split two haplotypes into distinct scaffolds. Those scaffolds filtered out as alternative haplotypes were gathered into the “Hybrid Alternative” additions (466 Mbp in 17k scaffolds, with longest 1.2 Mbp). Stage 2: To assist completeness, we aligned to Stage 1 Primary+Alternative 82k of 84k transcripts and gene fragments from a prior RNA-Seq-derived transcriptome ^87^, with 81k aligning to Primary.

To avoid loss of potential coding regions, we moved 317 scaffolds from Alternative to Primary, forming the “Hybrid-plus-Transcript Primary” assembly (872 Mbp in 4.0k scaffolds, with longest 6.7 Mbp and NG50 ≈1.2 Mbp), and “Hybrid-plus-Transcript Alternative” additions (412 Mbp in 16k scaffolds, with longest 0.8 Mbp). Stage 3: We increased NG50 by aligning Stage 2 Primary scaffold ends with bwa mem ^88^, merging scaffolds that had unique end matches of > 94% identity longer than 40 kbp. This created the “Hybrid-plus-Transcript-Merged Primary” assembly (861 Mbp in 3.2k scaffolds, with longest 10.2 Mbp and NG50 ≈1.9 Mbp) and “Hybrid-plus-Transcript-Merged Alternative” additions (16k scaffolds). Stage 4: Next, we generated Hi-C long-range linking information from an Illumina-sequenced library produced by Dovetail Genomics, which we used to re-scaffold with HiRise ^10^ after read alignment with a modified SNAP (http://snap.cs.berkeley.edu), dramatically increasing NG50. Scores from the HiRise learned likelihood model were used to identify and break presumed misjoins, identify prospective joins, and commit joins above a threshold; shotgun reads from Stage 1 were used to close gaps where possible. Stage 5: Finally, after HiRise, any redundant haplotype contigs remaining (that truly belong in the same place as the other haplotype in a scaffolded assembly) are expected to be adjacent to the other haplotype as this is as close as they can be placed under the linear ordering constraint of HiRise output. We used this property to remove remaining extra haplotype contigs by aligning adjacent contigs to each other and finding those smaller than their direct neighbor that had > 50% syntenic alignment with the neighbor, thereby moving 14 Mbp to Alternative and forming the final “Hi-C-Scaffolded-plus-Neighbor-Cleaned Primary” (“version 3.0”) assembly (Figure 1C). The twelve longest scaffolds represent near full-length chromosomes (Figures S1 and S2) and total 811 Mbp (96%) of non-gap sequence.

### Comparisons of Q. lobata and Q. robur assemblies to linkage map

The *Q. robur* × *Q. petraea* linkage groups (LGs)^11^ are taken from http://arachne.pierroton.inra.fr/cgi-bin/cmap/map_set_info?map set acc=51 using Table S3 from Lepoittevin, et al. ^89^ as sequence-defined SNPs, dropping SNPs associated to more than one LG. Genomic locations were identified with BLASTN+ 2.2.30 (*E* < 10^−15^, word size 8), keeping for each query all alignments with bitscore ≥ 97% of the top bitscore. We plot SNPs that have either a unique surviving alignment, or multiple alignments but all to the same chromosome and with chromosomal span of hits ≤ 2 Mbp wide.

### Nucleotide- and amino acid-level alignments and 2-D visualizations of sequence similarity, K_s_, and shared Pfam domains, variously within and between genomes for Q. lobata, Q. robur, Q. suber, Populus, Eucalyptus, Theobroma, and Coffea

Various alignments and visualizations appear in Figures 1E, 4A, and 4B, and at the project website (TBD) and assisted in *Q. lobata* with discovery and identification of the tandem-like blocks of duplicated genes and syntenic self-syntenies (SSB). The principal software components were LASTZ for nucleotide alignments (with masked repeats from RepeatMasker after RepeatModeler); BLASTP, Diamond, and Parasail for homologous gene pair detection (on respective genome project protein-coding gene models) and subsequent detailed alignment refinement; C++, Mathematica, and Perl for scripting and pixel generation/import; and ImageMagick and Adobe Photoshop for manipulation, browsing, and annotation of generally multi-gigapixel images of high resolution (e.g., 10 kbp/pixel). Pfam hits were determined with InterProScan or direct HMMer runs. *K_s_* was computed for all homologous protein pairs discovered with other tools by re-aligning with ‘needle’ from EMBOSS (http://emboss.sourceforget.net/), converting to the level of codons with ‘pal2nal.pl’ (http://www.bork.embl.de/pal2nal/), and finally computing *K_s_* with ‘codeml’ from PAML (http://abacus.gene.ucl.ac.uk/software/paml.html).

### Demographic history

We inferred demographic history using the PSMC’ algorithm ^15^ by mapping the *Q. lobata* and *Q. robur* sequencing project genomic shotgun reads to their respective reference assemblies, as well as high-coverage genomic reads for 19 *Q. lobata* individuals to the *Q. lobata* assembly (Table S1). We called heterozygous sites in each genome (forming a VCF file with all callable sites) and composed input for PSMC’ with vcfAllSiteParser.py (https://github.com/stschiff/msmc-tools). Masking and filters are as described in SI Section IV: Demographic analysis — Input to PSMC’. We ran PSMC’ using default parameters except 200 for maximum number of iterations. Because PSMC’ inference can be prone to biases, we assessed robustness of our conclusions (Figure 2B). To determine if inference is affected by re-use of the same reads as used to build the reference assembly, we analyzed the 19 re-sequenced *Q. lobata* individuals beyond the reference individual SW786. These showed similar population size changes (Figure 2C light blue) as SW786 (Figure 2C dark blue), suggesting little bias from re-use. We also assessed if PSMC’ is capable of accurate inference by generating simulated datasets following the inferred demographic history, and re-ran inference on these (Figure 2D). These runs suggested that the only major issue was the oldest population sizes often being over-estimated. We thus selected ancestral population sizes matching empirical genome-wide heterozygosities (Figures S8, S9, and S10) and trimmed display in Figure 2C accordingly (Figure S7 exhibits untrimmed trajectories). Finally, we tested whether PSMC’ could reliably infer changes in population size on timescales relevant to *Quercus*. We simulated 10 test datasets of each run type under our presented demographic models (in Figure 2C) using the coalescent simulator msprime ^90^. With each simulated genome, we computed heterozygosity and used PSMC’ to infer demography; see SI Section IV: Demographic analysis — Simulations in msprime. We found accurate inference of population sizes over time, except for the single oldest time step where it tends to be over-estimated (Figure 2D). Note that inferred demographic trajectories from whole genome-based methods such as PSMC’ can be complex but not predict empirical summary statistics such as the genome-wide distribution of heterozygosity ^91^.

### Repetitive sequences

The primary repeat analysis is outlined in Figure 3A, and began with construction of a *Q. lobata*-specific database of repeat families by RepeatModeler/Classifier open-1.0.8, which was then applied with RepeatMasker open-4.0.6. Family consensus sequences are not always full length for their class or irredundant by close sequence similarity; we applied PSI-CD-HIT 4.7 to family consensus sequences at 45% nucleotide identity (the level where, as the threshold was lowered, intracluster similarities stopped falling in frequency and began rising) and chose a canonical rotation and strand for tandem repeat units, so as to cluster families into repeat “superfamilies” (SFs). Generally, each SF was assigned the RepeatClassifier class of the longest member of the family that was not unknown (if any; approximately two-thirds of SF-covered base pairs were classifiable). Annotated intervals for a SF are the nucleotide-level union of all intervals for member families, and SFs were assigned “s1RF####” accessions roughly serialized by descending mass. For certain uses (e.g., gene annotation), we also applied structurally-aware LTRharvest and LTRdigest from GenomeTools 1.5.9 to specifically target the abundant LTR TEs, identifying 28k instances of total mass 184 Mbp (not much larger than the 179 Mbp in LTR-classified SFs). Further details are in SI Section V: Repetitive sequences.

### Annotation of protein-coding genes

Figure 7A outlines dataflow of the PCG modeling process employed. ***Pure Iso-Seq models*.** The Pacific Biosciences of California, Inc. (PacBio) pipeline generated 197k-223k nominally full length non-chimeric polished transcripts (“reads”) from the poly-A-selected strand-specific bud, leaf, and stem PacBio-sequenced Iso-Seq libraries. Pooling tissues, Minimap2 aligned each read to zero to five reference genome locations (96%–99% uniquely); 11% of alignments were filtered out based on empirical criteria. Preliminary exon–intron structure was obtained by focusing on the reference side of each alignment, ignoring short insertions and merging short deletions and gapless blocks. Inspection found compact and well-isolated gene loci with generally concordant pileups at each of the 24k tentative loci, which had highly variable coverage (1 to 14k reads each; 56% ≥5, 22% =1). Inspection of higher-coverage loci found reads within pileups to vary: (i) at the exact coordinate level (with exon-intron boundaries moved by typically < 10 nt vs. common); (ii) at the structural level (introns resized or deleted or inserted, generally in a small minority of reads, and at more loci and in more ways than likely by alternative splicing); and (iii) in extent (with some reads truncated, especially at the 5’ end, with loss of multiple exons possible). A consensus exon–intron model per locus was generated by resolving (i) via rounding boundaries within ±25 nt to a most common boundary; (ii) by generally keeping exons and introns only in at least half of reads; and (iii) by extending 5’ and 3’ ends to the furthest extent observed. CDS assignments were made considering three methods: (i) longest ORFs; (ii) filtered (including restriction to only near-best hits per locus) BLASTP-equivalent (Diamond 0.9.22.123) alignments (*E* < 0.001) of translations of all ORFs to the entire NCBI 2018-05-18 ‘nr’ database (22k consensuses had at least one hit, with 83% of top hits involving at least half of both the translated ORF and the NCBI sequence, and with 99% having ≥ 50% amino acid identity, and 90% having *E* < 10^−35^; assignment of an ORF required agreement among all surviving hits); and (iii) a log2-odds bicodon coding potential trained using a selected subset of the NCBI analysis. Partial (and six-frame) ORFs were permitted. A consensus was assigned CDS (i.e., an ORF) if (ii) identified an ORF, the longest member of (i) was of the same frame with non-empty intersection with that ORF, and (iii) was also of the same frame and with non-empty intersection. This attributed CDS (and, hence, UTR5 and UTR3) to 19k loci, with 95% on the consensus read strand and 85% having ≥ 50 nt of both UTR5 and UTR3. Hand inspection of a random subset found them to be of generally good quality (often needing no edits). Diverse AUGUSTUS hints were constructed from the Iso-Seq reads and pure Iso-Seq models for eventual use in the final AUGUSTUS run near the end of the PCG modeling process.

### AUGUSTUS bootstrap

The 19k pure Iso-Seq models were filtered to a very high confidence subset of 2,639, then thinned to 2,558 by choosing single representatives from homology clusters determined via Exonerate 2.4.0 affine:local protein alignments. These were split uniformly at random into 1,698-model training and 860-model test sets and used to bootstrap AUGUSTUS via “new_species.pl” (enabling UTRs) and “etraining”, then optimized with “optimize_augustus.pl”, and also used to train the splice model of Exonerate. **RNA-Seq.** Paired end 101+101 nt Illumina HiSeq 4000 RNA-Seq reads were also collected from rRNA-depleted strand-specific bud, leaf, and stem libraries. The 121M to 153M pairs per tissue were aligned with STAR 2.5.3a and assembled into nominal transcripts with reference guidance by StringTie 1.3.4d and Trinity 2.6.6; Trinity output was aligned back to the reference genome with GMAP 2017-11-15. A large collection of diverse AUGUSTUS hints were constructed from STAR observed genomic base pair coverage and empirical splices, StringTie reference-quoted transcripts, and GMAP alignments. **Known proteins.** Protein translations of the *Q. robur* and *Q. suber* PCG models were BLASTP-equivalent (Diamond) aligned to a temporary trained/optimized but unhinted AUGUSTUS run generating and retaining a very large number of suboptimal isoform models. The resultant hits were used to identify regions of interest on the *Q. lobata* reference genome, that were then aligned vs. *Q. robur* and *Q. suber* in splice-discovery detail with splice model-trained Exonerate. Numerous strong AUGUSTUS hints were then constructed from Exonerate’s alignments. **Repeats.** Reference genome base pairs masked by RepeatMasker from the species-specific database constructed by RepeatModeler were weakly hinted to AUGUSTUS as non-exonic. **DNA mCHG and mCHH (BS-Seq) patterns.** Similar to repeats, sufficiently high mCHG or mCHH levels from merging the three tissues of the DNA methylation analyses were weakly hinted to AUGUSTUS as non-exonic. (Both *a priori* expectation and empirical examination of preliminary AUGUSTUS runs without methylation-based hinting had these marks as very highly anti-correlated with PCGs. mCG was not used, as it is complex, being high both in repeats and in many PCGs due to gene body methylation of non-short genes.)

### Main AUGUSTUS run

The above data sources provided 94M hinting intervals of 12 types tagged as from 62 sources. (As AUGUSTUS scores cannot be configured to be continuous functions of hint evidence strength [e.g., numeric coverage level from RNA-Seq], continuous strengths were generally broken into small numbers of discrete bins, with fixed scoring per bin.) A three-line patch (in extrinsicinfo.cc) to the AUGUSTUS C++ source code was required to enlarge hard-coded limits. One top isoform model per locus was predicted by the trained, optimized, UTR-aware, and now hinted main AUGUSTUS run. **Filtered final PCG models.** Numerous models from the main AUGUSTUS run were, e.g., clearly transposons with no or little evidence of observed expression. Based on several indicators (including Salmon-quantified per-model RNA-Seq expression, overlap with annotated repeats, presence of LTRdigest/harvest or GyDB/HMMer transposon domains, average mCG and mCHG and mCHH levels, and Diamond and BLASTP+ alignments with NCBI ‘nr’ and *Q. suber* and *Q. robur* PCGs), we removed such and other hypothetical models with poor evidence.

### Enrichment analyses

Benjamini-Hochberg false discovery rate (FDR)-adjusted hypergeometric *p*-values were used to determine Pfam do main enrichment in targeted subsets of tandemly duplicated genes and genes within SSBs.

### Methylomes and analysis of tissue-specific methylation patterns

Sample collection, library preparation, sequencing, and initial methylation calling are described in SI Section VII: Methylomes and analysis of methylation patterns. Libraries were prepared using the TruSeq Nano DNA (Illumina) and Epitek kits (Qiagen), and sequenced as 100 nt single end reads on an Illumina HiSeq 4000 to median coverage 18–19-fold. Methylation levels were determined using Methylpy v1.4.6 ^92^. DeepTools v3.1.2 ^93^ computeMatrix and plotProfile were used to assess methylation levels with respect to gene models and repeat superfamilies (Figures 5A–F, Figure 5I, Figures S15 and S17, with default parameters except as described in the legend). Methylation levels for 100 bp windows were calculated by dividing the total number of reads calling ‘T’ (= methylated) by the total number of informative reads (‘C’ or ‘T’) for all genomic cytosine positions in the appropriate sequence context within the window. Genome wide average methylation levels (Figure 5 and Figure S14) were calculated by averaging 100 bp window levels for the twelve chromosomal scaffolds. Per-site methylation levels in Figure S14 were calculated by dividing reads showing methylation (‘T’) by all informative reads (‘C’ or ‘T’) for each position, plotting with R ggplot2 v3.3.2 ^94^. Designation of genomic regions with respect to genes (1 kbp up, 5’ UTR, etc.) was done with Bedtools v2.27.1 (BEDTools: a flexible suite of utilities for comparing genomic features https://doi.org/10.1093/bioinformatics/btq033) and bed12toAnnotation.awk (https://github.com/guigolab/geneid/blob/master/scripts/bed12toAnnotation.awk). PCG model spans do not overlap in our annotation; however, overlaps for 1 kbp upstream and 1 kbp downstream regions were removed from the 1k up and 1k down categories, including overlaps that spanned neighboring genes. Gene regions overlapping with intervals (200 kbp to 3 Mbp) covering peri-centromere regions were removed. Introns were separated into first intron vs. other introns. Chromosome scale plots of subcontext methylation (Figure 5I and Figure 6A) were calculated with Bedtools as the mean of the percent methylation at each genomic cytosine position in the appropriate sequence context within each 1 Mb window, every 1 Mb. *Populus* methylation data^46^ was for Tree 13 branch 1, from GEO (https://www.ncbi.nlm.nih.gov/geo/query/acc.cgi?acc=GSE132939 2020). Local correlations between methylation levels and gene count were determined using methods from Niederhuth, et al. ^43^ to maximize relevance of the comparison. Thus, using Bedtools, the genome was divided into 100 kbp windows with 50 kbp overlaps. Methylation for each 100 kbp window was from averaging 100 bp window methylation levels (as above). Genes per window were counted with Bedtools intersect, requiring at least 50% of the gene span to be inside the window. Correlation between gene count and methylation level was calculated with R cor()’s Pearson method with incomplete observations dropped.

## Supporting information

1. Auxiliary Spreadsheet 1

2. Auxiliary Spreadsheet 2

## Data availability

Data are available at NCBI (GCA_001633185.3, additional accessions TBD), European Variation Archive (accession TBD), the project website (valleyoak.ucla.edu), and the project genome browser (genomes.mcdb.ucla.edu/cgi-bin/hgTracks?dg=queLob3).

## Acknowledgments

We acknowledge the native peoples of California whose relationship with the land has allowed the peoples of California to benefit from oak ecosystems, and allowed our research team to study oaks. We acknowledge the University of California Natural Reserve System, and especially the UC-Santa Barbara Sedgwick Reserve, home of tree SW786. We thank Krista Beckley, Dylan Burge, and Andy Lentz for field work; Krista Beckley for DNA extractions and library preparation; Marco Morselli for assistance with bisulfite sequencing library preparation; Suhua Feng for assistance with DNA and RNA sequencing; and Luke Browne for maps in Figure 2. Sequencing was conducted at the following core facilities: Illumina and PacBio were carried out at the DNA Technologies and Expression Analysis Cores of the UC Davis Genome Center, supported by NIH Shared Instrumentation Grant 1S10OD010786-01; and whole genome resequencing, WGBS, and RNA-Seq for demographics, methylation, and transcriptomic studies, respectively, were done at the UCLA Broad Stem Cell Genome Core facility. We thank Lily Shiue and Thomas Swale, Dovetail Genomics, for the high-quality scaffolding performed by HiRise.

## Author contributions

VLS, MP, and SLS conceived the overall project design and management and obtained grant support; VLS initiated, coordinated, and supervised the project and manuscript. SJC annotated and analyzed genes and repeats, designed and conducted genome comparative analyses, and created/wrote figures, results, methods, and supplementary information (SI) these sections. STF-G analyzed methylomes, designed and conducted comparative methylation analyses, created/wrote figures, results, discussion, methods, and supplementary information (SI) for this topic; called genetic variants (GATK) for the demographic analysis; submitted genomic resources for public availability. AVZ and DP assembled and curated the genome sequence and contributed to results, methods, and supplementary information (SI) for these sections. JG and YZ analyzed genetic variation data. JG conducted demographic analysis. KEL designed and supervised the demographic analysis and JG and KEL contributed results, methods, and SI for this section. STF-G and SJC examined DUF247. CLH conducted lab preparation for DNA sequencing, resequencing, bisulfite sequencing and RNA sequencing. VLS, SJC, STF-G, AVZ, JG, PFG, KEL, MP, and SLS edited text. SJC edited manuscript figures. SJC, STF-G, and VLS interpreted data and wrote the manuscript.

