## Supplementary Information: methods, figures, tables for "High-quality genome and methylomes illustrate features underlying evolutionary success of oaks"

**Affiliations:**

#### Auxiliary documents separate from this Supplementary Information:

1. **Auxiliary Spreadsheet 1**: AuxSpread1_MethylationTrendsAdaptedFromNiederhuth.xlsx
   Adapted from Table S2 in Additional files associated with "Widespread natural variation of DNA methylation within angiosperms"[1] (https://genomebiology.biomedcentral.com/articles/10.1186/s13059-016-1059-0), giving basic methylation statistics for 34 angiosperms, to which we have added data for oaks, plus additional columns derived from the original data: (1) rank of correlation of mCHH vs. gene content; (2) estimate of chromosome arm gene density; and (3) strength of mCHH islands, both with and without respect to background mCHH level

2. **Auxiliary Spreadsheet 2**: AuxSpread2_SJC-PCG-HgeoEnrichments--GeneNameWord-Pfam—20200717.xlsx. Hypergeometric enrichment of words found in gene names and (separately) Pfam domains, for subsets of protein coding genes (PCGs) defined variously by degree of tandem duplication, membership in self-syntenic blocks (SSBs), intron size, expression level, protein length, and Local repeat density

### TABLE OF CONTENTS FOR SUPPLEMENTAL INFORMATION

### I. Sample collection, library preparation, sequencing, and initial data processing

#### A. Valley oak reference genome

All tissues collected from either *Quercus lobata* SW786 at Sedgewick Reserve in Santa Barbara, CA, or other *Q. lobata* trees throughout the California species range (Table S1) were placed immediately on dry ice. Plant tissue was stored at –80 °C until the day of extraction. The voucher specimen for tree SW786, collected March 2017, is D. O. Burge 2309, deposited at UC Davis (DAV). This healthy and prolific acorn producing adult has been included in several quantitative genetic and genomic studies [2-6].

***Illumina paired end and mate pair libraries.*** Leaf tissue for Illumina libraries was collected September 2014. Details for extraction of total genomic DNA, library preparation, and sequencing are described in Sork, et al. [7]. Briefly, DNA extractions were by a CTAB protocol. 266M HiSeq 2500 read pairs of 250 nt (175x coverage) were generated from two short insert paired end libraries, one with PCR enrichment and one without. 159M HiSeq 2500 read pairs of 150 nt (56x coverage) were generated from nine mate pair libraries of length 2.9 kb to 12 kb.

***Pacific Biosciences whole genome SMRTbell libraries.*** Leaf tissue for the PacBio DNA libraries was collected April 2016. High molecular weight (HMW) DNA was obtained through a nuclei isolation protocol based on "Preparing *Arabidopsis* Genomic DNA for Size-Selected ~20 kb SMRTbell™ Libraries" (Pacific Biosciences of California, Inc., 2013) and the Sean Gordon protocol [8]. Ten grams of fresh plant tissue was flash frozen with liquid nitrogen and ground with a mortar and pestle three times to obtain a fine powder, and transferred to a chilled Erlenmeyer flask. 300 mL of fresh sucrose-based extraction buffer (SBE) was prepared (2% w/v PVP, 10% v/v TKE, 500 mM sucrose, 4 mM spermidine trihydrochloride, 1 mM spermine tetrahydrochloride, 0.1% w/v ascorbic acid, 0.13% w/v sodium diethyldithiocarbamate, and adjusted to a pH of 9.0–9.1 with 1M KOH) with 600 µL of ß-mercaptoethanol (BME). 185 mL of SBE+BME was added to the ground tissue and placed on ice for 12–20 minutes with continuous swirling until the powder dissolved. The homogenate was filtered through two layers of Grade 50 cheesecloth (Lions Services, North Carolina, USA) into a clean 500 mL beaker, using an extra 15 mL of SBE+BME solution to ensure all particulates passed through the cheesecloth. Then, 10 mL of cold 10% Triton was added to the beaker, slowly along the side over the course of two minutes, while gently stirring with a magnetic bar, then kept on ice for eight minutes with intermittent gentle swirling. The mixture was then transferred into 4x 50 mL polypropylene Falcon tubes, and spun in a centrifuge at 650 x *g* (1,970 rpm) for 15 min at 4 °C. The supernatant was discarded and the pellet was gently resuspended in 10 mL of cold SBE+BME. The mixture was then transferred into 2x 50 mL polypropylene Falcon tubes and SBE+BME was added until each tube had a final volume of 30 mL. These were centrifuged at 650 x *g* (1,970 rpm) for 15 min at 4 °C. The supernatants were discarded and the pellets were resuspended in 1.44 mL of TE. The mixture was then divided into 4x 2mL tubes and 95 µL of cold 1M NaCl and 240 µL of cold 10 mg/ml RNase A was added to each tube and incubated at 65 °C for 30 min to digest RNA. Then 24 µl of cold 10 mg/ml Proteinase K was added to each tube and inverted gently 2x. Then 95 µl of room-temperature 10% SDS was added to each tube, inverted gently 2x, incubated at 45 °C for 60 min to digest proteins, then brought to room temperature. Samples from 2 mL tubes were combined into 2x 15mL Falcon tubes and 2.178 mL (or 1 volume) of phenol:chloroform:isoamyl alcohol was added and tubes were inverted gently, vortexed for two seconds, then centrifuged for five minutes at room temperature at 1,500 x g (2,300 rpm). The aqueous layer was transferred to new 15 mL Falcon tubes and the extraction with 1 volume of phenol:chloroform:isoamyl alcohol was repeated until the interface was clear. The clear extraction was then divided into 6x 2 mL tubes (~670 µL in each tube) and 70 µL (or ~0.1 volume) of 3M NaOAc (pH 5.2), and 750 µL (or ~1 volume) cold isopropanol was added and placed in the –20°C freezer for 30–60 minutes or at 4 °C overnight. The tubes were then centrifuged for 30 minutes at 13,000 rpm at 4 °C, then the supernatants were discarded. The pellets were then washed 2x with 500 µl 70% ethanol, centrifuged for >10 min at 13,000 rpm at 4 °C. Pellets were spun for two minutes at 13,000 rpm at 4 °C and the ethanol was decanted with a pipette. The pellets were then air dried at room temperature for 10 minutes, then resuspended in 30–50 µL TE per tube and allowed to rest in a 2–8°C fridge overnight to elute DNA. DNA was analyzed using the Nanodrop ND-1000 Spectrophotometer (Thermo Fisher Scientific, Waltham, MA), and run on a 0.8% agarose gel with 1 kb plus ladder and quantified using the Qubit 3.0 Fluorometer (Life Technologies, Carlsbad, CA).

The HMW DNA samples were sent to the DNA Technologies & Expression Analysis Core Laboratory at the University of California, Davis. The samples were purified using the "Guidelines for Using a Salt:Chloroform Wash to Clean up gDNA" protocol (Pacific Biosciences of California, Inc., 2014), then prepared into libraries using the "Procedure & Checklist – Preparing > 30 kb SMRTbell™ Libraries Using the Megaruptor® Shearing and BluePippin™ Size-Selection System" Protocol (Pacific Biosciences of California, Inc., 2016). Three libraries were prepared with 8 kb lower cut-off, 10 kb lower cut-off, and 20 kb lower cut-off size selections by BluePippin™ (Sage Science, Beverly, MA, USA). Seventeen v3 SMRT cells were run for the 8 kb cut-off, 11 cells for the 10 kb cut-off, and eight cells for the 20 kb cut-off. Sequencing polymerase was version 6 and chemistry was version 4 (P6C4). SMRT cells were sequenced on the RS II sequencer yielding 80x genome coverage.

***Dovetail whole genome HiC library.*** Leaf tissue was collected March 2017, of which 1 gram was sent to Dovetail Genomics, Scotts Valley, CA, USA. A Dovetail HiC library was prepared in a similar manner as described previously [9]. Briefly, for each library, chromatin was fixed in place with formaldehyde in the nucleus and then extracted. Fixed chromatin was digested with DpnII, the 5' overhangs filled in with biotinylated nucleotides, and then free blunt ends were ligated. After ligation, crosslinks were reversed and the DNA purified from protein. Purified DNA was treated to remove biotin that was not internal to ligated fragments. The DNA was then sheared to ~350 bp mean fragment size and sequencing libraries were generated using NEBNext Ultra enzymes and Illumina-compatible adapters. Biotin-containing fragments were isolated using streptavidin beads before PCR enrichment of each library. The libraries were sequenced on an Illumina HiSeq X to produce 454M 151+151 bp paired end reads, which provided 6,875x physical coverage of the genome (10–10,000 kb pairs).

#### B. Valley oak resequenced genomes

Leaf tissue samples for whole genome sequencing used in the demography studies (described below in Section IV: Demographic analysis) were collected from 19 *Q. lobata* adults (Table S1). Total genomic DNA was extracted from frozen leaf tissue using a prewash method [10] followed by a modified CTAB protocol [11] or the Plant DNeasy Kit protocol (Qiagen, Germany). Plants were frozen in liquid nitrogen and ground using a Mixer Mill MM301 (Retsch, Germany). The prewash method was repeated up to 3x until a clear supernatant was achieved. The resultant pellet was then used in a modified CTAB protocol in which the chloroform-isoamyl (24:1) step was repeated twice. DNA was analyzed using the Nanodrop ND-1000 Spectrophotometer (Thermo Fisher Scientific, Waltham, MA), and quantified using the Qubit 3.0 Fluorometer (Life Technologies, Carlsbad, CA).

Libraries were prepared following the Nextera XT DNA Library Prep Kit guidelines (Illumina, San Diego, CA). Dual index combinations for each sample were chosen based on the Nextera Low Plex Pooling Guidelines (Illumina, San Diego, CA). Samples were multiplexed in the following layout: eight lanes of six libraries per lane on 2016–12–09; three lanes of eight libraries per lane on 2017–10–11; seven lanes of 3–4 libraries per lane on 2018–09–06 based on coverage needs, and 11 lanes of 2–libraries per lane on 2019–04–01 based on coverage needs. Libraries were analyzed on the Agilent D1000 Screen Tape System on the Agilent 2200 TapeStation (Agilent Technologies, Santa Clara, CA, USA), and sequenced using an Illumina HiSeq 4000at the UCLA Stem Cell Center Core facility with 100 bp paired end reads to a coverage 17x–32x (mean 24x) (assessed by GATK v3.7-0-gcfedb67 DepthOfCoverage `--countType COUNT_FRAGMENTS --minMappingQuality 20 --minBaseQuality 10`).

Illumina reads were adapter trimmed and quality checked using TrimGalore! v.0.4.4 https://www.bioinformatics.babraham.ac.uk/projects/trim_galore/, calling Cutadapt 1.9.1 [12] with no quality trimming and minimum length 20 bp. Trimmed reads were aligned to the *Q. lobata* 3.0 reference genome using bwa mem v.0.7.12-r1039 (https://www.ncbi.nlm.nih.gov/pmc/articles/PMC2705234/). Read duplicates were flagged using Picard tools MarkDuplicates v.2.13.2-SNAPSHOT (http://broadinstitute.github.io/picard/). Variants and non-variants were called for all sites of each sample with GATK v3.7-0-gcfedb67 HaplotypeCaller `--heterozygosity 0.01 --indel_heterozygosity 0.001 -newQual --emitRefConfidence GVCF`, followed by genotyping of the whole population together with GATK v3.7-0-gcfedb67 GenotypeGVCFs with `--includeNonVariantSites --heterozygosity 0.01 --indel_heterozygosity 0.001`.

**Table S1.** Collection of resequenced *Q. lobata* adults. Sample IDs and locations of 19 *Q. lobata* adults sampled throughout the species range for whole genome resequencing in the demography study.

| Sample ID | Locality Name | Latitude (°) | Longitude (°) |
|---|---|---|---|
| QL.CHE.100 | Cheeseboro (CHE) | 34.1636 | –118.7241 |
| QL.CHI.3 | Chico (CHI) | 39.7119 | –121.7842 |
| QL.CLO.4 | Clearlake Oaks (CLO) | 39.0219 | –122.7135 |
| QL.CVD.8 | Cloverdale (CVD) | 38.8544 | –123.0319 |
| QL.FHL.5 | Fort Hunter Liggett (FHL) | 35.9804 | –121.2328 |
| QL.GRV.2 | Gravelly Valley (GRV) | 39.4302 | –122.9754 |
| QL.GRV.7 | Gravelly Valley (GRV) | 39.4485 | –122.9640 |
| QL.JAS.5 | Jasper Ridge (JAS) | 37.4032 | –122.2436 |
| QL.LAY.5 | Laytonville (LAY) | 39.7460 | –123.5242 |
| QL.LAY.6 | Laytonville (LAY) | 39.6722 | –123.4807 |
| QL.LYN.4 | Lynch Canyon Road (LYN) | 35.7878 | –120.9391 |
| QL.MAR.B | Mariposa (MAR) | 37.4611 | –119.8797 |
| QL.MCK.5 | Middle Creek CG (MCK) | 39.2524 | –122.9516 |
| QL.MOH.3 | Morgan Hill (MOH) | 37.1649 | –121.7148 |
| QL.MTR.3 | Mountain Ranch (MTR) | 38.2750 | –120.5058 |
| QL.PEN.5 | Penn Valley (PEN) | 39.2034 | –121.1902 |
| QL.ROV.3 | Round Valley (ROV) | 39.7483 | –123.2484 |
| QL.SUN.5 | Sunol (SUN) | 37.5987 | –121.8751 |
| QL.UKI.5 | Ukiah (UKL) | 39.0924 | –123.2197 |

#### C. Transcriptomes

***Pacific Biosciences RNA long read (Iso-Seq) libraries for tree SW786 bud, leaf, and stem tissues.*** Bud, leaf and stem tissue samples for Iso-Seq libraries were collected from tree SW786 in October 2017. RNA extractions were performed between November 6–8, 2017 using a modified version of the Conifer RNA prep protocol from the Cronn Lab (https://openwetware.org/wiki/Conifer_RNA_prep) and the Spectrum Plant Total RNA kit (Sigma, St. Louis, MO, USA). Plant tissues (100 mg each of leaves, buds, and stems) were flash frozen in liquid nitrogen and ground with a mortar and pestle to a fine powder. Powdered tissues were transferred to cold 2 mL tubes and 1.8 mL of cold RNA Extraction Buffer + DTT was added. RNA Extraction Buffer consists of 8M Urea, 3M LiCl, 1% polyvinylpyrrolidone K-60, 5 mM DTT (added just before use; 1M stock). Tubes were then vortexed for 30 seconds, incubated at 4 °C for 30 minutes, then centrifuged at 4 °C for 30 minutes at 20,000 rcf. The supernatant was discarded and the pellet was used as the starting material for the Spectrum Plant Total RNA kit Protocol A, adding 750 µL of Binding Solution, and performing on-column DNase I digestion. RNA quality and quantity were assessed using the Agilent RNA ScreenTape System on the Agilent 2200 TapeStation system (Agilent Technologies, Santa Clara, CA, USA).

RNA was further prepared following the "Guidelines for Preparing cDNA Libraries for Isoform Sequencing (Iso-Seq™) User Bulletin" (Pacific Biosciences of California, Inc., 2014) and the "Procedure & Checklist – Iso-Seq™ Template Preparation for Sequel™ Systems" (Pacific Biosciences of California, Inc., 2017). PolyA$^+$ RNA was extracted from total RNA using the Ambion® Poly(A) Purist™ MAG Kit (Invitrogen, Carlsbad, CA, USA) following the manufacturer's protocol. First strand cDNA synthesis was performed using the SMARTer® PCR cDNA Synthesis Kit (Takara Bio, Inc., Kusatsu, Shiga Prefecture, Japan), with three reactions using 3.5 µL of PolyA$^+$ RNA per tissue, for a total of nine first strand synthesis reactions. Each of nine reactions were diluted with 90 µL of EB buffer, then pooled according to tissue type. PCR cycle optimization resulted in the following PCR conditions, with 24x 50 µL reactions per tissue type: 95 °C for 2 minutes for initial denaturation, then *n* cycles (*n* = 10, 12, and 14 for leaf, bud, and stem) of 98 °C for 20 seconds, 65 °C for 15 seconds, 72 °C for 4 minutes, then 72 °C for 5 minutes for the final extension. Twelve reactions per tissue type were pooled for 1X AMPure XP (Beckman Coulter, Inc., Pasadena, CA, USA) bead purification, and 12 reactions per tissue type were pooled for 0.4X AMPure XP bead purification. Samples were sent to the DNA Technologies & Expression Analysis Core Laboratory at the University of California, Davis for size selection, enrichment, library preparation, and sequencing. A second bead size selection was performed, 1X and 0.4X (Fractions 1 and 2, respectively) for two size fractions and a size selection of 5–10 kb using the BluePippin Size Selection System. Six libraries were made from these different size selections following the "Procedure & Checklist – Iso-Seq™ Template Preparation for Sequel™ Systems" protocol: Libraries 1, 2, and 3 (leaf, bud, and stem) full length (Fractions 1 and 2), and Libraries 4, 5, and 6 (leaf, bud, and stem) 5–10 kb size selection (Fraction 3). For each of bud, leaf, and stem, libraries were pooled for sequencing (5:1, full length: Fraction 3) for a total of three libraries that were each sequenced on a single cell. The cells were loaded on the Sequel by Magbead/ v2 SMRT cell/ P2.1C2.1 (polymerase version 2.1 and chemistry version 2.1).

Raw reads (from subreads BAM files) for each of the three tissues were processed using PacBio's Iso-Seq classify bioinformatics pipeline [8], although clustering was skipped and replaced with filtering of the Minimap2 read alignments (described in the methods section of the manuscript). Specifically, the two Iso-Seq classify bioinformtaics pipeline steps were: 1.ccs with --minLength=50 --maxLength=12000 --minPasses=1 --minPredictedAccuracy=0.8 --minZScore=-999 --maxDropFraction=0.8, and 2. pbtranscript classify with --min_seq_len 100. The resulting putatively full length non-chimeric reads were aligned to the genome with intron-enabled Minimap2 [13] using -ax splice -uf --secondary=no. Final aligned reads and raw subread bam files are available from NCBI accession TBD.

***Illumina RNA short read (RNA-Seq) libraries for tree SW786 bud, leaf, and stem tissues.*** Bud, leaf, and stem tissue samples for RNA-Seq libraries were collected from tree SW786 in October 2017. RNA was extracted February 16, 2018. Total RNA was depleted of rRNA using the Ribo-Zero rRNA Removal Kit (Plant Leaf) (Illumina, San Diego, CA, USA). Total RNA input amounts were 4.2 µg for bud, 5 µg for leaf, and 4.1 µg for stem. The Individual Washing option was used for washing the magnetic beads, and the 500 ng–to–1.25 µg input RNA recipe was used for hybridizing the probes. RNA samples depleted of rRNA were cleaned with ethanol precipitation, incubated with Elute, Prime Fragment High Mix at 85 °C for 6 minutes, and quantified using the Qubit® RNA HS Assay Kit with the Qubit 3.0 Fluorometer (Life Technologies, Carlsbad, CA). Libraries were constructed using the TruSeq Stranded Total RNA protocol with a positive control (Illumina, San Diego, CA, USA). First strand and second strand cDNA synthesis, dA-tailing, ligation, purification, and enrichment steps were performed following the manufacturer's instructions (Illumina, San Diego, CA, USA). Libraries were analyzed using the Agilent D1000 Screen Tape System on the Agilent 2200 TapeStation (Agilent Technologies, Santa Clara, CA, USA). Fragments were found to be too small (~275 bp), so an extra size selection step was performed with AMPure XP beads at a concentration of 0.65x, to yield fragments in the 400–700 bp range. Libraries were quantified using the Qubit® dsDNA BR Assay Kit on the Qubit® 3.0 Fluorometer (Life Technologies, Carlsbad, CA). Libraries were pooled and sequenced on an Illumina HiSeq 4000 at the UCLA Broad Stem Cell Center Core facility.

#### D. Methylomes

***Whole genome bisulfite libraries for tree SW786 bud, catkin, and young leaf tissues.*** Tissue samples for assaying methylation were collected from tree SW786 on three different months of 2017: February (bud), March (catkin), and April (young leaves). Total genomic DNA was extracted from frozen leaf tissue on August 24, 2017 using a prewash method (Li et al., 2007) followed by a modified CTAB protocol (Doyle and Doyle, 1987). Plants were frozen in liquid nitrogen and ground using a Mixer Mill MM301 (Retsch, Germany). The prewash method was repeated up to 3x until a clear supernatant was achieved. The resultant pellet was then used in a modified CTAB protocol in which the chloroform-isoamyl (24:1) step was repeated twice. Total genomic DNA at a concentration of 500 ng in 60 µL was sonicated using an S2 Focused-ultrasonicator (Covaris, Woburn, MA, USA) for 60 seconds to obtain fragments in the 200–300 bp range (duty cycle: 10%, intensity: 5, cycles/burst: 200, mode: frequency sweeping). Using reagents from the TruSeq Nano DNA Library Prep Kit (Illumina, San Diego, CA, USA), sheared DNA samples were end repaired as in the TruSeq protocol, then purified with AMPure beads at a concentration of 1.6x. Fragments were then adenylated and adapters ligated as in the TruSeq protocol, except that 1 µL of Illumina TruSeq Adapters were used in the final reactions. The ligation reactions were purified with AMPure beads at a concentration of 1.2x, then purified with beads again at a concentration of 1x. Samples were then treated with bisulfite using the EpiTect kit (Qiagen, Hilden, Germany) according to the manufacturer's protocol, with one exception in that the bisulfite DNA conversion was performed twice for a total of 10 hours of incubation. Two amplification reactions were then performed for each sample (20 µL of bisulfite converted DNA, 2.5 µL Illumina TruSeq primer cocktail, 25 µL MyTaq Mix (Bioline, Taunton, MA), 2.5 µL $H_2O$ per PCR reaction) under the following conditions: initial denaturation at 98 °C for 30 s; 12 cycles of 98 °C for 15 s, 60 °C for 30 s, 72 °C for 30 s; final extension at 72 °C for 5 min. The final PCR products were purified using AMPure XP beads. Libraries were analyzed on the Agilent D1000 Screen Tape System on the Agilent 2200 TapeStation (Agilent Technologies, Santa Clara, CA, USA). All samples were sequenced once on a single Illumina HiSeq 4000 lane with 100 bp single end reads at the UCLA Broad Stem Cell Core facility, yielding median genomic coverage of 18x–19x.

Reads were trimmed and inspected with Trim Galore! v0.4.4 [14], which calls Cutadapt [15] and FastQC [16], with a quality score cutoff of 20 and minimum length of 80 bp. Trimmed reads were aligned to the *Q. lobata* 3.0 reference genome using Methylpy v1.4.6 [17], which converts the reference genome for alignment of BS-Seq data, aligns with Bowtie2 [18], estimates the bisulfite non-conversion rate from an unmethylated control (in our case, the *Q. lobata* chloroplast), performs binomial tests to distinguish methylated sites above the estimated non-conversion noise level, and outputs counts of covering methylated and unmethylated reads for each genomic cytosine site. Parameters for the methylpy single end pipeline command were `--remove-clonal True --min-mapq 30 --min-base-quality 1 --trim-reads False --unmethylated-control chrC --binom-test True --min-cov 3`. Aligned reads were inspected for methylation bias by read position using MethylDackel v0.4.0 mbias [19] and sequencing depth was assessed using DeepTools v.3.1.2 plotCoverage[20].

#### II. Validation and orientation of chromosomes

To confirm the correspondence of the twelve longest *Q. lobata* 3.0 assembly scaffolds as chromosomes, we used an existing moderate density physical map of *Q. robur* x *Q. petraea* ("2015 composite") [21] consisting of 4,217 SNP markers (after dropping 22 named SNPs associated to two physical locations each) in twelve linkage groups (LGs). *Q. robur* is also in section *Quercus* and is probably separated from *Q. lobata* by 30M years [22]. These SNPs are a subset of 7,913 [23] identified by more than 100 nt context each (typically 100 nt on both sides). We aligned marker sequences to our assembly with BLASTN 2.2.30+ ($E < 10^{-15}$), retaining all hits for each query with ≥ 97% bitscore of the top hit. Approximately 82% of the 7,913 were genetically mapped to *Q. lobata* uniquely, 14% to exactly two locations, 3% to more than two, and 1% were unmapped; all hits had nt identity > 69% and aligned ≥ 57 nt, and 90%+ variously had nt identity > 93%, aligned ≥ 105 nt, covered > 52% of the query, and had $E \leq 10^{-42}$. Of the 4,217 SNPs on an LG, we dropped 1% that were genetically unmapped to *Q. lobata*, kept 86% uniquely mapped, dropped 5% mapped to multiple scaffolds, kept 8% that were multiply mapped but to a single scaffold with span of all hits ≤ 2 Mbp wide, and dropped 0.5% that were multiply mapped with wider spans. Analysis with the *Q. robur* assembly was with the same procedure and parameters. We found a predominantly monotonic one-to-one correspondence between LGs and the twelve largest scaffolds of our assembly (Figure S1, Figure S2, and Table S2), and thus renamed our scaffolds as chromosomes, adopting the LG (and *Q. robur*) numbering (but not necessarily the LG orientation, where we instead follow the *Q. robur* assembly — hence, we essentially adopt both the numbering and orientation of *Q. robur*).

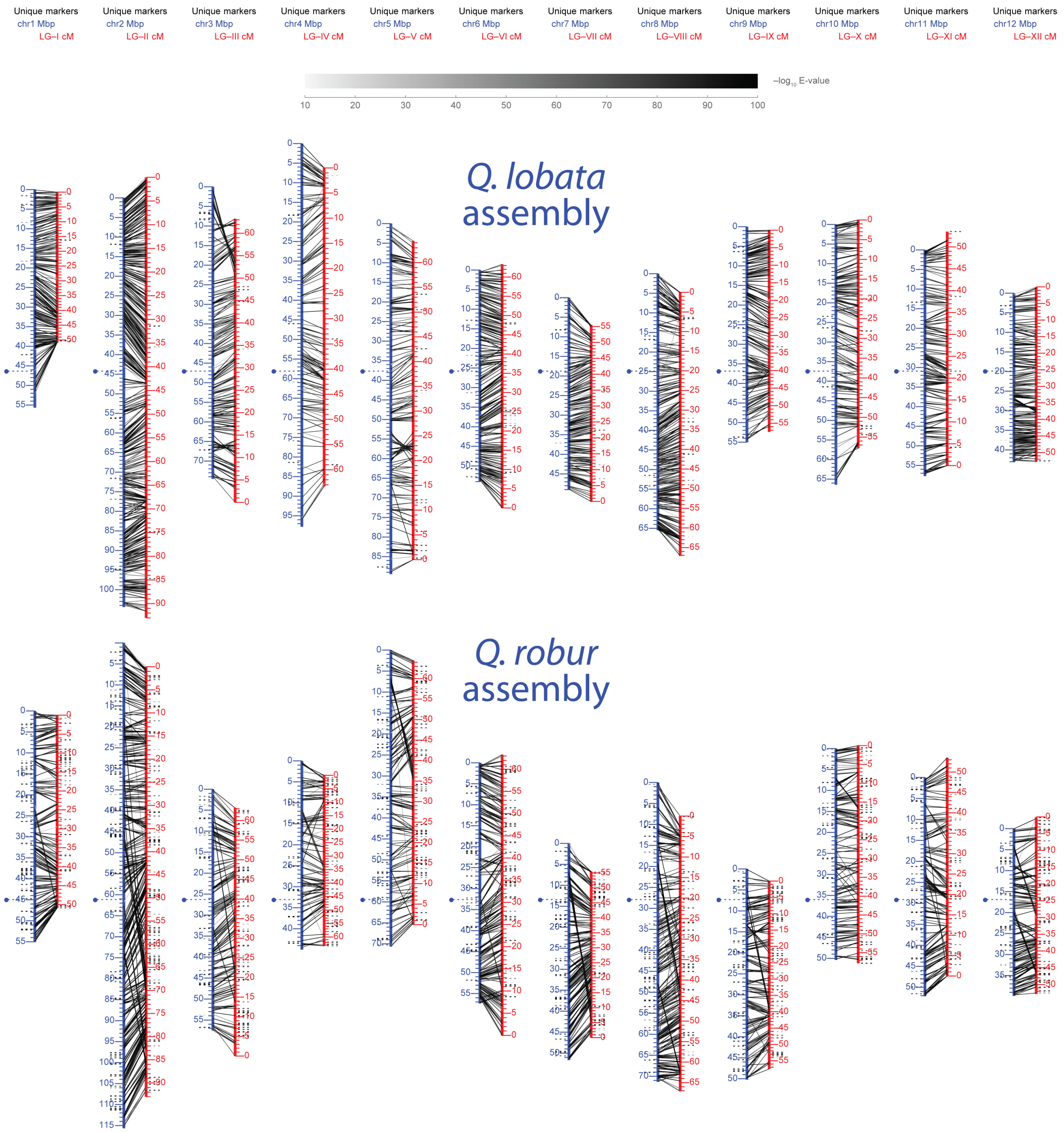

**Figure S1.** *Q. lobata* and *Q. robur* assemblies vs. *Q. robur* x *Q. petraea* linkage map: 1-D view. Lines connect sequence context-defined SNPs in the physical map (blue centimorgan scales) to assembly locations (red Mbp scales, via sequence alignment of the typically ±100 nt of context for the SNP).

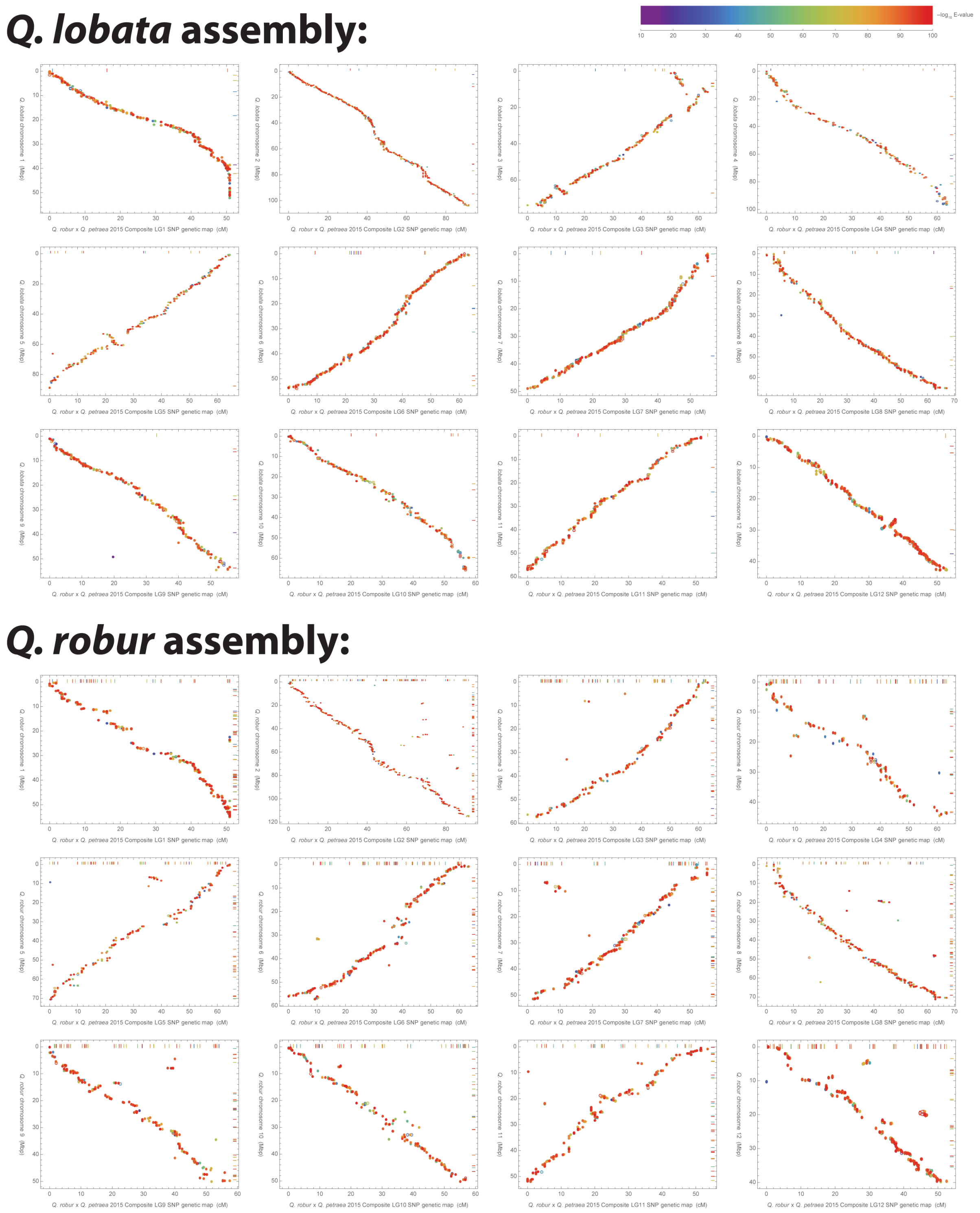

**Figure S2.** *Q. lobata* and *Q. robur* assemblies vs. *Q. robur* x *Q. petraea* linkage map: 2-D view. While both assemblies have each scaffold that is declared chromosomal stand predominantly in a one-to-one monotonic relationship with the physical map LG of the same number, the *Q. lobata* assembly (which did not use the physical map for sequence construction) shows many fewer anomalies (despite being more distant to the map's cross). Points along very top and right edges of plots are those of pairings where chromosome and LG number disagree.

**Table S2.** Statistics of *Q. lobata* and *Q. robur* assemblies vs. *Q. robur* x *Q. petraea* linkage map.

| ***Q. lobata:*** | | | | | | | | |
|---|---|---|---|---|---|---|---|---|
| **[A]** | **[B]** | **[C]** | **[D]** | **[E]** | **[F]** | **[G]** | **[H]** | **[I]** |
| | | | **uniquely** | | | | **multiply** | |
| **Physical map linkage group** | # SNPs physical map assigns to just this LG | filter [B] to those with ≥1 *Q. lobata* alignment | filter [C] to those with all alignments to same *Q. lobata* chrom./scaffold | filter [D] to those aligning to chr1..12 | filter [E] to those aligning to same chr# as LG# | *[F] as % of [C]* | filter [C] to those with ≥1 alignment to same chr# as LG# | *[H] as % of [C]* |
| **LG I** | 308 | 308 | 300 | 300 | 297 | *96.4%* | 303 | *98.4%* |
| **LG II** | 706 | 703 | 668 | 668 | 663 | *94.3%* | 697 | *99.1%* |
| **LG III** | 299 | 294 | 289 | 289 | 284 | *96.6%* | 289 | *98.3%* |
| **LG IV** | 227 | 227 | 211 | 210 | 207 | *91.2%* | 223 | *98.2%* |
| **LG V** | 298 | 297 | 271 | 271 | 259 | *87.2%* | 283 | *95.3%* |
| **LG VI** | 404 | 392 | 374 | 374 | 359 | *91.6%* | 376 | *95.9%* |
| **LG VII** | 325 | 322 | 301 | 301 | 296 | *91.9%* | 317 | *98.4%* |
| **LG VIII** | 433 | 423 | 405 | 405 | 395 | *93.4%* | 412 | *97.4%* |
| **LG IX** | 289 | 286 | 272 | 272 | 270 | *94.4%* | 284 | *99.3%* |
| **LG X** | 288 | 288 | 271 | 271 | 266 | *92.4%* | 283 | *98.3%* |
| **LG XI** | 294 | 293 | 280 | 280 | 275 | *93.9%* | 286 | *97.6%* |
| **LG XII** | 346 | 341 | 336 | 336 | 335 | *98.2%* | 340 | *99.7%* |
| ***total*** | ***4,217*** | ***4,174*** | ***3,978*** | ***3,977*** | ***3,906*** | ***93.6%*** | ***4,093*** | ***98.1%*** |

| ***Q. robur:*** | | | | | | | | |
|---|---|---|---|---|---|---|---|---|
| **[A]** | **[B]** | **[C]** | **[D]** | **[E]** | **[F]** | **[G]** | **[H]** | **[I]** |
| | | | **uniquely** | | | | **multiply** | |
| **Physical map linkage group** | # SNPs physical map assigns to just this LG | filter [B] to those with ≥1 *Q. robur* alignment | filter [C] to those with all alignments to same *Q. robur* chrom./scaffold | filter [D] to those aligning to chr1..12 | filter [E] to those aligning to same chr# as LG# | *[F] as % of [C]* | filter [C] to those with ≥1 alignment to same chr# as LG# | *[H] as % of [C]* |
| **LG I** | 308 | 305 | 299 | 289 | 242 | *79.3%* | 245 | *80.3%* |
| **LG II** | 706 | 696 | 688 | 662 | 569 | *81.8%* | 574 | *82.5%* |
| **LG III** | 299 | 297 | 287 | 273 | 208 | *70.0%* | 217 | *73.1%* |
| **LG IV** | 227 | 224 | 204 | 181 | 140 | *62.5%* | 149 | *66.5%* |
| **LG V** | 298 | 292 | 286 | 267 | 216 | *74.0%* | 218 | *74.7%* |
| **LG VI** | 404 | 400 | 390 | 374 | 312 | *78.0%* | 320 | *80.0%* |
| **LG VII** | 325 | 321 | 314 | 294 | 261 | *81.3%* | 267 | *83.2%* |
| **LG VIII** | 433 | 426 | 410 | 389 | 362 | *85.0%* | 375 | *88.0%* |
| **LG IX** | 289 | 286 | 275 | 256 | 210 | *73.4%* | 217 | *75.9%* |
| **LG X** | 288 | 284 | 272 | 264 | 220 | *77.5%* | 228 | *80.3%* |
| **LG XI** | 294 | 292 | 283 | 277 | 242 | *82.9%* | 247 | *84.6%* |
| **LG XII** | 346 | 342 | 339 | 322 | 253 | *74.0%* | 256 | *74.9%* |
| ***total*** | ***4,217*** | ***4,165*** | ***4,047*** | ***3,848*** | ***3,235*** | ***77.7%*** | ***3,313*** | ***79.5%*** |

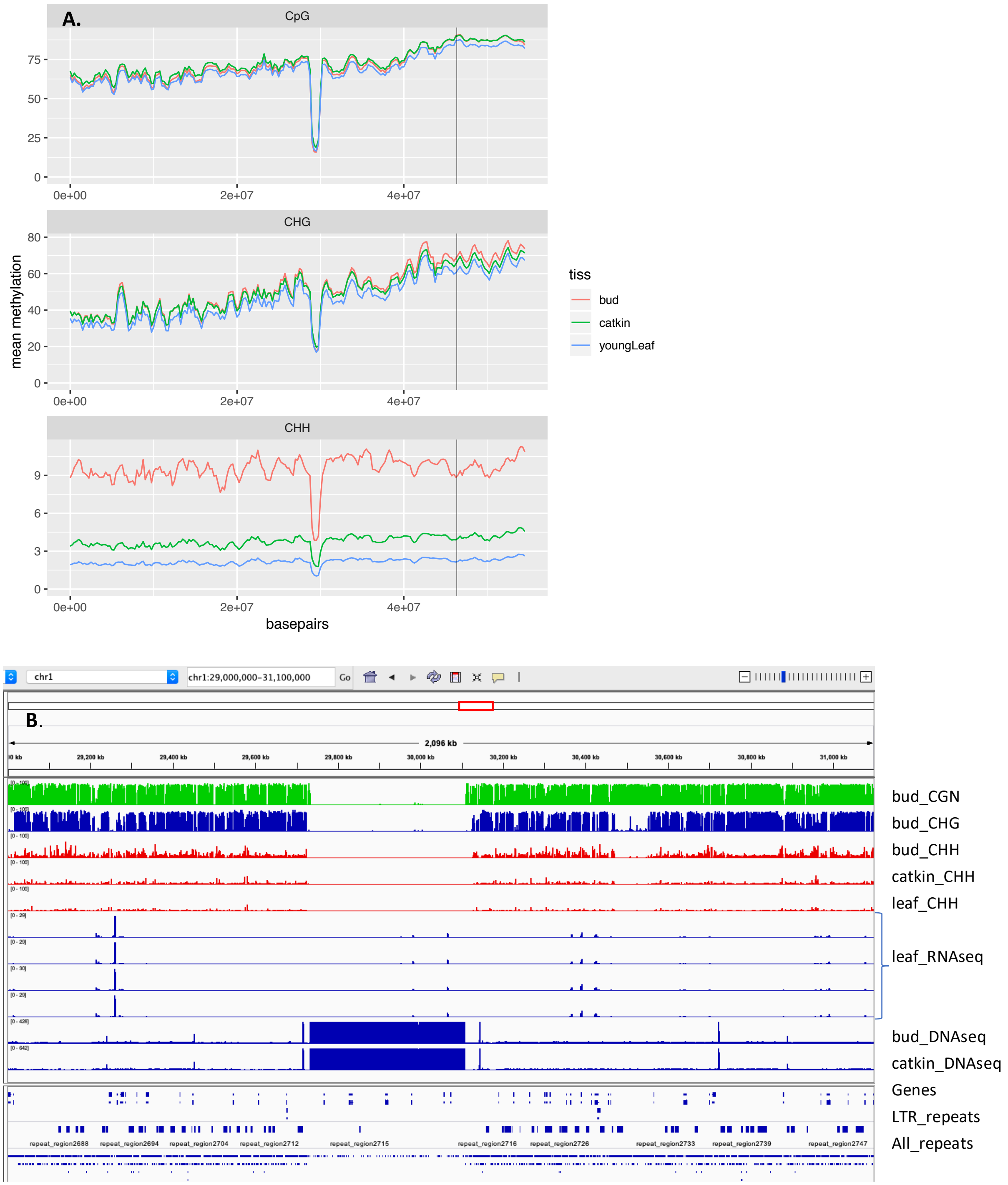

**Figure S3.** Mis-assembled mitochondrial sequence in pre-final *Q. lobata* version 3.0 chromosome 1. **(A)** Mean methylation level (CG top, CHG middle, CHH bottom) for 1 Mbp windows every 250 kbp. **(B)** IGV genome browser screenshot showing selected methylation levels (top five tracks), Illumina RNA-Seq read coverage (next four tracks), coverage by Illumina genomic reads (next two tracks), gene annotations (next track), and repeats (bottom two tracks). The strong dip in methylation levels and large increase in genomic read coverage are coincident with a mis-assembly that placed a region of the mitochondrion sequence into near-final chromosome 1 at one-based inclusive–inclusive coordinate span 29,726,880 to 30,108,053 bp (on the '+' strand). In the final 3.0 assembly release, this coordinate span has been replaced with gaps (to not shift coordinates at this late stage).

#### III. Analysis of heterozygosity

For comparison with the *Q. robur* genome [24], we analyzed the heterozygosity of our Q. *lobata* genomes in two different ways. The first way was to compute Tajima's $\pi$ [25] in non-overlapping 500 kbp windows across our 19 individuals. To do this, we used the Python function `windowed_diversity()` from the scikit-allel 1.2.1 package [26], with `window_size` set to 500 kbp. This function computes Tajima's $\pi$ by using allele frequencies of SNPs to compute the total number of pairwise differences across all samples. This number of total differences is then divided by the total number of callable sites in each window. Callable sites refers to the number of sites that passed our filters in each window (see **Input to PSMC'** below).

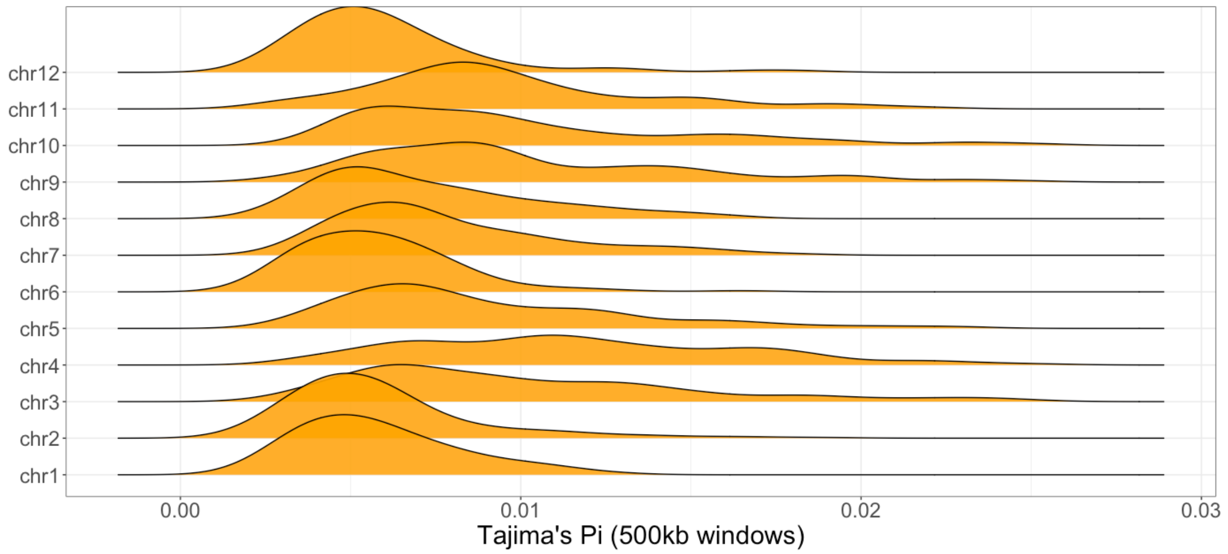

| Chromosome | Number Of 500kb Windows | Total Callable Sites | Total Number of Variants | Mean Tajima's Pi |
|---|---|---|---|---|
| 1 | 112 | 22,005,804 | 1,024,226 | 0.00580 |
| 2 | 209 | 41,297,702 | 1,746,992 | 0.00588 |
| 3 | 150 | 26,943,781 | 1,918,986 | 0.01004 |
| 4 | 196 | 30,807,490 | 2,422,621 | 0.01165 |
| 5 | 179 | 32,089,293 | 2,011,761 | 0.00895 |
| 6 | 109 | 22,882,763 | 1,074,710 | 0.00580 |
| 7 | 99 | 19,481,942 | 1,132,154 | 0.00780 |
| 8 | 131 | 26,525,174 | 1,307,858 | 0.00724 |
| 9 | 111 | 21,226,684 | 1,411,928 | 0.01000 |
| 10 | 133 | 23,245,880 | 1,600,529 | 0.00991 |
| 11 | 116 | 21,556,495 | 1,353,750 | 0.00954 |
| 12 | 87 | 19,006,813 | 847,498 | 0.00600 |

**Figure S4.** Distribution of Tajima's $\pi$ across the *Q. lobata* genome. **Top:** per-chromosome distribution of Tajima's $\pi$[25] across our 19 samples of diploid *Q. lobata* genomes. **Bottom:** per chromosome total number of 500 kbp windows, number of callable sites, and number of heterozygous positions for our samples.

The second way we summarized heterozygosity was by computing a heterozygosity rate. In this analysis, we examined each of the 19 diploid genomes independently. For each of the 19 genomes, we considered non-overlapping windows of 500 kbp. In each window, we counted the total number of heterozygous sites divided by the number of callable sites. This is the average number of heterozygous positions per callable base pair.

Both approaches gave similar results. Across chromosomes, both the heterozygosity rate and Tajima's $\pi$ had similar magnitudes and ranged from ~0 .005 to ~0.01. Likewise, both Tajima's $\pi$ and the heterozygosity rate have similar distributions within a single chromosome.

For our PSMC' analysis, we computed the heterozygosity rate of the *Q. robur* reference genome, the *Q. lobata* reference genome, and one resequenced *Q. lobata* genome (QL.LAY.5.00F). For the *Q. lobata* reference genome, we found 1,716,263 heterozygous positions out of 349,858,917 sites (~0.50%), and for the *Q. robur* reference genome, we found 2,268,413 heterozygous positions out of 309,542,806 sites (~0.73%). For the *Q. lobata* resequenced genome, we limited our analysis to filtered sites in QL.LAY.5.00F shared by both QL.LAY.5.00F.2018 and the reference genome, and found 2,025,194 heterozygous positions out of 307,071,743 sites (~0.66%).

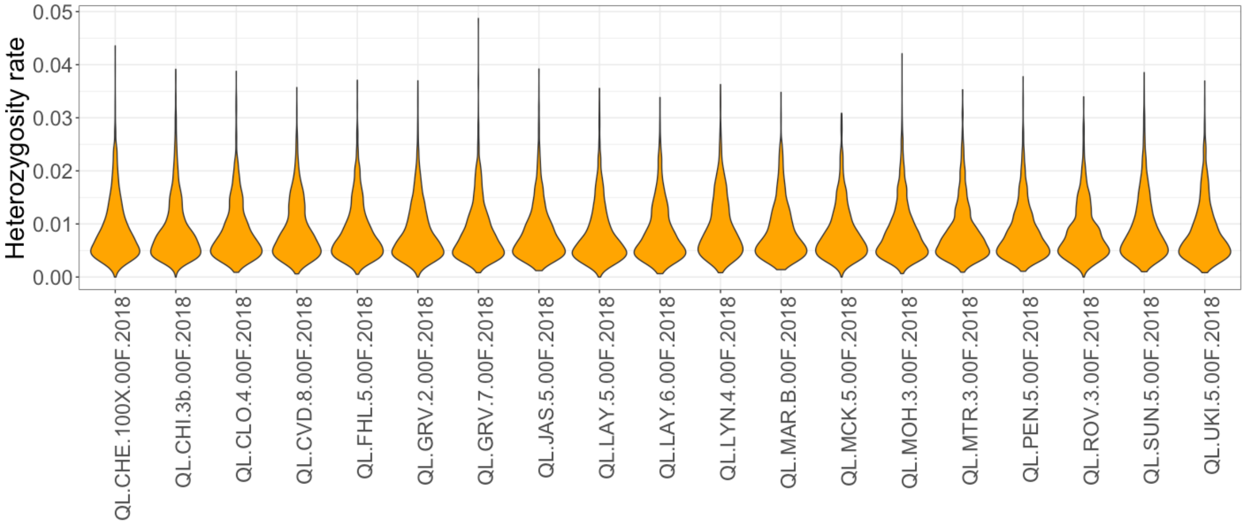

| Chromosome | Number Of Windows | Total Callable Sites | Number of Heterozygous Positions | Heterozygosity |
|---|---|---|---|---|
| chr1 | 2,128.00 | 317,717,076.00 | 1,925,442.00 | 0.006060 |
| chr2 | 3,971.00 | 585,250,875.00 | 3,355,901.00 | 0.005734 |
| chr3 | 2,850.00 | 345,332,020.00 | 3,149,517.00 | 0.009120 |
| chr4 | 3,724.00 | 381,396,603.00 | 4,295,624.00 | 0.011263 |
| chr5 | 3,401.00 | 418,965,586.00 | 3,617,755.00 | 0.008635 |
| chr6 | 2,071.00 | 331,233,288.00 | 1,925,282.00 | 0.005812 |
| chr7 | 1,881.00 | 270,412,151.00 | 2,120,958.00 | 0.007843 |
| chr8 | 2,489.00 | 373,392,027.00 | 2,652,599.00 | 0.007104 |
| chr9 | 2,102.00 | 285,985,560.00 | 2,695,443.00 | 0.009425 |
| chr10 | 2,527.00 | 303,504,786.00 | 2,694,611.00 | 0.008878 |
| chr11 | 2,204.00 | 290,439,119.00 | 2,611,644.00 | 0.008992 |
| chr12 | 1,653.00 | 272,784,844.00 | 1,602,538.00 | 0.005875 |

**Figure S5.** Distribution of heterozygosity rate (heterozygosity per bp) across the *Q. lobata* genome.
**Top:** distribution of heterozygosity rate across our 19 samples. **Bottom:** per chromosome total number of 500 kbp windows, callable sites, and number of heterozygous positions in our samples.

#### IV. Demographic analysis

##### Methods of analyses

**Inference of demographic history.** We used the Pairwise Sequentially Markovian Coalescent (PSMC′) model to infer changes in effective population size in *Q. lobata* and *Q. robur* over time [27]. With a single diploid genome, PSMC′ utilizes the spatial distribution of heterozygous sites to first infer a distribution of times to the most recent common ancestor (TMRCA) across a whole genome. With this distribution of TMRCAs, PSMC′ can then estimate the effective population size $N_e$ across the evolutionary history of a population using the inverse relationship between the coalescence rate and the effective population size [27]. Although the PSMC model was first developed to study the demographic history of humans [27], it has been used in the study of animals with distinct phylogenetic histories [28-30] as well as a variety of plants [31-35].
**Input to PSMC′.** For our analysis with PSMC′ we first masked out all genome gaps and repeats from the *Q. lobata* reference and resequenced genomes, and the *Q. robur* reference genome. Additionally, insertions and deletions were masked out. For the reference genomes, the mean depth (DP) for variants was 110 reads and the standard deviation of DP was 60. We set the maximum DP filter for the non-variant sites in the reference genomes to be the mean DP + 4*(standard deviation) which is 350, and the minimum DP was set to be the mean DP divided by 3 (110/3=37). Variant sites in the reference genomes that satisfied any of our filter conditions (DP > 350, FS > 60, MQ < 40, QD < 2, SOR > 3, RPRS < -8, MQRankSum < -12.5) were also excluded from analysis

To ensure that the demographic history obtained for *Q. lobata* was not biased by mapping its sequencing reads back to its own assembled genome, we also ran PSMC′ on 19 additional resequenced *Q. lobata* genomes (Table S1). We generated PSMC′ input using only callable sites, which we define as having a minimum depth (DP) of greater than 12 reads with a mapping quality (MQ) greater than 20, and a quality (QUAL) score greater than 10. The mean coverage of callable sites for all 19 resequenced samples was greater than the recommended mean genome coverage of ≥ 18x [36] in all but 5 samples. These 5 samples had mean coverages that spanned 14.89x to 17.58x. Additionally, we removed all indels, variant sites immediately upstream and downstream of insertions and deletions, multiallelic sites, and repetitive sequences. 56.7% of the genome was removed due to repeat masking, which is greater than the ≤ 25% missing data threshold recommended by Nadachowska-Brzyska, et al. [36]. However, because of the overwhelming presence of transposable elements and repetitive sequences in *Q. lobata*, we masked out these sequences to avoid incorporating incorrectly called SNPs that are the result of mapping errors. PSMC′ was run with default settings except for the maximum number of iterations being set to 200. Because PSMC′ was designed to be used on human genomes, it begins its expectation-maximization algorithm to infer the ratio of recombination and mutation rates at a value of 0.25. Although starting at this ratio of 0.25 might be appropriate for humans, it is currently unclear how the coupling of long lifespan [37] and non-human reproductive biology (for example possible somatic generation of diversity being passed onto the next generation[24]) of oaks contributes to this ratio in *Q. lobata*. By allowing for more iterations of the expectation-maximization algorithm, we allow for a larger space of recombination to mutation rate ratios to be explored. Qualitatively we did not see large differences in the demographic trajectory depending on the number of maximum iterations set except in the ancient time steps.

**Estimation of neutral mutation rate.** Neutral mutation rates for *Q. lobata* and *Q. robur* are needed to scale PSMC′ output into the units of years ago and effective population size. Unfortunately, published estimates of these quantities are not available. Thus, we estimated the neutral mutation rate from sequence divergence. Assuming the divergence between *Q. lobata* and *Q. robur* is much greater than the expected levels of polymorphism in the ancestral species we estimated a mutation rate using the relationship between divergence and split time [38]. To compute a mutation rate, we used MUMMER [39] to align the *Q. lobata* version 3.0 reference genome and the *Q. robur* reference genome to each other). We then calculated divergence by counting the number of positions that differ between the aligned reference genomes that have a 1 to 1 mapping and divided this number by the total number of aligned nucleotides. In this computation, we masked out repeats and genome gaps in both genomes and found 241,827,461 matching nucleotides between *Q. robur* and *Q. lobata* and 4,555,467 mismatching nucleotides. Then, using an estimated split time of 35 million years and a generation time for *Q. robur* of 30 years and *Q. lobata* of 50 years we estimated a mutation rate of $1.01 \times 10^{-8}$ bp/gen. The generation time for *Q. robur* was based on estimates of other temperate tree species, such walnut [35], and the generation time for *Q. lobata* was set at 50 years because maximum life span of Q. lobata is greater (1000 years versus 600-800 years) and other life history traits such as larger acorn crop sizes and ages of standing tree populations are older for *Q. lobata* than *Q. robur*.

Accurate estimates of mutation rates are difficult to experimentally measure in woody plants [40]. Additionally, it is difficult to estimate accurate neutral mutation rates for these organisms with sequence divergence. It is possible that our neutral mutation estimates are inaccurate due to factors that are not constant over time such as differences in DNA-repair mechanisms, generation times, metabolic rates, in our inability to incorporate uncertainty in fossil identification, uncertainty in estimates of fossil ages, and the large variance around the substitution rate for any given time period [41]. However, different estimates of the mutation rate and generation time scale the axes but do not change the overall shape and pattern of the inferred effective population size trajectory (see Figure S6). Therefore, our qualitative conclusions about the demographic history of *Q. lobata* should be relatively unaffected by these possible biases.

**Simulations in msprime.** To qualitatively assess whether PSMC' can accurately infer population size changes similar to those for oak trees, we used coalescent simulations implemented in msprime to simulate data under our inferred demographic models for each of the three types of genomes. For the *Q. lobata* reference genome, *Q. robur* reference genome, and the *Q. lobata* resequenced genome, the inferred demographic history outputted by PSMC' is defined by 40 points. We scaled these 40 points into effective population size ($N_e$) and the number of generations before the present (γ) using the mutation rate of 1.01 x $10^{-8}$ bp/gen (see section above, **Estimation of neutral mutation rate**) and the following formulas:

$$\gamma = \psi/\mu$$
$$N_e = (1/\lambda)/(2 * \mu)$$

where:

γ = Number of generations before the present

$N_e$ = Effective population size

μ = Neutral mutation rate in bp per generations

ψ = PSMC' inferred left time boundary

λ = PSMC' inferred Lambda_00

With 40 pairs of $N_e$ and γ, we generated a corresponding msprime function. Each change in $N_e$ was done instantaneously with a growth rate of 0. To generate one replicate of a simulated genome, we simulated 12 independent replicates of chromosomes of length 29 Mbp. The recombination rate for our simulations was set to be uniform across each simulated chromosome with a rate of 2 x $10^{-8}$ bp/gen. The mutation rate for our simulations was also set to be uniform across each simulated chromosome with a rate of 1.01 x $10^{-8}$ bp/gen. After each simulation completed, we used msprime to output each simulated diploid chromosome as a VCF. We then generated the input to PSMC' (a "multihetsep" file) from a single VCF using a custom script that can be found at github.com/jessegarcia562/psmc2msprime. With 12 simulated chromosomes and their corresponding multihetsep files, we then utilized PSMC' with the default settings and 200 iterations to infer the demographic history of one simulated genome.

In this paper, we studied three genome types: *Q. lobata* reference genome, *Q. lobata* resequenced genome, and *Q. robur* reference genome. For each genome type, we performed the described simulation and PSMC' inference 10 times. These analyses left us with 10 simulated genomes for each genome type and therefore 10 PSMC' inferred demographic histories for each genome type. For each of the 30 genomes, we computed heterozygosity by dividing the total number of heterozygous sites in the simulated genome by the total simulated genome length (12 simulated chromosomes * 29Mbp per chromosome = 348 Mbp).

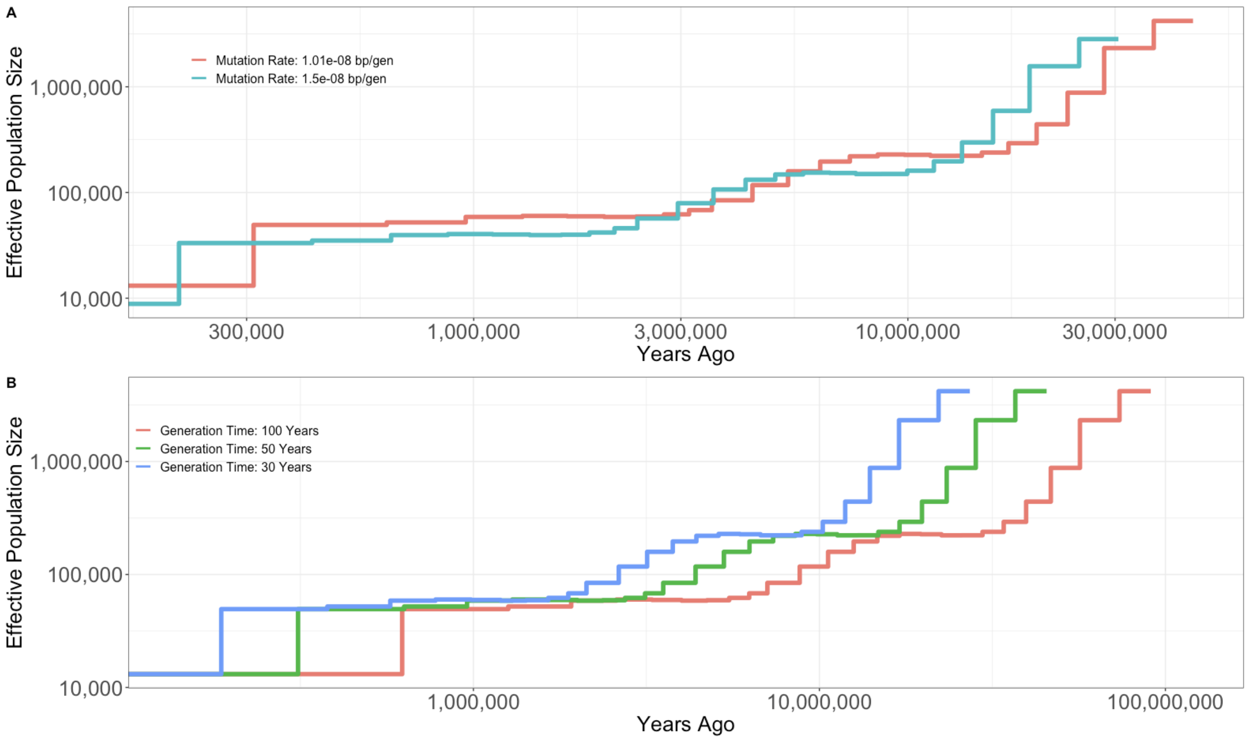

**Figure S6.** PSMC' inference on the *Q. lobata* reference genome when using different generation times and mutation rates. **(A)** Assuming a generation time of 50 years, different estimates of the mutation rate would move the demographic trajectory along both the *y*-axis and *x*-axis. However, different estimates would not change the overall shape of the curve. The mutation rate $1.01{\cdot}10^{-8}$ bp per generation was estimated from the divergence between the *Q. lobata* and *Q. robur* reference genomes (see **Estimation of neutral mutation rate** above). The $1.5{\cdot}10^{-8}$ bp per generation mutation rate illustrated here was chosen arbitrarily to illustrate the effect changing mutation rate has on effective population size. (**B**) Assuming a mutation rate of $1.01{\cdot}10^{-8}$ bp per generation, different estimates of generation time would only move the demographic trajectory along the *x*-axis, as larger generation times would push the estimates farther into the past.

**Identifying a trim point.** While PSMC' can infer complex population size-change models, these models may not accurately predict simple empirical summary statistics such as the genome-wide distribution of heterozygosity (Beichman, Phung, and Lohmueller (2017)). Although it is unclear precisely why this occurs, one hypothesis is that methods such as PSMC' might overestimate the ancestral size of a population (Beichman, Phung, and Lohmueller (2017)). In order to present a demographic history that accurately predicts both the empirical genome-wide rate of heterozygosity and the empirical genome-wide distribution of TMRCA, we attempted to correct for the possible overestimation of the ancestral size from the initial (full model) (Figure S8). PSMC' inference. Because the demographic trajectories for each genome type all appeared to be monotonically decreasing (moving forward in time) in our ancient time steps, we decided to use each predicted time step as a possible ancient ancestral population size. Specifically, we had in total 40 inferred pairs of $N_e$ and γ that defined the demographic trajectory for each genome type. From the original 40 pairs of points PSMC' inferred, we created 39 new demographic trajectories by iteratively removing the most ancient (largest in magnitude γ) time step. For example, while 40 points describe the full PSMC' demographic model, after removing the most ancient time step, we can generate a new demographic trajectory that is instead defined by only 39 points. This iterative process of generating new demographic trajectories results in one full model (all 40 pairs of $N_e$ and γ) inferred by PSMC' and 39 trimmed models (a model defined by 39 points, a model defined by 38 points. . . a model defined by 1 point). Importantly, the population size remains at the same size as the last point defining the demographic history for an infinite amount of time going back into the past. Thus, this trimming strategy resulted in changing the ancestral population sizes of the PSMC' inferred demographic model.

Following the methods described in **Simulations in msprime,** we simulated 1Mb of sequence under each of our 40 models for each genome type and computed the predicted heterozygosity of each model (Figure S8). We then visually compared the fit of the simulated distribution of heterozygosity to the values observed empirically. Our best models for *Q. lobata* reference genome, *Q. robur* reference genome and *Q. lobata* resequenced genome were defined by 32, 32, and 28 points respectively, although none of our 40 models could precisely predict the exact genome-wide heterozygosity for the respective genome type. This suggests that the true demographic history is likely more complex and is not entirely captured with these size change models. Nevertheless, trimming allows the demographic models presented in Figure 2 to more closely match the heterozygosity in the observed data than what the untrimmed model predicted (Figures S9 & S10).

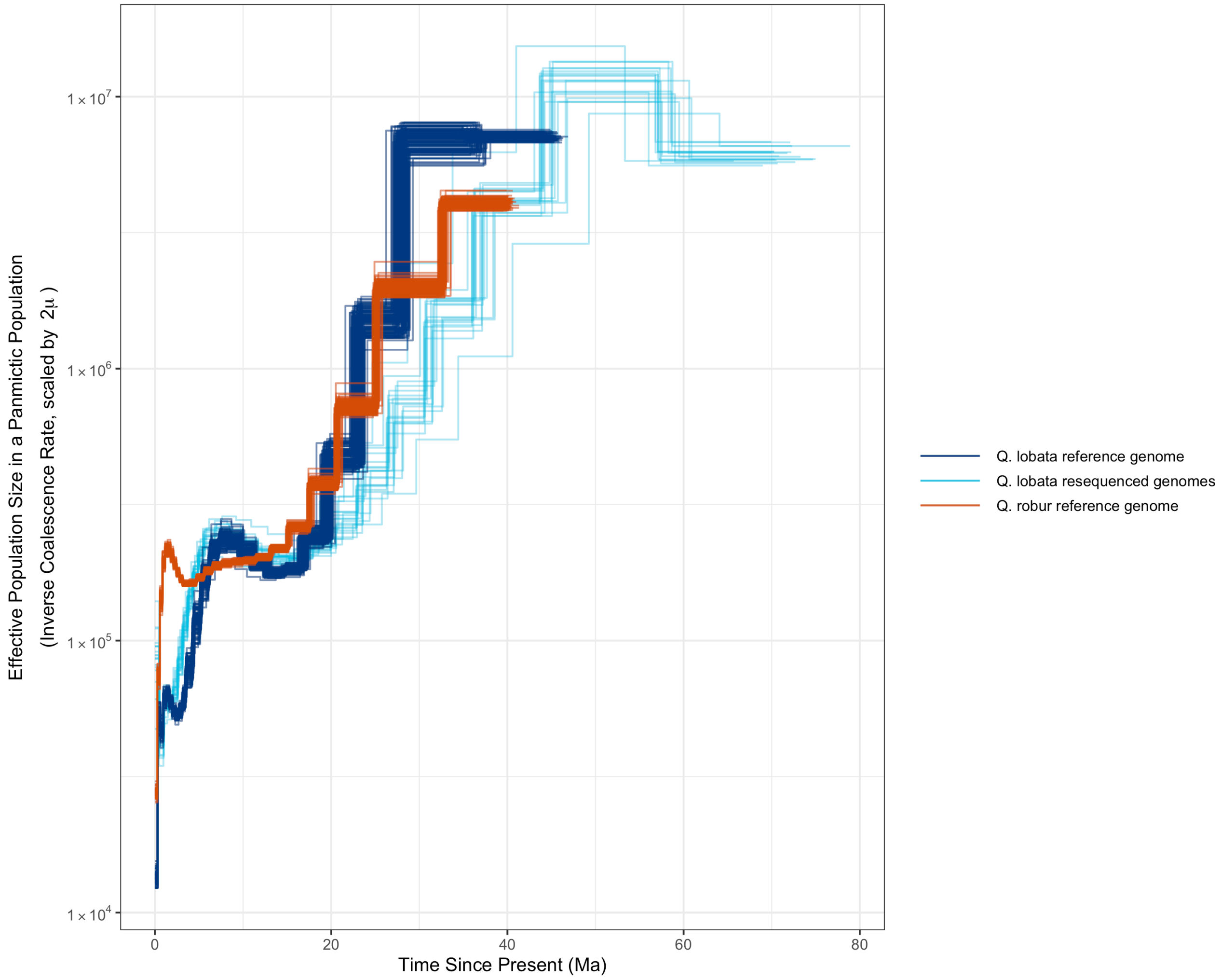

**Figure S7.** Full demographic models inferred by PSMC′. This figure differs from main text Figure 2C in that none of these models have their ancestral population sizes trimmed to fit the empirical rate of heterozygosity observed; visualized here is the unprocessed raw output from PSMC′ scaled by the estimated mutation rate and generation time for each species.

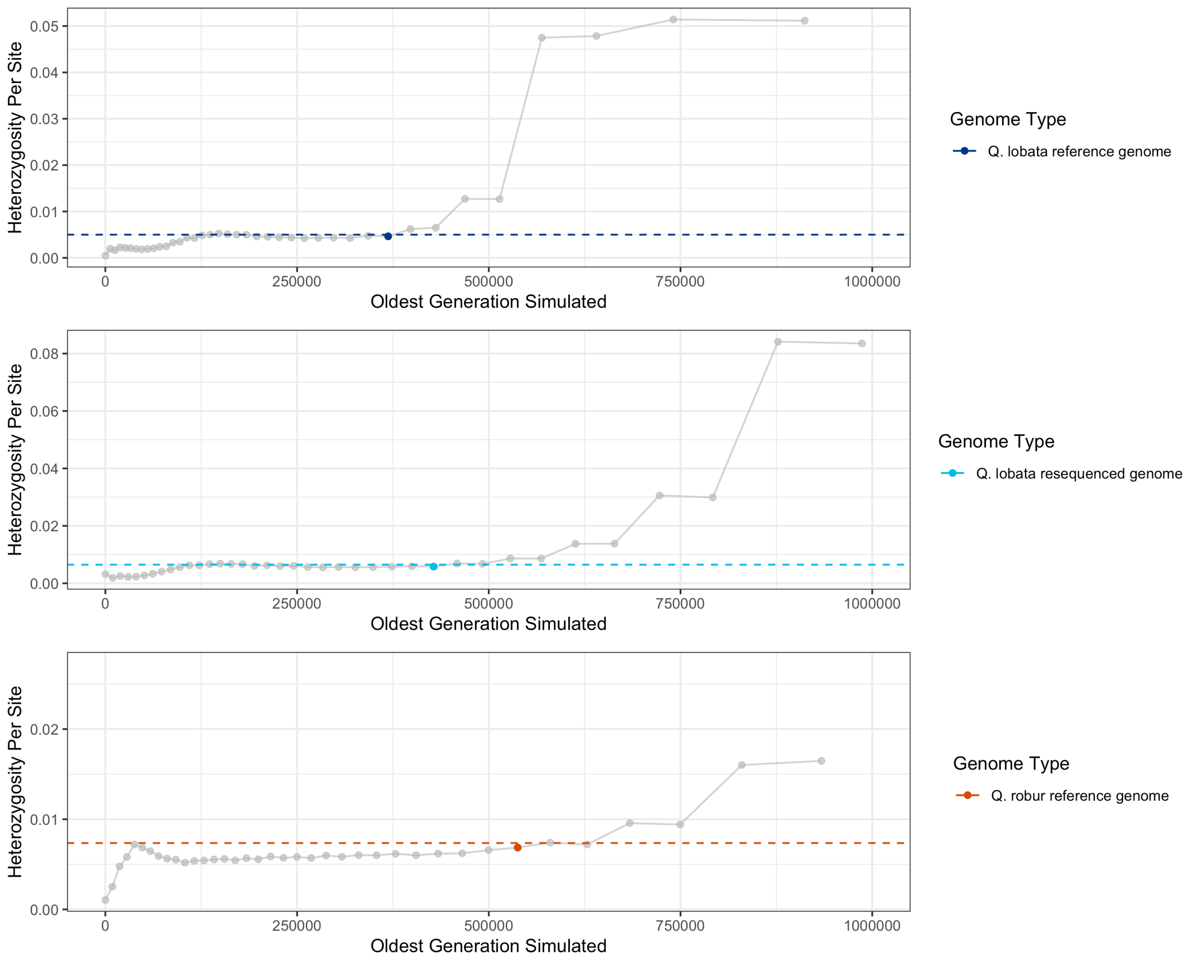

**Figure S8.** Predicted heterozygosity for 1 Mbp regions for all 40 models for each genome type. Dashed lines are the respective empirical heterozygosity rate for each type. Highlighted points represent the models that we chose to represent each genome: we chose Model 32 for both the *Q. lobata* and *Q. robur* reference genomes, and Model 28 for the *Q. lobata* resequenced genome.

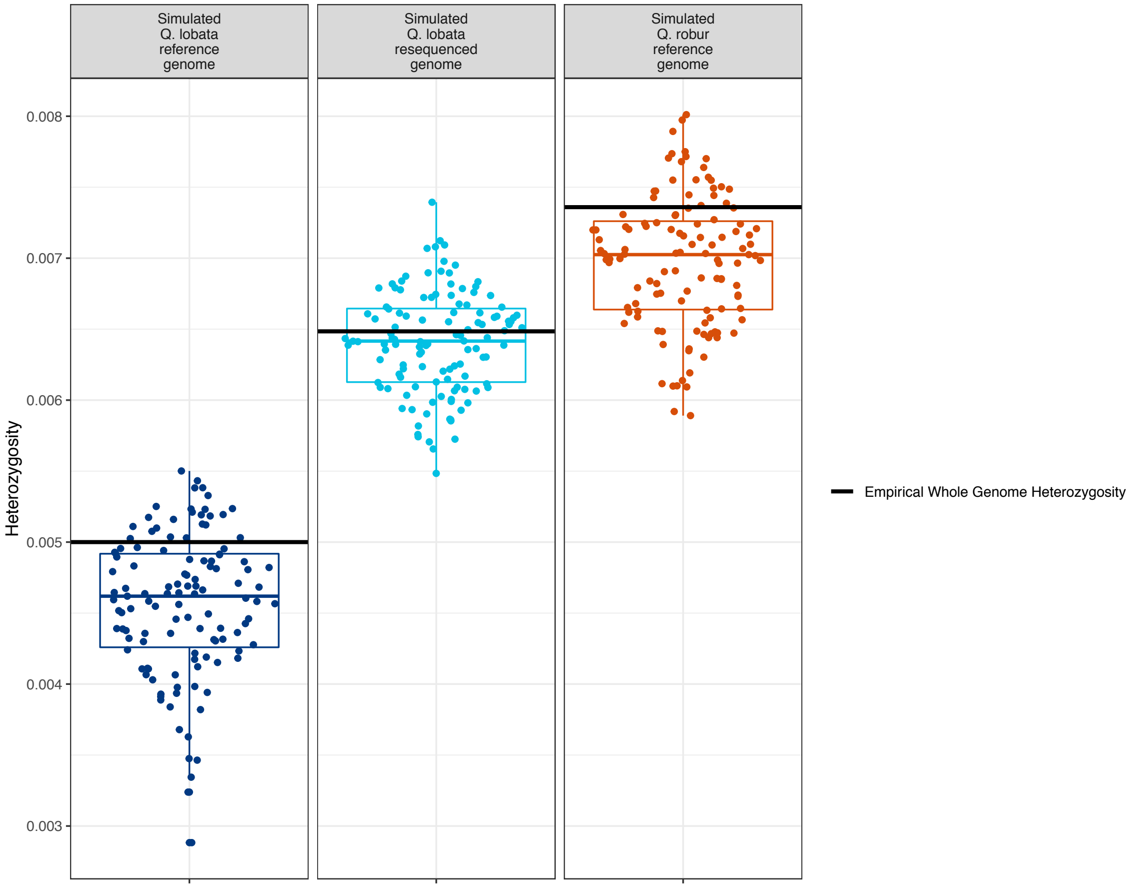

**Figure S9.** Predicted heterozygosity for 120 simulated 1 Mbp regions of the best fitting models for each genome type.

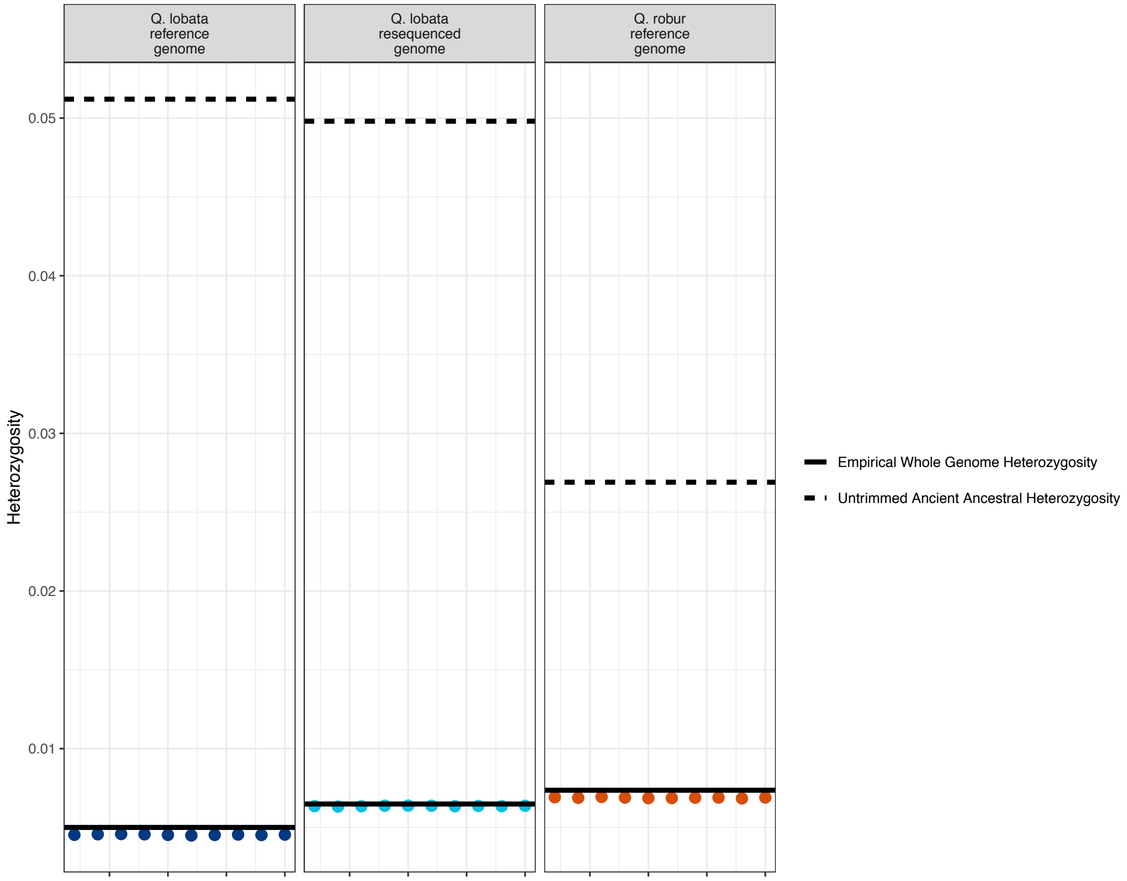

**Figure S10.** Predicted heterozygosity of full untrimmed PSMC' models compared to the predicted heterozygosity of best fitting trimmed ancestral models. Best fitting trimmed models are models which have reduced ancestral population sizes (relative to the full untrimmed PSMC' models), and when simulated with msprime fit empirical whole genome heterozygosity. Models 32, 28, and 32 were chosen for the *Q. lobata* reference genome, the *Q. lobata* resequenced genomes, and the *Q. robur* reference genome, respectively. Each point is the heterozygosity of a simulated genome under our trimmed ancestral model.

#### V. Repetitive sequences

##### *Additional findings and methods*

A database generated by RepeatModeler consists of repeat “families”, each given by a consensus nucleotide sequence derived from a multiple alignment of some high-copy homologous regions of the genome, and many families are automatically placed into a particular major (e.g., “Long Terminal Repeat” — LTR) and minor class (e.g., “Gypsy”) by a subcomponent (RepeatClassifier) aligning against known repeats (with variable accuracy; we did not revise by manual curation). The consensuses are not always full length for their class or irredundant by close sequence similarity; for *Q. lobata*, we applied PSI-CD-HIT 4.7 to cluster at 45% nucleotide identity (the level where, as the threshold is lowered, intracluster similarities stop falling in frequency and begin rising) and chose a canonical rotation/strand for tandem repeat units so as to cluster families into “superfamilies” (SFs), generally assigning to each SF the major/minor class of the longest member family that was not unknown (if any). Annotated intervals for a SF are the nucleotide-level union of all intervals for its member families, and SFs are given “s1RF####” accession numbers (roughly by descending mass of nucleotides masked). We also applied LTRharvest and LTRdigest from GenomeTools 1.5.9 to specifically target the prevalent LTRs, which identified 28K instances covering a total of 184 Mbp (only slightly more than the 179 Mbp RepeatClassifier declared as LTR).

Examination of instances of individual SFs identified s1RF1096 as the telomeric tandem repeat (the common plant unit $(AAACCCT)_n$ when at 5′ ends, and mostly restricted in occurrence to edges of assembled chromosomes), as well as 148 bp complex tandem unit s1RF0004 (GCTCATGGGC CCCCGACCCG AGTTAGAAAA TTCAAAAAAT AAATGCAAAA AAATTCTAAA AATTAAAAAA CATCATCCAG GCTTCATTTC AAGACGAAAA CGGGTCAGAG ACAGGCCGAA AAATAGAGAA CAAAAATTTC ATTCCTAA) which exists in relatively large total quantity (≈3 Mbp) and is essentially restricted to exactly one locus per chromosome, strongly suggesting this identifies centromeres, with s1RF0004 reminiscent of, e.g., CEN180 of *Arabidopsis* [42]; over the project, further evidence (gene density profiles and DNA methylation patterns) accumulated additional support. Approximate intervals spanning centromeres are given in the table below. Clustering of chromosomal distributions of SFs (Figure S11) indicated that the main chromosome-scale distributional features of repeats are associated with distance to centromeres. The distributions are well-summarized per SF by average distance of the SF members to the centromeres (Figure S12), and were used to identify SFs with unusual preference for or avoidance of the centromeres (Figure 3C–D). The SFs so-identified have striking distributional concentrations that are nearly completely diluted away if only the distribution of all repeats taken together is examined (which is nearly uniform across chromosomes). These concentrations mainly fall to individual SFs and are not strongly associated with entire major repeat classes. Exonic density from protein-coding genes also shows a notable gradient, being lower near centromeres.

| Chromosome | From | To |
|---|---|---|
| chr1 | 45,377,794 | 47,303,741 |
| chr2 | 42,723,096 | 45,702,847 |
| chr3 | 46,505,435 | 47,582,132 |
| chr4 | 57,272,007 | 58,869,737 |
| chr5 | 37,258,031 | 38,038,545 |
| chr6 | 25,012,718 | 26,574,917 |
| chr7 | 18,052,972 | 19,452,171 |
| chr8 | 24,326,695 | 25,399,835 |
| chr9 | 36,750,921 | 36,955,183 |
| chr10 | 36,598,025 | 38,256,999 |
| chr11 | 30,653,296 | 31,212,808 |
| chr12 | 19,137,856 | 20,625,243 |

(1-based inclusive–inclusive intervals on ‘+’ strands for approximate intervals that contain centromeres)

Identification of large arrays of rDNA was attempted. *In silico* isolation of a canonical rDNA tandem unit* was complicated by the unit’s borders incorporating a complex multi-scale tandem repeat (with $(GGCCTT)_n$ as short bottom-level unit), with individual copies of the rDNA unit highly diverging there. Alignments of Illumina short reads to a 9.1 kbp consensus (that included full 18S+28S) vs. generic homo- and heterozygous regions of the nuclear genome suggest a total of ≈8 Mbp of rDNA from ≈870x copies of the rDNA unit, similar to the ≈750x of *Arabidopsis* chromosomes 2+4 and the ≈840x of *Chlamydomonas* chromosomes 1+7+15 [43]. The clean final assembly, however, contains only ≈0.3 Mbp aligning to the 9.1 kbp consensus. Thus (as is common), the assembly highly under-represents rDNA. The highest concentration found is on chromosome 1 (the only acrocentric chromosome) in the short right arm (where LG1 is constant in a large block, a phenomenon that occurs only once across all LGs).

*Best-guess canonical rDNA tandem unit — starts with a 2,943 bp spacer, with first ≈1.6 kbp repetitive; NCBI web BLASTN finds *Quercus robur* EF208967.1 for this, and in fact a note for that says "derived from 8Kb rDNA unit":

```
GGCCTTGGCTGCGTGCCTTGGCCTTGGCAGCATGCCTTGGCCTTGGCTGCCTTGGCCTTGGCTACGTGCCTTGGCCTTGGCAGCATGCCTTGGGGGCCTGCCTTGGCCTTGGCTGCATGCCTTGGCCTTGGCTGCATGGGCCTTGGCTGCGTGCCATGGGGGGCAGCCCC
TGGCCTTGGCAGCATGCATGCCATGGCCTTGGGCCTTGGCCGCGTGCCTTGGCCTTGGCTGCTGCCTTGCCCACAAATTTCGAGTGATTTCCCAATAAATGAGGGTTTTTTGGAAAAGGAGGTTATTTTCCCCCCAAAGAGGCAGGCGTTGGGCATGGCAGGGTGCCCAG
GGGCATGCCCGCATGCACGCCACGCCGCATCGCCCCGCATCGTCCCGCACTCCGGCAAAATTGGCTCGTGCCTTTTCCCCTTTTTGTGTTCCTAAATTCAGTCCATGATTTTAGAGGACGTTTCCAACAAGCGGTTCGGCGTTCCGAGCAGTTTTGAATTTTTTATGATT
TTTCCTATTTTCTGGATTCCGAAAATCATAAAAAATAAAATATGTTGAATCTGGCCACCAAATTTTGACAGGTTGAAGGTTAGATTTTTCTTAGCATGTGTACAAAAAATCAGGGCAAGACTCCAAGAATTGCTCCAAAAAAGACCCATTATTCCTCCTCAACAAATCAA
TGTTTCCTCGTGCCAGTGTGGATTTTTTCCTAAGCGCTCTTTAGGGGGGGAAGAGTTGGAGGTGTCCGAAGAGGCTATCTAGGCTTGGCAGCAGTAGCCCCCCCAAGTGCTGCCACGGCCTTGTAGGGGTCCCAAGGGCATGCCACGGCCTTGGCATCCCACCCACGGGG
CATGCCACGCCCTCGCCCACGGGGCATGCTAGGGCCTTGCTGCTGCCTGCGCCCAGGGGCATGCCAGCGTGCCCACCTGCCTGCACCCAGGGGCATGCCAGCGTGCCCACGCAAGCATGCCATGGCCTTGGCTGCGTGCCTTGGCCTTGGCAGCATGCACGCAAGCAAGC
ATGCATGCCTTGGCCTTCGCTGCCTGCCCATGGGGGCTGCCTTGGCCTTGGCTGCTGCATGCCCATAGGGGGCTGCCTTGGCCTTGGCTGCATGCCTTGGCCTTGGCCTTGGATGCATGCCTTGCCCATGGCCTTGGCCACATATTTGGAGTGATTTCCTAAAAATGAGG
GTTTTTTGGAAAATGAGGTTATTTCCCCACGAGCAGGCAGGCAGTGGGCATCCCAGGGGCATGCCCGCGGGCACGCCGTGCCGCATCGCCCCGCATCGTCCCGCACTCCGGAAAAATTGGCTCGTGCCTTTTTCCATTTTTGTGTCCCTAAATTCATTCCATGATTTTAG
AGGAGGATTCCAGCAAGCGGTTCGGCGTTCCGAGCGTTTTTGAATTTTTTATGATTTTTCCCATTTTCCGGCTTCCTAAAATCGTAAAAAATAAAATATGTTGAATCTGGCCTCCATATTTTGACAGGTTGAAGGTTGGATTTTTCTTAGCATGTGTGCAAAAAATCAGG
GCAAGACTCCAAGAATTGCTCCGAAAATGACCCATTATTCCTCCTCAACAAATCAATGTTTCCTCGTGCCAGAGCGGATTTTTTCCTAAGGGCTCTTTAGGGGGGAGGAGGTATTCGGCGATGCACAGGGGCGACCCCTCTTGAGCTTGGCCACGGGTAGGCATGCTCAT
GACGAGCCCTCAGGCACCCAACCCCGCGTGCTCCATGGCGCGAGGTGGGCCCTAAATGCATCGCCGGGGTGGACCCAGGTTGGCATTTCACGTGTACGTGGTATAGCTGCGGCGCGCTCTTGCAAGGCGGACGGTGTTAGTTTCACCCATTGCCACGCAGAGCAAGCCTA
AGGTGATGATCAAACGCATTGTGAAAGCTTTCTCTGGTGCCACTTTTTCCTCGAGGCCACACCCACCCGTGCCCAGTGGCGGGGGGTCGATGGTCTCAGTAGCCGCGTGGTGGGGGGATGCATGCATTCTTGGTTTAGCTTGGTCACGCTATCGCAACGCGGACGGTGTT
AGTTTCACCCATTGCTGCGCGAAGCAAGTTCCATGCGACTATCGGGCTTGTGTGTGTCTTTCTTACTACGCGGTTGCGTGGTAGCAGTGGCGGCGGCGGCGGCGGCGGCGACGGCGACGACGGCAACGACAGCAGTGTCCCTCGATGTCCCTCGATGTCCCTTGTAGTTC
CTGTGTGTGGTTGGTGAGCCATGCAAGCTTGGTATAGCTTGGGCACGCTCTCGCAAAGCGGACGGTGTTCGTTTCACCCATTGCTGCGCGAAGCAAGTCCCACGGCGACCATCGGGCCTCTGCGCGGCGACGACCCATCCTTGCAGGCACGTGGCTTAGTAGTGTTGTCC
TTCACGCCCAGCGGTGCGGACTCGGCGCCACTCGATGCTTCCTCATGCACGGCAGGCCTTGCGGCGTGTCGTTGTGGGGTACGTTTGGTGGTGTTGCGGTCTAGTGTACGTGATAGCGTGTGAGTGGTGGCAGGGTTGCATGGCTTGGCAGGCTCCGTGCTCGCGCATCG
AACTGTCCGGCGTGCTCCCAATCAGCGTTGTTCCTAGCGTCGCTCGGACGCAATTCGGGTCCCTGTGTTGCATACCTGCCTCTAAGGCACTCGTCCCTCTAGTTGATTCGTTCCTAGTCGACGCTCCTCACGGGGCGTCGGCAGGACCTCGAAGCCGTCCTCGTGTCCCA
CGCGTGCCTCGCGGCCTCCGCGTTGCCGATGTGGACCACGTGGGCGTGCTCGTGGCCTCGGATGCAGAACACCATGTGGGTTTGGGGCCTTCGGCCCCCTTTGCCAACGTACCTAGCGAGCGTCATCGCTCTGCCCCGCACGATCGCCGTGCTTGTCCGTGCCCTTCCTT
GCCCTCGGGCGAGCCAGGGCCTCCGGGCGGCGCCGGCATCGACGAGGAATGCT
```

Next is 1,808 bp 18S (per RNAmmer):

```
ACCTGGTTGATCCTGCCAGTAGTCATATGCTTGTCTCAAAGATTAAGCCATGCATGTGTAAGTATGAACTAATTCAGACTGTGAAACTGCGAATGGCTCATTAAATCAGTTATAGTTTGTTTGATGGTACCTGCTACTCGGATAACCGTAGTAATTCTAGAGCTAATACG
TGCAACAAACCCCGACTTCTGGAAGGGATGCATTTATTAGATAAAAGGCCGACGCGGGCTCTGCTCGCTGCTCTGCTGATTCATGATAACTCGACGGATCGCACGGCCATCGTGCCGGCGACGCATCATTCAAATTTCTGCCCTATCAACTTTCGATGGTAGGATAGTGG
CCTACTATGGTGGTGACGGGTGACGGAGAATTAGGGTTCGATTCCGGAGAGGGAGCCTGAGAAACGGCTACCACATCCAAGGAAGGCAGCAGGCGCGCAAATTACCCAATCCTGACACGGGGAGGTAGTGACAATAAATAACAATACCGGGCTCTCACGAGTCTGGTAAT
TGGAATGAGTACAATCTAAATCCCTTAACGAGGATCCATTGGAGGGCAAGTCTGGTGCCAGCAGCCGCGGTAATTCCAGCTCCAATAGCGTATATTTAAGTTGTTGCAGTTAAAAAGCTCGTAGTTGAACCTTGGGTTGGGCAGAGCGGTCCGCCCCTGGTGTGCACCGG
TCTGCTCGTCCCTTCTACCGGCGATGCGCTCCTGGCCTTAACTGGCCGGGTCGTGCCTCCGGTGCTGTTACTTTGAAGAAATTAGAGTGCTCAAAGCAAGCCTACGCTCTGGATACATTAGCATGGGATAACATCATAGGATTTCGGTCCTATTCTGTTGGCCTTCGGGA
TCGGAGTAATGATTAACAGGGACAGTCGGGGGCATTCGTATTTCATAGTCAGAGGTGAAATTCTTGGATTTATGAAAGACGAACAACTGCGAAAGCATTTGCCAAGGATGTTTTCATTAATCAAGAACGAAAGTTGGGGGCTCGAAGACGATCAGATACCGTCCTAGTCT
CAACCATAAACGATGCCGACCAGGGATCGGCGGATGTTACTTATAGGACTCCGCCGGCACCTTATGAGAAATCAAAGTCTTTGGGTTCCGGGGGGAGTATGGTCGCAAGGCTGAAACTTAAAGGAATTGACGGAAGGGCACCACCAGGAGTGGAGCCTGCGGCTTAATTT
GACTCAACACGGGGAAACTTACCAGGTCCAGACATAGTAAGGATTGACAGACTGAGAGCTCTTTCTTGATTCTATGGGTGGTGGTGCATGGCCGTTCTTAGTTGGTGGAGTGATTTGTCTGGTTAATTCCGTTAACGAACGAGACCTCAGCCTGCTAACTAGCTATGCGG
AGGTGACCCTCCGCGGCCAGCTTCTTAGAGGGACTATGGCCGCTTAGGCCAAGGAAGTTTGAGGCAATAACAGGTCTGTGATGCCCTTAGATGTTCTGGGCCGCACGCGCGCTACACTGATGTATTCAACGAGTTTATAGCCTTGGCCGACAGGCCCGGGTAATCTTTGA
AATTTCATCGTGATGGGGATAGATCATTGCAATTGTTGGTCTTCAACGAGGAATTCCTAGTAAGCGCGAGTCATCAGCTCGCGTTGACTACGTCCCTGCCCTTTGTACACACCGCCCGTCGCTCCTACCGATTGAATGGTCCGGTGAAGTGTTCGGATCGCGGCGACGTG
GGCGGTTCGCTGCCGGCGACGTCGCGAGAAGTCCACTGAACCTTATCATTTAGAGGAAGGAGAAGTCGTAACAAGGTTTCCGTAGGTGAACCTGCGGAAGGATCATTG
```

Next is 227 bp spacer between 18S and 28S:

```
TCGAAACCTGCACAGCAGAACGACCCGCGAATTGGTTACAACCGACGGGGGGCGGGGGGCGTTCGTCGCCCCCTCGCCCCCTCCTGCGGGCGGGGACCTCGTGTCTCCTGCCCGCAAACCGAACCCCGGCGCGGAACGCGCCAAGGAAATCTAACCAAGAGAGCCATGCC
GGAGGCCCCGGACACGGTGCGCCCCCGGCGTCGGCGTCTTATGAATTATTCAAAACG
```

Next is 4,129 bp 28S (per RNAmmer); tail ≈300 bp is repetitive and similar to 2,943 bp spacer head:

```
ACTCTCGGCAACGGATATCTAGGCTCTCGCATCGATGAAGAACGTAGCGAAATGCGATACTTGGTGTGAATTGCAGAATCCCGCGAATCATCGAGTTTTTGAACGCAAGTTGCGCCCGAAGCCATTCGGCCGAGGGCACGTCTGCCTGGGTGTCACGCATCGTTGCCCCC
CTCAAACTCCGGTTCGGGCGGGGCGGAAGTTGGCCTCCCGTGCGTGCCTGCACGCGCGGTTAGCCCAAAAGCGAGTCCTCGGCGACGAGCGCCACGACAATCGGTGGTTTTTTTACCCTCGTTCCTCGTCGTGCGTGCCCCGTCGCCCGAACGCGCTCCTCCGACCCTCA
CGCGTCGCCTCGGTGGCGCTCCCAACGCGACCCCAGGTCAGGCGGGACTACCCGCTGAGTTTAAGCATATCAATAAGCGGAGGAAAAGAAACTTACAAGGATTCCCCTAGTAACGGCGAGCGAACCGGGAACAGCCCAGCTTGAGAATCGGGCGCCCTCACGGGCGTCTC
CGAATTGTAGTCTGGAGAAGCGTCCTCAGCGGCGGACCGGGCCCAAGTCCCCTGGAAGGGGGCGCCGGAGAGGGTGAGAGCCCCGTCGTGCCCGGACCCTGTCGCACCACGAGGCGCTGTCGGCGAGTCGGGTTGTTTGGGAATGCAGCCCCAATCGGGCGGTAAATTCC
GTCCAAGGCTAAATACGGGCGAGAGACCGATAGCAAACAAGTACCGCGAGGGAAAGATGAAAAGGACTTTGAAAAGAGAGTCAAAGAGTGCTTGAAATTGTCGGGAGGGAAGCGGATGGGGGCCGGCGATGCGCCCCGGTCGGATGTGGAACGGCGACAGCCGGTCCGCC
GATCGACTCGGGGCGTGGACCGATGCGGATTGCGGCGGCGGCCCAAGCCCGGGCTGTAGTTATGCCCGTGGAGACGTCGTTGCCGCGATCGTGGTTGGCAGCGCGCGCCTCACGGCGTGCCTCGGCATCTGCGCGCTCCTGGCATCGGCCTGCGGGCTCCCCATTCGGCC
CGTCTTGAAACACGGACCAAGGAGTCTGACATGTGTGCGAGTCAACGGGCCAGTAAACCCGTAAGGCGCAAGGAAGCTGATTGGCGGGATCCCCTTGAGGGTTGCACCGCCGACCGACCTTGATCTTCTGAGAAGGGTTCGAGTGAGAGCATACCTGTCGGGACCCGAAA
GATGGTGAACTATGCCTGAGCGGGGCGAAGCCAGAGGAAACTCTGGTGGAGGCCCGCAGCGATACTGACGTGCAAATCGTTCGTCTGACTTGGGTATAGGGGCGAAAGACTAATCGAACCGTCTAGTAGCTGGTTCCCTCCGAAGTTTCCCTCAGGATAGCTGGAGCCCA
CGTGCGAGTTCTATCGGGTAAAGCCAATGATTAGAGGCATCGGGGGCGCAACGCCCTCGACCTATTCTCAAACTTTAAATAGGTAGGACGGCGCGGCTGCTTCGTTGAGCCGCGCCAAGGAATCGAGAGCTCCAAGTGGGCCATTTTTGGTAAGCAGAACTGGCGATGCG
GGATGAACCGGAAGCCGGGTTACGGTGCCCAACTGCGCGCTAACCTAGAACCCACAAAGGGTGTTGGTCGATTAAGACAGCAGGACGGTGGTCATGGAAGTCGAAATCCGCTAAGGAGTGTGTAACAACTCACCTGCCGAATCAACTAGCCCCGAAAATGGATGGCGCTG
AAGCGCGCGACCTATACCCGGCCGTCGGGGCAAGTTCTAGGCCCCGATGAGTAGGAGGGCGCGGCGGTCGCTGCAAAACCTGGGGCGCGAGCCCGGGCGGAGCGGCCGTCGGTGCAGATCTTGGTGGTAGTAGCAAATATTCAAATGAGAACTTTGAAGGCCGAAGAGGG
GAAAGGTTCCATGTGAACGGCACTTGCACATGGGTTAGTCGATCCTAAGAGACGGGGGAAGCCCGTCTGATAGCGTGCTAAGCGCGAGCTTCGAAAGGGAATCGGGTTAAAATTCCTGAACCGGGACGTGGCGGCTGACGGCAACGTTAGGGAGTCCGGAGACGTCGGCG
GGGGCCTCGGGAAGAGTTATCTTTTCTGTTTAACAGCCTGCCCACCCTGGAAACGGCTCAGCCGGAGGTAGGGTCCAGCGGCTGGAAGAGCACCGCACGTCGCGTGGTGTCCGGTGCGCCCCCGGCGGCCCTTGAAAATCCGGAGGACCGAGTGCCTCCCACGCCCGGTC
GTACTCATAACCGCATCAGGTCTCCAAGGTGAACAGCCTCTGGTCGATGGAACAATGTAGGCAAGGGAAGTCGGCAAAATGGATCCGTAACCTCGGGAAAAGGATTGGCTCTGAGGGCTGGGCACGGGGGTCCCAGTCCCGAACCCGTCGGCTGTCGGCGGACTGCTCGA
GCTGCTCCCGCGGCGAGAGCGGGTCGCCGCGTGCCGGCCGGGGGACGGACTGGGAACGACCGCTTCGGCGGTCTTCCCCGGGCGTCGAACAGTCGACTCAGAACTGGTACGGACAAGGGGAATCCGACTGTTTAATTAAAACAAAGCATTGCGATGGTCCCTGCGGATGC
TCACGCAATGTGATTTCTGCCCAGTGCTCTGAATGTCAAAGTGAAGAAATTCAACCAAGCGCGGGTAAACGGCGGGAGTAACTATGACTCTCTTAAGGTAGCCAAATGCCTCGTCATCTAATTAGTGACGCGCATGAATGGATTAACGAGATTCCCACTGTCCCTGTCTA
CTATCCAGCGAAACCACAGCCAAGGGAACGGGCTTGGCAGAATCAGCGGGGAAAGAAGACCCTGTTGAGCTTGACTCTAGTCCGACTTTGTGAAATGACTTGAGAGGTGTAGGATAAGTGGGAGCCGAAAGGCGAAAGTGAAATACCACTACTTTTAACGTTATTTTACT
TATTCCGTGAATCGGAAGCGGGGCTCTGCCCCTCTTTTTGGACCCAAGGCTCGCCTCGGCGGGCCGATCCGGGCGGAAGACATTGTCAGGTGGGGAGTTTGGCTGGGGCGGCACATCTGTTAAAAGATAACGCAGGTGTCCTAAGATGAGCTCAACGAGAACAGAAATCT
CGTGTGGAACAAAAGGGTAAAAGCTCGTTTGATTCTGATTTCCAGTACGAATACGAACCGTGAAAGCGTGGCCTATCGATCCTTTAGACCTTCGGAATTTGAAGCTAGAGGTGTCAGAAAAGTTACCACAGGGATAACTGGCTTGTGGCAGCCAAGCGTTCATAGCGACG
TTGCTTTTTGATCCTTCGATGTCGGCTCTTCCTATCATTGTGAAGCAGAATTCACCAAGTGTTGGATTGTTCACCCACCAATAGGGAACGTGAGCTGGGTTTAGACCGTCGTGAGACAGGTTAGTTTTACCCTACTGATGACAGTGTCGCAATAGTAATTCAACCTAGTA
CGAGAGGAACCGTTGATTCGCACAATTGGTCATCGCGCTTGGTTGAAAAGCCAGTGGCGCGAAGCTACCGTGCGCTGGATTATGACTGAACGCCTCTAAGTCAGAATCCGGGCTAGAAGCGACGCGTGCGCCCGCCGCCCGATTGCCGACCTGCAGTAGGGGCCCTCGGG
CCCCCAGAGGCACGTGTCTTTGGCCAAGCCCCCGCGGCGGACGAGCCGCGTGGGCCGCCATGAAGTATAATTCCCACCGAGCGGCGGGTAGAATCCTTTGCAGACGACTTAAATACGCGACGGGGTATTGTAAGTGGCAGAGTGGCCTTGCTGCCACGATCCACTGAGAT
TCAGCCCTGTGTCGCTTCGATTCGTCCCTCCCCCCTCTCCTCTCCCAATCCCCCCCTCAAAAAAAAAATCAACTCTTTGCCCCGTGCCCTCCGGAGGCTAGCCCCCCGAGTGCCTTCGCTGCGTGCCCCTTGCCCATGACAGGCTGCCTTGGCCTTGGCCTTCGCTGCTT
GCCTATGGGGGGTGCCTTGGCCTTGGCTGCGTGCCCTTGCCTTGGCAGCATGCCCATGGGGGGCTGCTGCGTGCCCTTGCCCTTGGCTGCATGCCCGTGGGGGGCTGCTGCTGCCTTGGCCTTGGCTGCATGCCTTGGCCTTGGCCTTGGCTGCATGCCTTGGCCTTGGC
CTTGGCTGCGTGCCTTGGCCTTGGCCTTGGCTGCATGCCTTGGCCTTGG
```

Final 18 bp is not included by RNAmmer as part of 28S, but is part of repetitive tail approximately closing circle:

```
CTGCATGCCATGGCCTTG
```

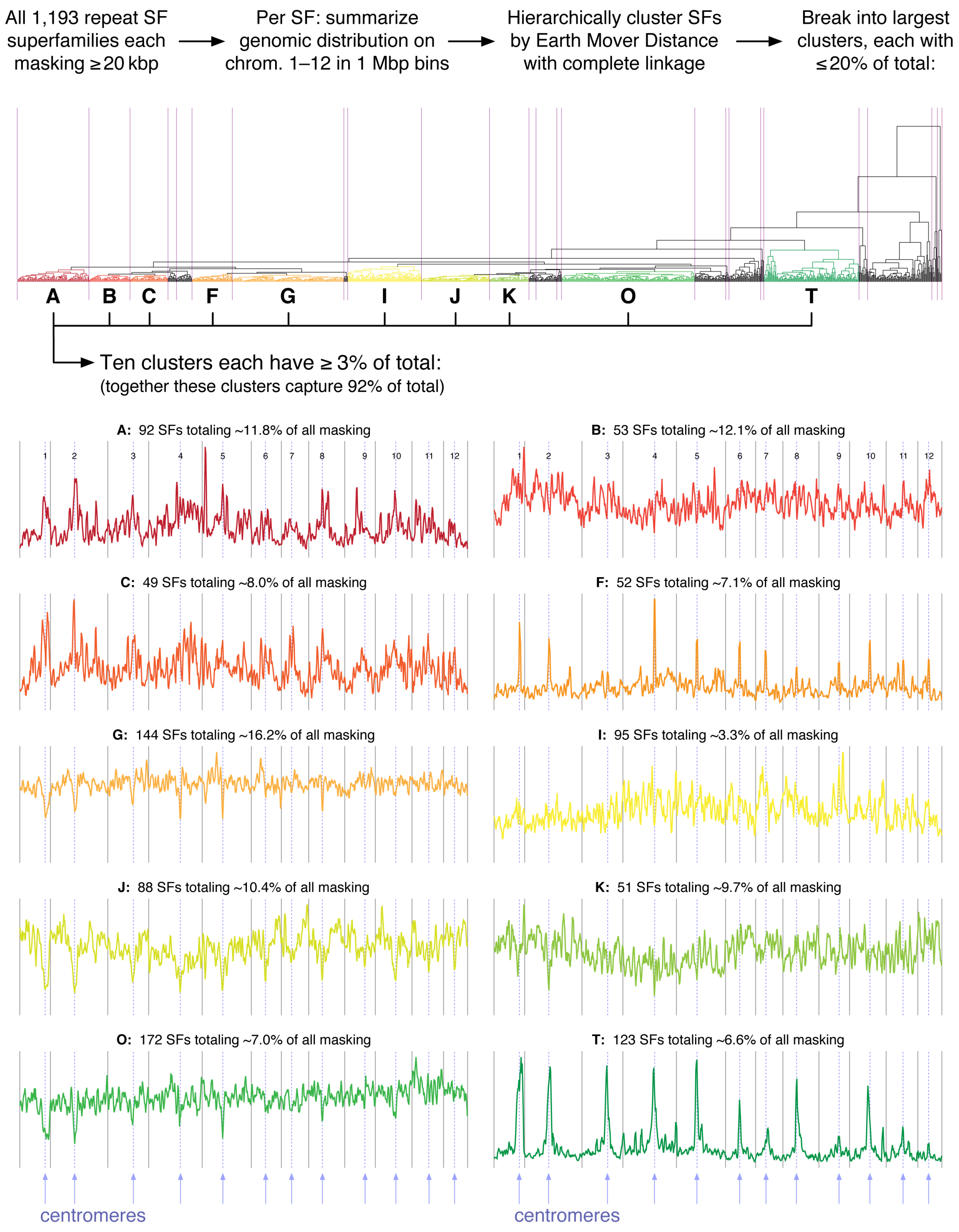

**Figure S11.** Unsupervised clustering indicates that the dominant chromosome-scale distributional features of repeats in *Q. lobata* are correlated with distance to the centromeres.

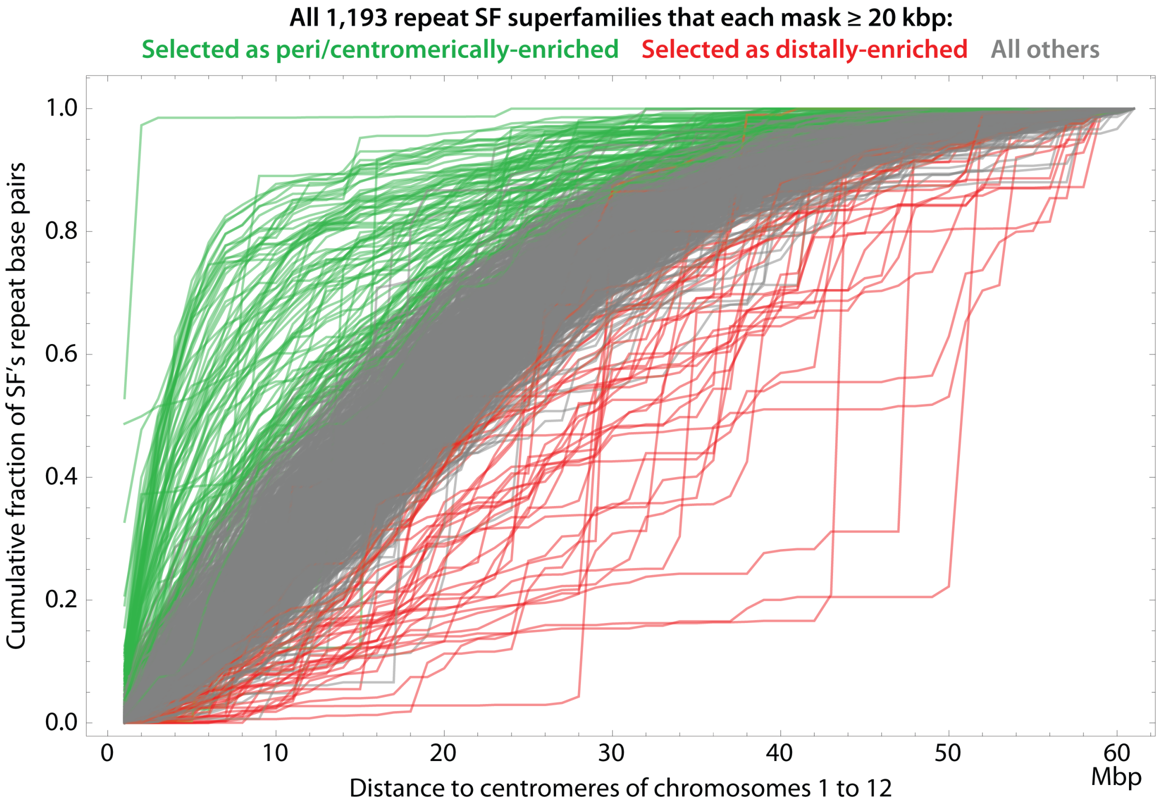

**Figure S12. Average centromeric distance summarizes repeat per-SF distribution of distances well.** For each repeat superfamily (SF) of minimal size (≥ 20 kbp masked), the cumulative distribution function (CDF) of the distance of the repeat's base pairs to the centromeres is shown. Coloring by green, red, and gray is as in Figure 3C and shows that the average centromeric distance per SF summarizes the distributions well, with outliers at the distribution level essentially coinciding with outliers at the average level. Curve plotting order is randomized across all 1,193 SFs.

#### VI. Gene prediction and annotation

Basic statistics for the three *Quercus* protein-coding gene (PCG) sets are given in Table S3. (For *Q. suber*, 12% of PCG loci have multiple transcript models; a single longest isoform was chosen per locus.) *Q. robur* has many fewer models (26k), while *Q. suber* more (49k, but a more comparable 36k with at least one intron). Further, at least 11k *Q. suber* models are incomplete, and 1k actually interpolate CDS beyond the assembly. By models-per-Mbp-of-non-gap-assembly, *Q. robur* is low (33), with *Q. lobata* and *Q. suber* quite similar (47 and 53). *Q. robur* calls a total of only 30 Mbp of CDS (gene spans cover just 10% of its assembly) vs. 50 and 67 Mbp (25% and 20%) for *Q. lobata* and *Q. suber*. While every *Q. lobata* model has both UTR5 and UTR3 annotated (affecting many size-related quantities of Table S3), only about half of *Q. robur* and *Q. suber* models have UTRs, with *Q. suber* tending short and *Q. robur* shorter when they do have UTRs. CDS lengths are fairly similar (and have similar depletion of repetitive sequence), although *Q. suber* and (to a lesser extent) *Q. robur* tend to have fewer exons, perhaps due to their higher assembly incontiguity. While no *Q. lobata* models have CDS that contain assembly gaps, 0.2% and 0.8% do so in *Q. suber* and *Q. robur*; for exons, this rises to 0.2% vs. 0.3% and 1.1%, and for gene spans to 0.6% vs. 6.5% and 6.9% (suggesting the other assemblies unsurprisingly have gaps concentrated in introns). Based on a HMMer search for GyDB domains, *Q. robur* is the most conservative, where only 0.1% of models have at least one domain strongly indicative of a transposon (rising to 0.5% for domains correlated with transposons); *Q. lobata* is somewhat higher (0.7% and 1.4%), but *Q. suber* is much higher (3.0% and 4.6%).

**Table S3. Statistics of protein-coding gene (PCG) models for *Q. lobata*, *Q. robur*, and *Q. suber*.**

| **Statistic** | ***Q. lobata*** | ***Q. robur*** | ***Q. suber*** |
|---|---|---|---|
| **# PCG (Protein-Coding Gene models)[*]** | **39,373** | **25,808** | **49,388** |
| # PCG$^5$ (PCGs with non-empty UTR5) | 39,373 | 13,625 | 24,282 |
| # PCG$^3$ (PCGs with non-empty UTR3) | 39,373 | 14,132 | 24,348 |
| # PCG$^i$ (PCGs with at least one intron) | 34,859 | 20,356 | 35,822 |
| # PCGs with an ostensibly complete[†] CDS | 39,373 | 25,808 | 38,499 |
| # PCGs with CDS not entirely[‡] from the assembly | 0 | 0 | 1,151 |
| # PCGs with ≥ 1 H [H or M][§] transposon domain | 288 [537] | 21 [130] | 1,462 [2,275] |
| Knt between adj. PCG spans: average [median] | 15.6 [8.5] | 27.7 [14.8] | 13.1 [5.3] |
| Span kilobases per PCG: average [median] | 5.4 [4.2] | 3.1 [2.3] | 3.9 [2.3] |
| Exonic kilobases per PCG: average [median] | 2.3 [2.0] | 1.3 [1.1] | 1.6 [1.4] |
| CDS kilobases per PCG: average [median] | 1.3 [1.0] | 1.2 [0.9] | 1.4 [1.1] |
| UTR5 kilobases per PCG$^5$: average [median] | 0.4 [0.3] | 0.2 [0.1] | 0.2 [0.2] |
| UTR3 kilobases per PCG$^3$: average [median] | 0.7 [0.5] | 0.1 [0.1] | 0.3 [0.2] |
| Intronic kilobases per PCG$^i$: average [median] | 3.5 [2.4] | 2.2 [1.5] | 3.1 [1.5] |
| # exon intervals per PCG: average [median] | 5.5 [4.0] | 4.4 [3.0] | 4.1 [3.0] |
| # CDS intervals per PCG: average [median] | 4.8 [3.0] | 4.3 [3.0] | 3.9 [2.0] |
| # UTR5 intervals per PCG$^5$: average [median] | 1.3 [1.0] | 1.0 [1.0] | 1.2 [1.0] |
| # UTR3 intervals per PCG$^3$: average [median] | 1.4 [1.0] | 1.0 [1.0] | 1.1 [1.0] |
| Mbp in union of all PCG… exons [CDS] | 92.2 [49.9] | 34.8 [30.3] | 78.6 [67.0] |
| Mbp in union of all PCG… UTR5 [UTR3] | 15.7 [26.6] | 2.4 [2.1] | 4.8 [6.8] |
| Mbp in union of all PCG… introns [spans] | 121.2 [213.5] | 45.1 [79.9] | 111.1 [189.3] |
| % of asm. in union PCG… exons [CDS] | 10.9% [5.9%] | 4.3% [3.7%] | 8.2% [7.0%] |
| % of asm. in union PCG… UTR5 [UTR3] | 1.9% [3.1%] | 0.3% [0.3%] | 0.5% [0.7%] |
| % of asm. in union PCG… introns [spans] | 14.3% [25.2%] | 5.5% [9.8%] | 11.7% [19.9%] |
| % of non-gap assembly that is repetitive[\|] | 54.4% | 54.3% | 51.6% |
| % repetitive[\|] in union PCG… exons [CDS] | 16% [14%] | 12% [13%] | 13% [14%] |
| % repetitive[\|] in union PCG… UTR5 [UTR3] | 18% [18%] | 17% [6%] | 12% [8%] |
| % repetitive[\|] in union PCG… introns [spans] | 26% [22%] | 18% [16%] | 30% [23%] |
| % PCG w/ ≥ 1 asm. gap in… exons [CDS] | 0.2% [0.0%] | 1.1% [0.8%] | 0.3% [0.2%] |
| % PCG w/ ≥ 1 asm. gap in… UTR5 [UTR3] | 0.1% [0.1%] | 0.2% [0.1%] | 0.0% [0.0%] |
| % PCG w/ ≥ 1 asm. gap in… introns [span] | 0.4% [0.6%] | 6.0% [6.9%] | 6.3% [6.5%] |

[*] Explicit non-nuclear assembly components are removed, and only a single longest PCG isoform is kept per PCG-containing gene locus.

[†] *Ostensibly complete* means the CDS, as derived exclusively from the assembly, starts on a codon boundary with a start codon, ends on a codon boundary with a stop codon, and has no internal stop codons.

[‡] The NCBI genebuild pipeline (*Q. suber*) can make models that apply edits (e.g., additions of 100's of basepairs) to the reference assembly; generally all table data is based on the pure-assembly portions.

[§] GyDB 2019-03-21 HMMer 3.2.1 full-sequence hits of E-value $\leq 10^{-5}$; 'H' (high) is ≥ 1 of GAG/GAGCOAT/RT/INT/galadriel/TAV, 'M' (medium) is ≥ 1 of AP/RNaseH/CHR/DUT/MOV/ENV.

[|] All (non-gap) basepairs that are masked by a run of RepeatMasker-after-RepeatModeler (as in Figure 3A) for each assembly.

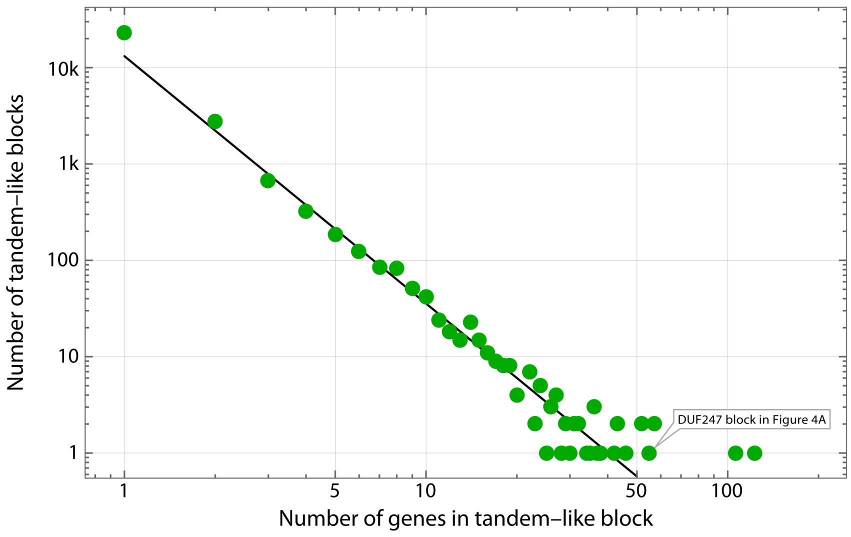

**Figure S13**. Log–log frequency versus size of tandem-like duplicated gene blocks. Black line is fitted power decay rate (number ≈ 13,139 / size$^{2.567}$).

**R-gene identification.** Gururani, et al. [44] provide an overview of the numerous types of plant disease resistance genes (R-genes), which allow plants to detect a pathogen attack from bacteria, viruses, nematodes, oomycetes, fungi, and insects and facilitate a counter attack against the pathogen. In reviewing studies of R-genes, they propose eight classes of R-gene architecture comprised of membrane spanning domains, amino acid motifs and other domains as follows:

I. Genes encoding for proteins with a nucleotide-binding site (NBS), a leucine rich repeat (LRR) and a putative coiled coil domain (CC) at the N- terminus.
II. Cytoplasmic proteins with LRR and NBS motifs and (TIR) domain.
III. Extra cytoplasmic leucine rich repeats (eLRR), attached to a transmembrane domain (TrD) , without a NBS motif.
IV. An extracellular LRR domain, a transmembrane domain (TrD) and an intracellular serine-threonine kinase (KIN) domain.
V. The putative extracellular LRRs, along with a PEST (Pro-Glu-Ser-Thr) domain for protein degradation, and short proteins motifs (ECS).
VI. A membrane protein domain (TrD), fused to a putative coiled coil domain (CC)
VII. The Arabidopsis RRS1-R gene conferring resistance to the bacterial phytopathogen *Ralstonia solanacearum*, and it is a new member of the TIReNBSeLRR R protein class. RRS1-R has a C-terminal extension with a putative nuclear localization signal (NLS) and a WRKY domain.
VIII. The enzymatic R-genes which contain neither LRR nor NBS groups.

We represent these groups as:

1. ``NBS LRR TIR''
2. ``NBS LRR CC''
3. ``LRR TrD''
4. ``LRR TrD KINASE''
5. ``TrD CC''
6. ``LRR TrD PEST''
7. ``TIR NBS LRR NLS WRKY''
8. ``KINASE + KINASE KINASE + HM1''

In our analysis, we kept track of whether it has zero versus one or more copies of the following classes of Pfam domains, Coils, and transmembrane hits found in the oak protein coding genes in the assemblies of *Q. lobata*, *Q.*

*robur*, and *Q. suber*. We identified the coiled coils and transmembranes using TMHMM 2.0c for transmembrane prediction and Coils 2.2.1 for coiled coils, via InterProScan 5.34-73.0. Other domains are panther-based names.

N := NB-ARC
T := TIR / TIR_2
L := LRR_1..6,8,9 / LRRNT_2
P := Pkinase / Pkinase_C / Pkinase_Tyr
C := coiled coil
M := transmembrane

To parallel the major classes of Gururani et al. [44], we assigned each of the PGCs ,Ignoring order and copy number, to these eight patterns of R-genes.

1. NTL___
2. N_L_C_
3. __L__M
4. __LP_M
5. ____CM
6. __L__M + PEST
7. NTL___ + NLS + WRKY
8. ___P__ + ___P__ + HM1

We then simplified the reduced the number of patterns with the following rules. Pattern 4 is more restrictive than pattern 3, so we dropped it. Pattern 6 is more restrictive than pattern 3, so we dropped it. Pattern 7 is more restrictive than pattern 1, so we dropped it. Pattern 8 is again a strange case because the `___P__'s generate many false positives and it is unclear what Pfam domains correlate with what they call `HM1'. In the end, we used the four patterns below as a superset of the domains in the remaining patterns, using `*' as a wildcard:

1. NTL***
2. N*L*C*
3. **L**M
5. ****CM

To obtain counts of R-genes in each oak species (Table S4), we assigned R-gene associated domain enriched PGCs as R-genes if they included:

(Pfam `NB-ARC' domain) OR (Pfam `TIR' or `TIR_2' domain), OR Pfam `LRR_1', `LRR_2', `LRR_3', `LRR_4', `LRR_5', `LRR_6', `LRR_8', `LRR_9', or `LRRNT_2' domain), OR
TMHMM transmembrane segment AND [Pfam `Pkinase', `Pkinase_C', o `Pkinase_Tyr' domain])

We also assigned these R-genes to confidence categories based on the descriptions below. Based on the first highest certainty category, we counted 751 R-genes for Q. lobata, 632 R-genes for Q. robur, and 723 R-genes for Q, suber. However, these three oaks may have many more R-genes. Our other two categories for R-genes yield an additional 2176, 1645, and 2182 candidate R-genes for the three species, respectively.

**Categories of genes assigned as R-genes, with separation of the patterns into confidence categories**

STRONG EVIDENCE FOR R-gene assignment: few unnamed domains and named domains look okay
HIGH EVIDENCE For R-genes assignment: named domains look okay but a substantial or high fraction are unnamed.
ENRICHED domains for R-genes: most domains are unnamed but ~3/4ths of named domains are R-suggestive
MARGINALLY POSSIBLE as R-genes: low numbers so doesn't affect total count of R-genes

**Categories of genes not assigned as an R-gene**

LOW (but non-zero) evidence for R gene assignment: only ~10% of named domains are suggestive of R-genes
MODERATE NEGATIVE EVIDENCE FOR R-GENE ASSIGNMENT: (no domain names or generic kinases)
STRONGER NEGATIVE EVIDENCE FOR R-GENE ASSIGNMENT: few R-gene domains but will include some R-genes

**Table S4.** Counts of R-genes found in three oak species that include combinations of domains identified by Gururani et al.[44] as one of 8 classes of plant resistance genes. The domain combinations are grouped by likelihood of acting as an R-gene and degree of enrichment. (See text above fore details)

| | Domain combinations | Q. lobata (39,373) | Q. robur (25,808) | Q. suber (49,388) |
|---|---|---|---|---|
| Strongly Enriched R-genes | NT____ | 180 | 128 | 157 |
| | N_L_C_ | 76 | 43 | 42 |
| | NTL___ | 68 | 55 | 60 |
| | N_L___ | 35 | 56 | 48 |
| | NT__C_ | 19 | 9 | 32 |
| | N___CM | 13 | 13 | 7 |
| | NT___M | 10 | 4 | 5 |
| | NTL__M | 6 | 3 | 2 |
| | NTL_C_ | 6 | 11 | 5 |
| | N_L_CM | 5 | 1 | 3 |
| | NT__CM | 4 | 0 | 1 |
| | _T__CM | 1 | 0 | 1 |
| | _TL__M | 1 | 0 | 0 |
| | Subtotals | 424 | 323 | 363 |
| Highly Enriched R-genes | __L__M | 302 | 295 | 330 |
| | __LPCM | 14 | 9 | 22 |
| | N____M | 11 | 5 | 8 |
| | Subtotals | 327 | 309 | 360 |
| Enriched potential R-genes | ___P_M | 754 | 466 | 663 |
| | __LP_M | 382 | 234 | 342 |
| | N___C_ | 344 | 223 | 308 |
| | N_____ | 266 | 314 | 347 |
| | __L___ | 241 | 228 | 356 |
| | _T____ | 102 | 139 | 101 |
| | ___PCM | 65 | 28 | 39 |
| | __L_C_ | 22 | 13 | 26 |
| | Subtotals | 2176 | 1645 | 2182 |
| Possibly some R-genes | _T___M | 15 | 10 | 6 |
| | __L_CM | 4 | 7 | 13 |
| | _T__C_ | 3 | 8 | 4 |
| | __LPC_ | 2 | 1 | 4 |
| | N__P__ | 2 | 1 | 0 |
| | _TL___ | 0 | 0 | 1 |
| | N__P_M | 0 | 1 | 0 |
| | N_L__M | 0 | 4 | 1 |
| | Subtotals | 26 | 32 | 29 |
| Unlikely many R-genes | __LP__ | 34 | 18 | 23 |
| | Subtotals | 34 | 18 | 23 |
| Few, if any, R-genes | ___P__ | 729 | 657 | 873 |
| | ____CM | 725 | 405 | 925 |
| | ___PC_ | 83 | 43 | 113 |
| | Subtotals | 1537 | 1105 | 1911 |
| Not R-genes | _____M | 7007 | 5017 | 8251 |
| | ____C_ | 4143 | 2342 | 6303 |
| | Subtotals | 11150 | 7359 | 14554 |

#### VII. Methylomes and analysis of methylation patterns

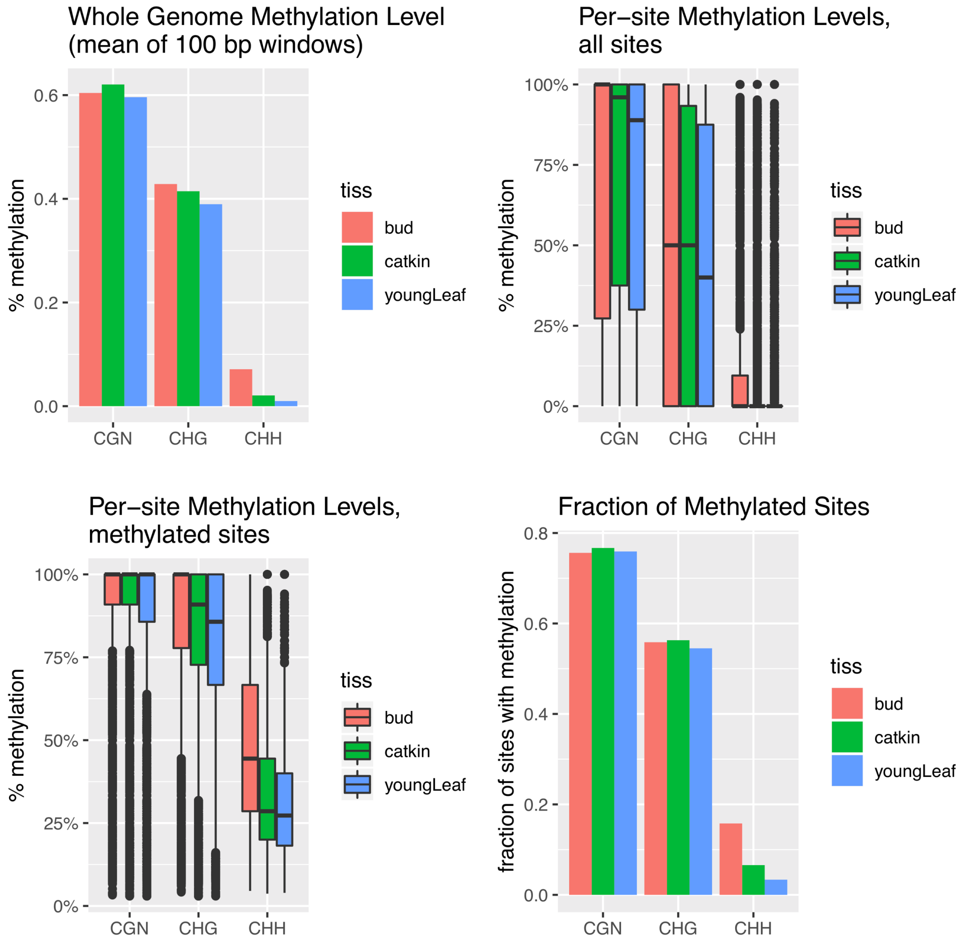

**Figure S14.** Genome methylation levels for three tissues and three methylation contexts. Whole genome average methylation was calculated by averaging the methylation levels for 100 bp windows across chr. 1–12. Per-site methylation levels are for sites with a minimum strand-specific coverage of three reads, and is shown for all such sites and for sites considered methylated (by MethylDackel's binomial test for above background/non-conversion). Also shown are the fraction of sites that are considered methylated (minimum coverage of three reads). Methylation levels are consistent with a total absence (i.e., at bisulfite non-conversion estimated as ≈0.5%) at the majority of CHH sites (84%–97%), a large portion of CHG sites (43%–45%), and some CG sites (24%–25%), with methylation averages for the remaining sites much higher than the overall averages (mCHH 27%–45%, mCHG 86%–100%, mCG 100%).

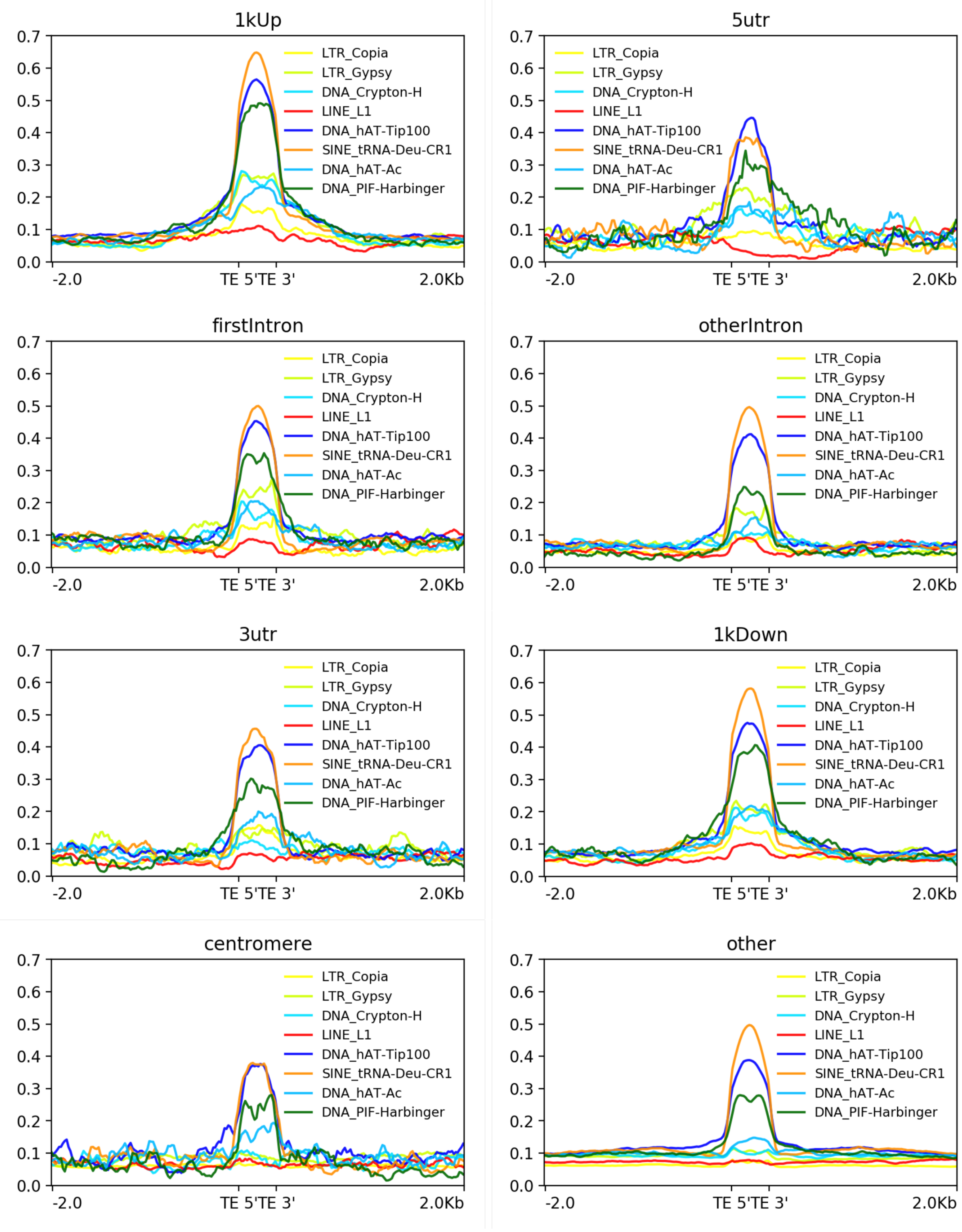

**Figure S15.** CHH methylation levels in bud tissue across repeats. The “SINE_tRNA-Deu-CR1”, “DNA_hAT-Tip100”, and “DNA_PIF-Harbinger” show consistently high levels of mCHH across all regions. CDS regions are not shown due to small numbers of instances for some superfamilies. The “DNA_CMC-ENspm” and “DNA_MuLE-MuDR” were removed due to too few instance bases in several regions.

**Figure S16.** (3 pages)**.** Subcontext methylation for *Q. lobata* chromosomes 1 to 12 in 1 Mbp windows. For each chromosome, **top panel** is number of protein coding genes, **middle panel** is mean mC̲HG by 3 nt subcontext, and **bottom panel** is mean mC̲HH by 3 nt subcontext.

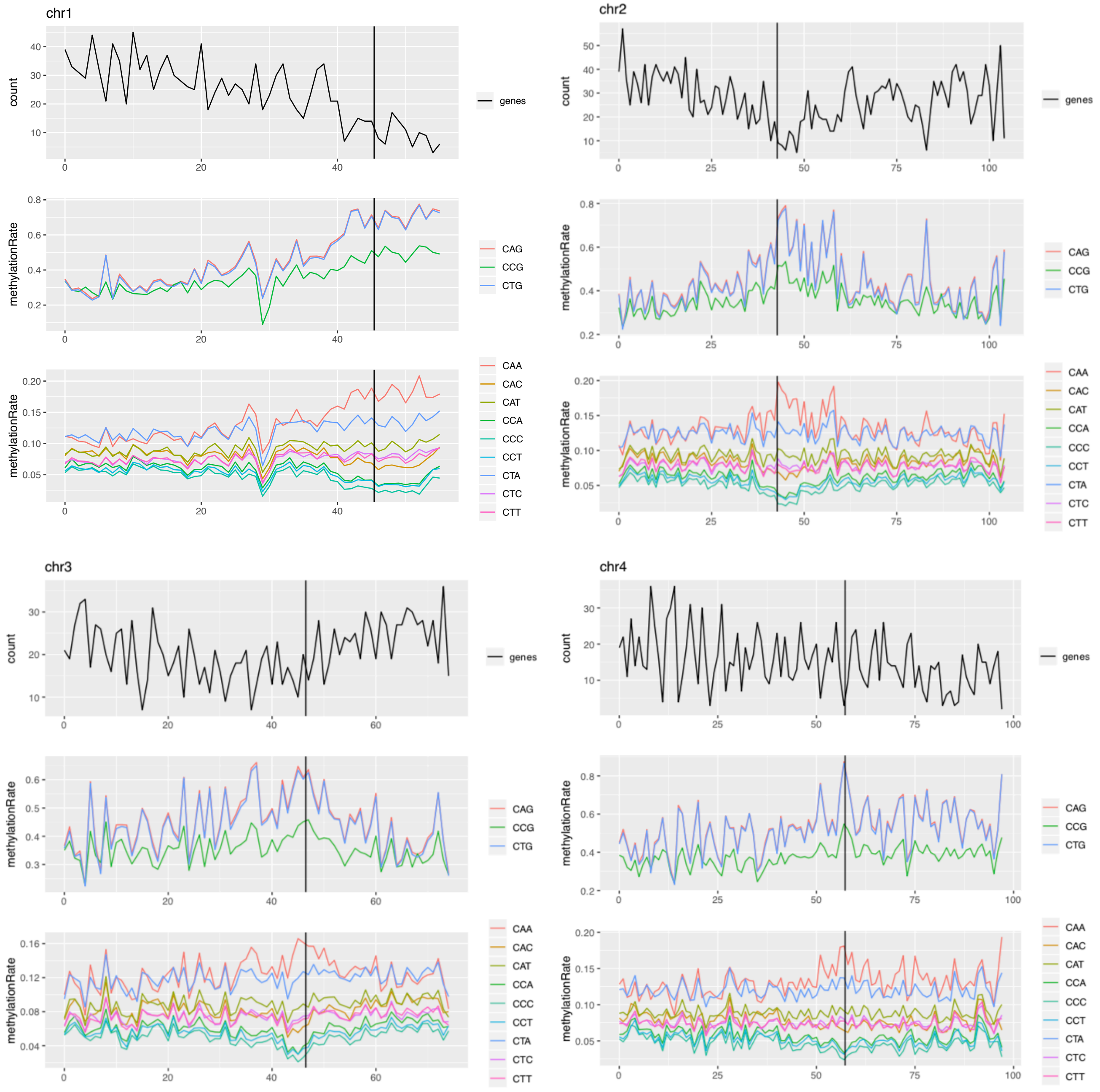

**Figure S16. (part 2 of 3 parts)**

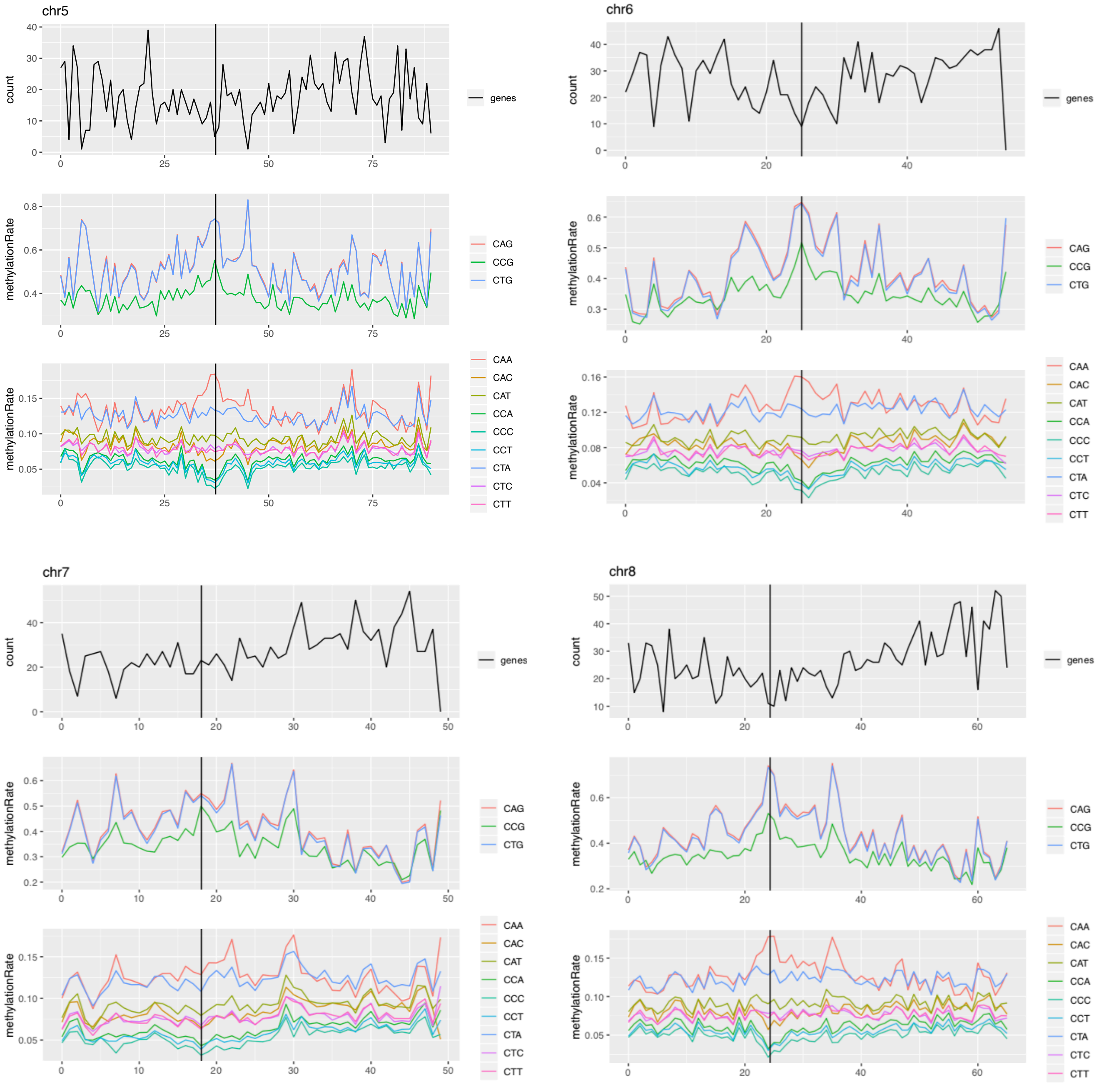

**Figure S16. (part 3 of 3 parts)**

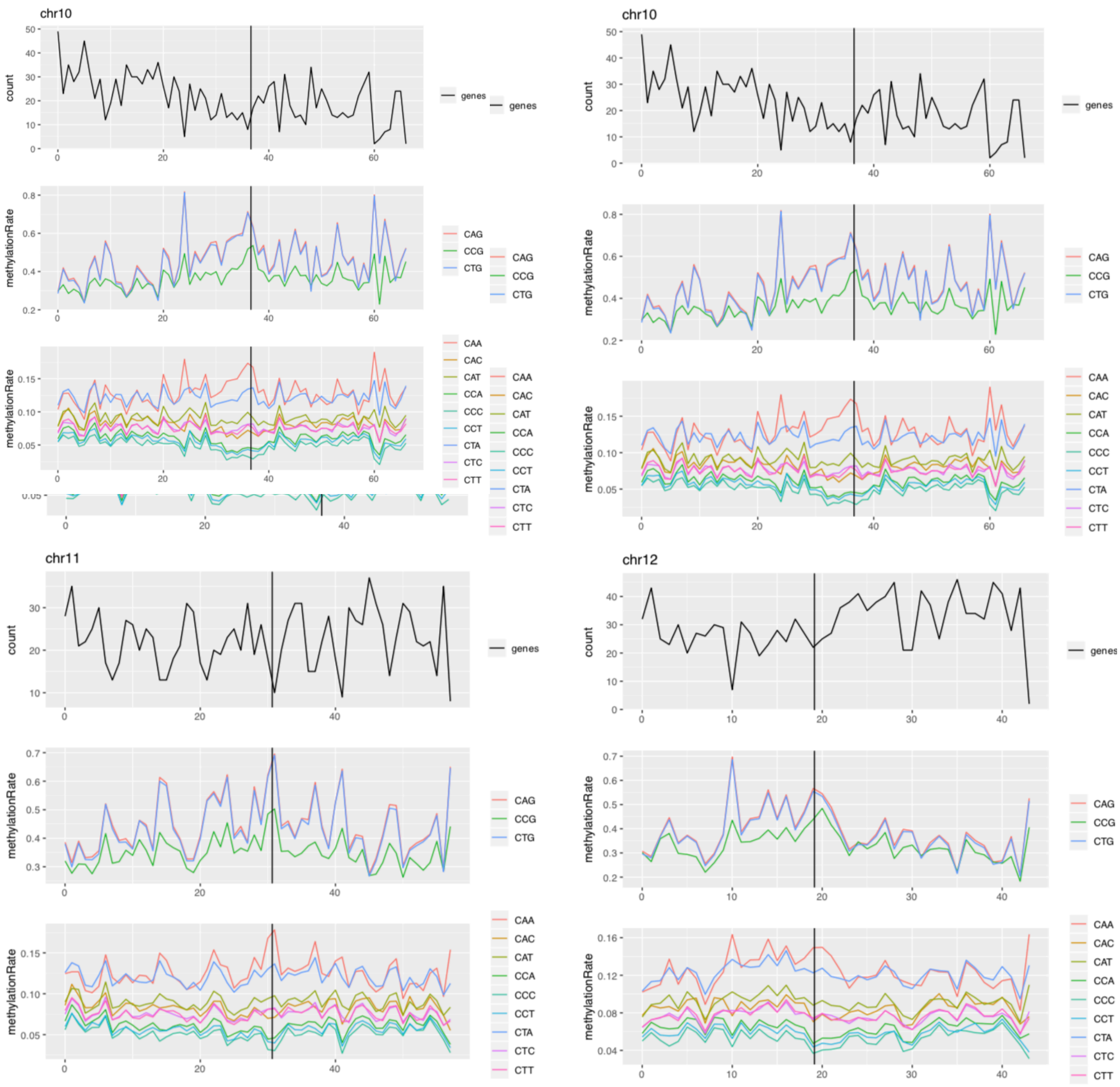

**Figure S17.** Intergenic subcontext mCHH by size of region. Average bud tissue mCHH by 3 nt subcontext for intergenic regions, from a protein-coding gene's transcription end site (TES) to the next PCG's transcription start site (TSS), normalized to 5 kbp long and separated into six intergenic size ranges (one range per panel).

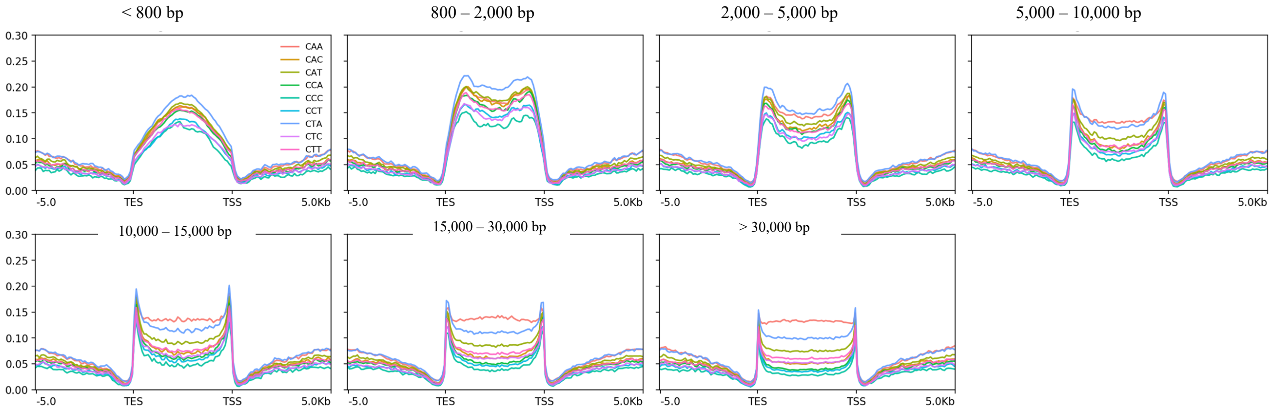

**Figure S18.** Genic region methylation of oak in comparison with 34 angiosperms. Plots show mCG (blue), mCHG (green), and mCHH (maroon) upstream, across, and downstream averaged over genes, and are reprinted from Figure S18 from Niederhuth, et al. [1], with oak (*Q. lobata* young leaf and bud) added for comparison.

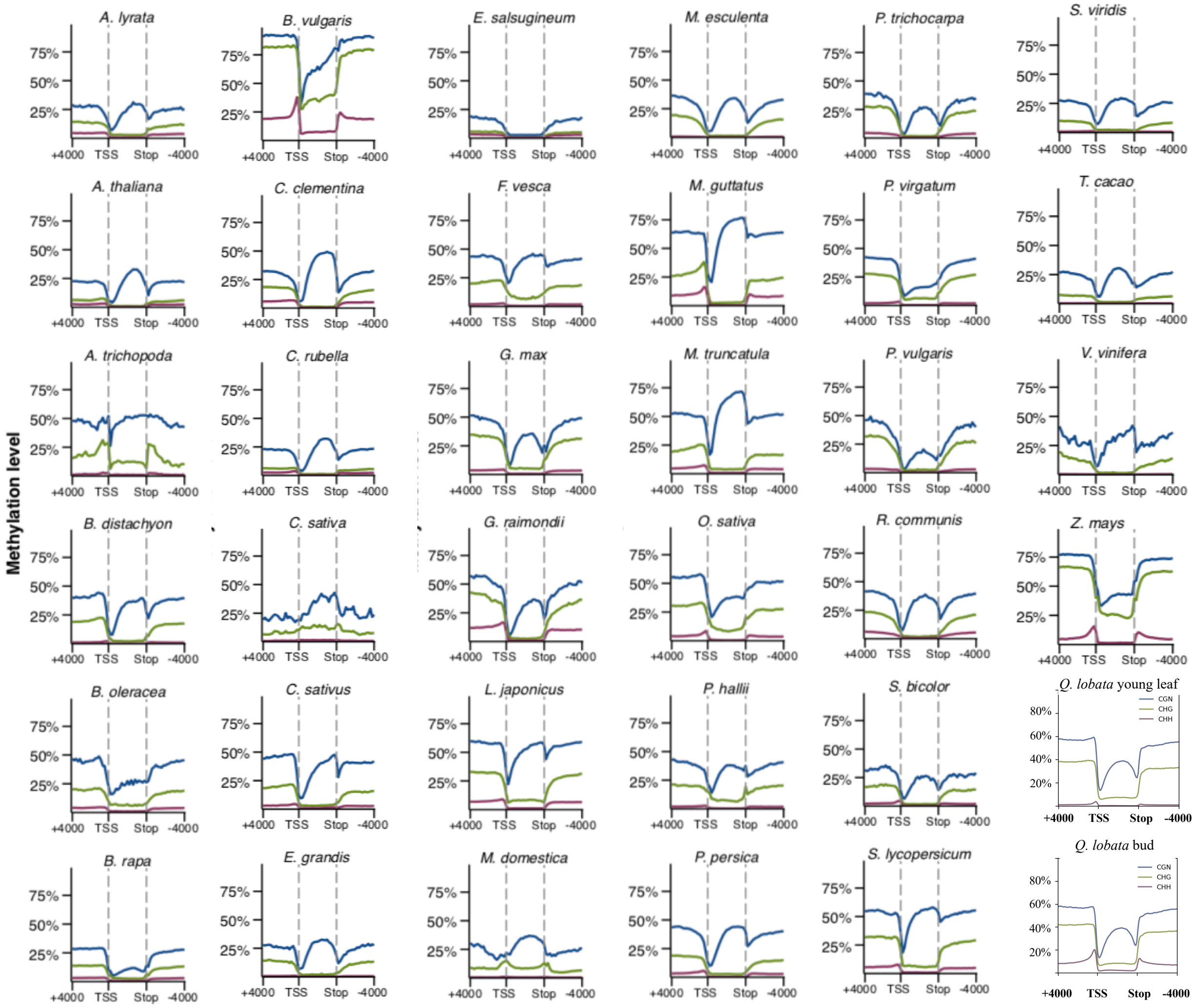

**Figure S19** (2 pages)**.** Chromosomal overviews of methylation and protein-coding gene (PCG) counts. Plots are reprinted from Figure S10 in Niederhuth, et al. [1], limited to 24 taxa each having a chromosome-level assembly, to which we add plots for *Quercus lobata* with as similar methods as possible. **(A)** Subpanels are ordered column to column approximately by PCG density, from heterogeneous to homogeneous. This order is roughly by magnitude of difference between smallest (mostly pericentromeric) and largest (mostly chromosome arms) gene count windows. Subpanels show methylation levels and gene counts for 100 kbp windows every 50 kbp across chromosome 1 or the largest scaffold for each taxon. For *Q. lobata*, chromosome 2 was used since chromosome 1 is unusual (the lone acrocentric chromosome). mCG is shown in blue, mCHG in green, mCHH in maroon, and gene counts in red. Despite having a relatively high total PCG count (39,373), oaks are among the lowest for chromosome arm gene density, with only maize clearly lower. Methylation levels also usually correlate with prevalence of repeats, and show very distinct patterns in the initial columns vs. much more homogenous levels toward the later columns. Gene count *y*-axis upper limit is variable (determined by peak). (**B**) Plots for *Z. mays* and *Q. lobata* are placed side by side to show similarity, with the *Q. lobata* gene count *y*-axis scale made to match *Z. mays*. (**C**) Plots of methylation and PGC for all *Q. lobata* twelve chromosomes.

(**A**)

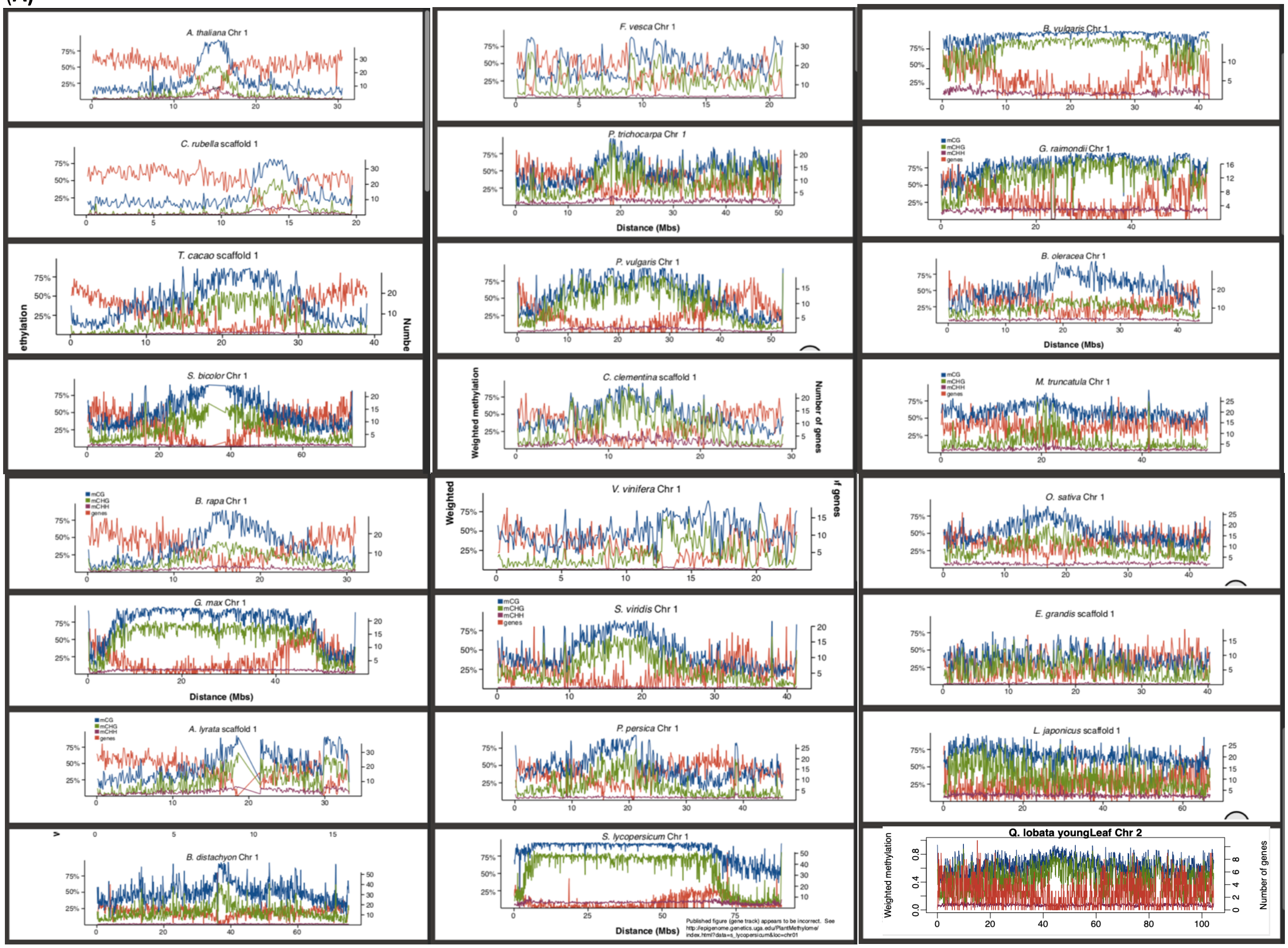

**(B)**

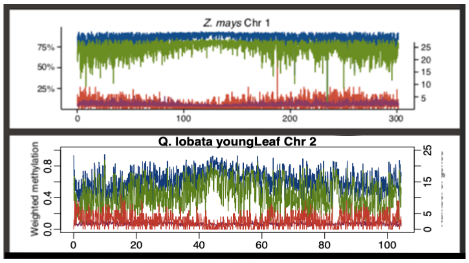
Z. mays Chr 1
Q. lobata youngLeaf Chr 2
Weighted methylation

**(C)**

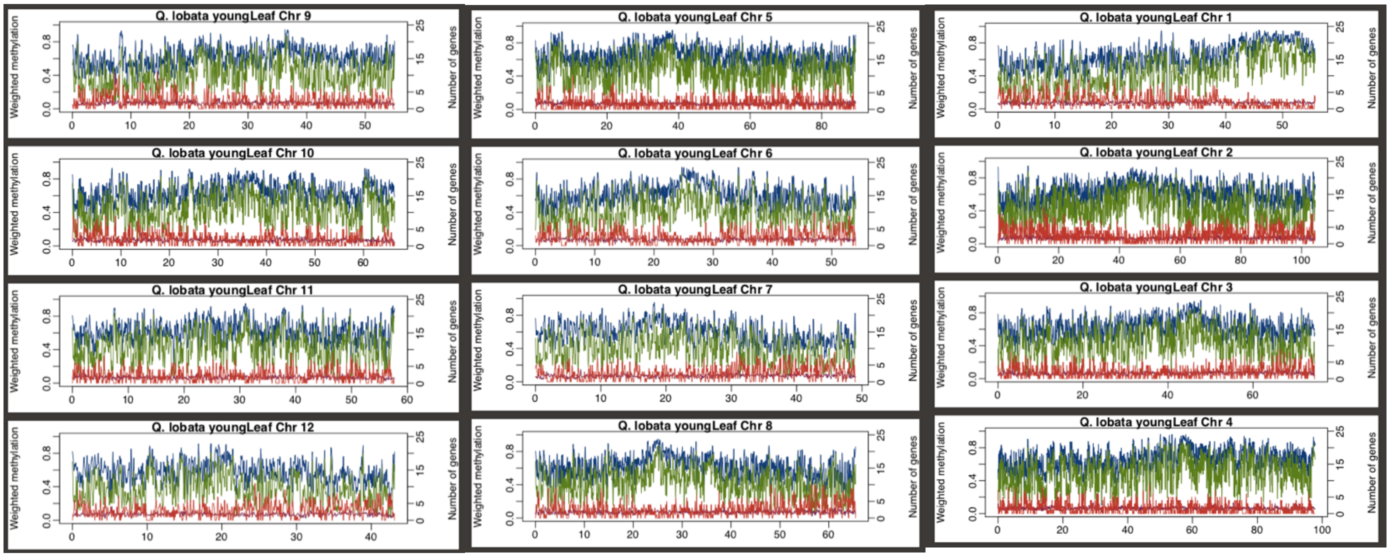
Q. lobata youngLeaf Chr 9
Q. lobata youngLeaf Chr 5
Q. lobata youngLeaf Chr 1
Q. lobata youngLeaf Chr 10
Q. lobata youngLeaf Chr 6
Q. lobata youngLeaf Chr 2
Q. lobata youngLeaf Chr 11
Q. lobata youngLeaf Chr 7
Q. lobata youngLeaf Chr 3
Q. lobata youngLeaf Chr 12
Q. lobata youngLeaf Chr 8
Q. lobata youngLeaf Chr 4
Weighted methylation
Number of genes

**Figure S20.** Subcontext methylation for *Populus trichocarpa* chromosomes 1 to 19. Mean mC̲HH by 3 nt subcontext, 1 Mbp window every 1 Mbp. Methylation data is from tree 13.1 of Hofmeister. *et al.* [45]

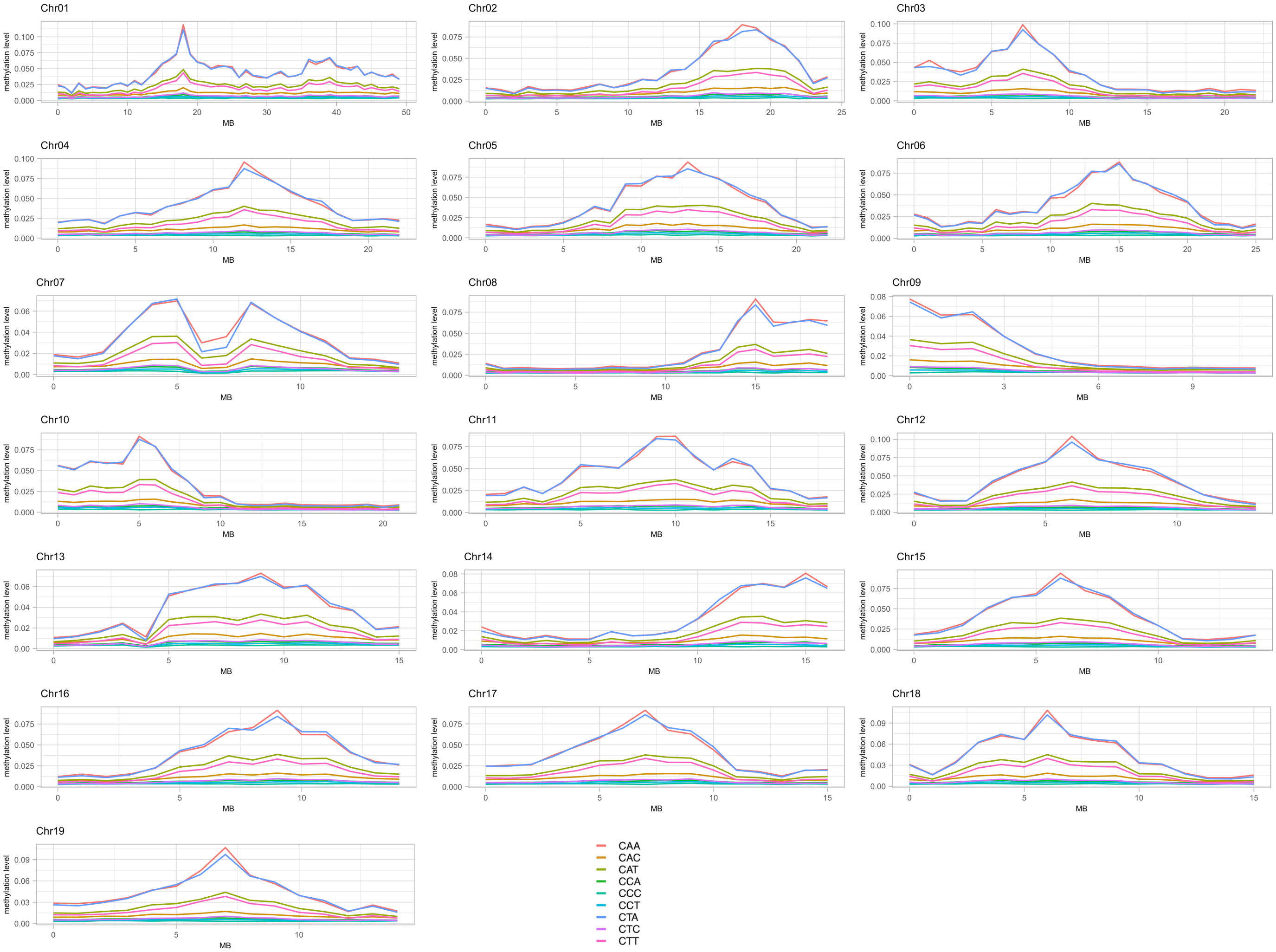

#### XII. Additional Tables

**Table S5**. Top Pfam domains enriched in the most heavily tandemly duplicated PCG families. The 414 PCGs in the enrichment set are those participating in at least one tandem block of size 30 PCGs. Each list shows enrichment for Pfam domains with Benjamini-Hochberg FDR-adjusted *q*-value < 0.1, or hypergeometric *p*-value < 0.0002. (Subsetted from Auxiliary Spreadsheet 1 (AuxSpread1_SJC-PCG-HgeoEnrichments--GeneNameWord-Pfam—20200717.xlsx.) Tandemness is defined via global amino acid identity ≥ 30%.

| *Pfam short name* | *Hgeo. p-value* | *BH FDR q-value* | *Obs./ expect* | *Subset has…* | *in:* | *Bkgnd. has…* | *in:* | *Pfam accn.* | *Pfam type* | *Pfam long name* |
|---|---|---|---|---|---|---|---|---|---|---|
| **DUF247** | 6.69E-87 | 2.78E-83 | 23.06 | 80 | 1,451 | 185 | 77,362 | PF03140 | Family | Plant protein of unknown function |
| **Stress-anti-fung** | 5.41E-67 | 1.12E-63 | 13.54 | 82 | 1,451 | 323 | 77,362 | PF01657 | Family | Salt stress response/antifungal |
| **FBA_3** | 1.14E-54 | 1.58E-51 | 14.15 | 65 | 1,451 | 245 | 77,362 | PF08268 | Domain | F-box associated domain |
| **NB-ARC** | 8.58E-48 | 8.91E-45 | 5.39 | 113 | 1,451 | 1,118 | 77,362 | PF00931 | Domain | NB-ARC domain |
| **FBA_1** | 1.05E-33 | 8.69E-31 | 11.79 | 44 | 1,451 | 199 | 77,362 | PF07734 | Family | F-box associated |
| **ADH_N_2** | 5.01E-29 | 3.47E-26 | 34.99 | 21 | 1,451 | 32 | 77,362 | PF16884 | Family | N-terminal domain of oxidoreductase |
| **F-box** | 1.29E-28 | 7.66E-26 | 5.64 | 64 | 1,451 | 605 | 77,362 | PF00646 | Domain | F-box domain |
| **Pkinase** | 6.49E-28 | 3.37E-25 | 3.09 | 123 | 1,451 | 2,125 | 77,362 | PF00069 | Domain | Protein kinase domain |
| **Pkinase_Tyr** | 1.19E-26 | 5.50E-24 | 2.99 | 123 | 1,451 | 2,196 | 77,362 | PF07714 | Domain | Protein tyrosine kinase |
| **ADH_zinc_N** | 1.16E-21 | 4.84E-19 | 12.20 | 27 | 1,451 | 118 | 77,362 | PF00107 | Family | Zinc-binding dehydrogenase |
| **ADH_zinc_N_2** | 1.74E-18 | 6.56E-16 | 18.46 | 18 | 1,451 | 52 | 77,362 | PF13602 | Domain | Zinc-binding dehydrogenase |
| **PPR_1** | 4.05E-15 | 1.40E-12 | 2.48 | 92 | 1,451 | 1,980 | 77,362 | PF12854 | Repeat | PPR repeat |
| **S_locus_ glycop** | 4.78E-15 | 1.53E-12 | 6.08 | 30 | 1,451 | 263 | 77,362 | PF00954 | Domain | S-locus glycoprotein domain |
| **PAN_2** | 1.08E-14 | 3.20E-12 | 6.14 | 29 | 1,451 | 252 | 77,362 | PF08276 | Domain | PAN-like domain |
| **DUF3403** | 2.72E-14 | 7.52E-12 | 9.61 | 20 | 1,451 | 111 | 77,362 | PF11883 | Family | Domain of unknown function (DUF3403) |
| **LRRNT_2** | 4.71E-11 | 1.22E-08 | 3.25 | 42 | 1,451 | 689 | 77,362 | PF08263 | Family | Leucine rich repeat N-terminal domain |
| **LRR_1** | 1.26E-10 | 3.09E-08 | 2.44 | 64 | 1,451 | 1,400 | 77,362 | PF00560 | Repeat | Leucine Rich Repeat |
| **B_lectin** | 2.13E-10 | 4.92E-08 | 4.12 | 29 | 1,451 | 375 | 77,362 | PF01453 | Domain | D-mannose binding lectin |
| **F-box-like** | 2.97E-10 | 6.50E-08 | 4.85 | 24 | 1,451 | 264 | 77,362 | PF12937 | Domain | F-box-like |
| **PPR_2** | 1.28E-04 | 2.65E-02 | 1.51 | 85 | 1,451 | 2,994 | 77,362 | PF13041 | Repeat | PPR repeat family |
| **LRR_8** | 2.79E-04 | 5.52E-02 | 1.68 | 51 | 1,451 | 1,616 | 77,362 | PF13855 | Repeat | Leucine rich repeat |

**Table S6** (2 pages)**.** Within non-tandemly duplicated genes, top hypergeometrically-enriched Pfam domains for those genes participating in at least two SSB-supporting gene pairs. The enrichment set contains 955 PCGs and the background 23,174. Listed Pfams have Benjamini-Hochberg FDR-adjusted *q*-value < 0.05, or hypergeometric *p*-value < 0.0005. Tandemness is defined via global amino acid identity ≥ 30%.

| *Pfam short name* | *Hgeo. p-value* | *BH FDR q-value* | *Obs./ expect* | *Subset has…* | *in:* | *Bkgnd. has…* | *in:* | *Pfam accn.* | *Pfam type* | *Pfam long name* | *Note* |
|---|---|---|---|---|---|---|---|---|---|---|---|
| **AP2** | 6.78E-13 | 2.82E-09 | 5.52 | 26 | 1,920 | 103 | 42,020 | PF00847 | Domain | AP2 domain | Transcription factor |
| **WRKY** | 1.90E-08 | 3.95E-05 | 5.17 | 17 | 1,920 | 72 | 42,020 | PF03106 | Domain | WRKY DNA -binding domain | Transcription factor |
| **ATP-synt_C** | 2.34E-07 | 2.27E-04 | 16.41 | 6 | 1,920 | 8 | 42,020 | PF00137 | Family | ATP synthase subunit C | Enzyme |
| **Roc** | 2.73E-07 | 2.27E-04 | 4.90 | 15 | 1,920 | 67 | 42,020 | PF08477 | Domain | Ras of Complex, Roc, domain of DAPkinase | Signal transduction |
| **DUF4050** | 2.34E-07 | 2.27E-04 | 16.41 | 6 | 1,920 | 8 | 42,020 | PF13259 | Family | Protein of unknown function (DUF4050) | Unknown |
| **Ras** | 7.37E-07 | 5.10E-04 | 4.56 | 15 | 1,920 | 72 | 42,020 | PF00071 | Domain | Ras family | Signal transduction |
| **Myb_DNA-binding** | 2.14E-06 | 1.16E-03 | 2.46 | 33 | 1,920 | 294 | 42,020 | PF00249 | Domain | Myb-like DNA-binding domain | Transcription factor |
| **zf-Dof** | 2.23E-06 | 1.16E-03 | 8.34 | 8 | 1,920 | 21 | 42,020 | PF02701 | Family | Dof domain, zinc finger | Transcription factor |
| **zf-C3HC4_2** | 2.94E-06 | 1.35E-03 | 3.17 | 21 | 1,920 | 145 | 42,020 | PF13923 | Domain | Zinc finger, C3HC4 type (RING finger) | Transcription factor |
| **Hpt** | 4.35E-06 | 1.80E-03 | 21.89 | 4 | 1,920 | 4 | 42,020 | PF01627 | Family | Hpt domain | Signal transduction |
| **RRM_5** | 5.09E-06 | 1.92E-03 | 6.57 | 9 | 1,920 | 30 | 42,020 | PF13893 | Domain | RNA recognition motif (a.k.a. RRM/RBD/RNP domain) | RNA binding |
| **zf-C3HC4** | 6.90E-06 | 2.00E-03 | 2.76 | 24 | 1,920 | 190 | 42,020 | PF00097 | Domain | Zinc finger, C3HC4 type (RING finger) | Transcription factor |
| **zf-RanBP** | 7.14E-06 | 2.00E-03 | 7.30 | 8 | 1,920 | 24 | 42,020 | PF00641 | Domain | Zn-finger in Ran binding protein and others | Transcription factor |
| **Myb_DNA-bind_6** | 7.69E-06 | 2.00E-03 | 2.68 | 25 | 1,920 | 204 | 42,020 | PF13921 | Domain | Myb-like DNA-binding domain | Transcription factor |
| **zf-C3HC4_3** | 7.70E-06 | 2.00E-03 | 4.31 | 13 | 1,920 | 66 | 42,020 | PF13920 | Domain | Zinc finger, C3HC4 type (RING finger) | Transcription factor |
| **HCO3_cotransp** | 6.58E-06 | 2.00E-03 | 10.94 | 6 | 1,920 | 12 | 42,020 | PF00955 | Family | HCO3- transporter family | Transporter |
| **Abhydrolase_2** | 2.09E-05 | 5.11E-03 | 17.51 | 4 | 1,920 | 5 | 42,020 | PF02230 | Domain | Phospholipase/Carboxylesterase | Enzyme |
| **EamA** | 2.31E-05 | 5.33E-03 | 4.97 | 10 | 1,920 | 44 | 42,020 | PF00892 | Family | EamA-like transporter family | Transporter |
| **Pkinase_Tyr** | 5.11E-05 | 1.12E-02 | 1.68 | 63 | 1,920 | 822 | 42,020 | PF07714 | Domain | Protein tyrosine kinase | Signal transduction |
| **Pkinase** | 5.41E-05 | 1.12E-02 | 1.69 | 61 | 1,920 | 790 | 42,020 | PF00069 | Domain | Protein kinase domain | Signal transduction |
| **DUF1218** | 7.26E-05 | 1.31E-02 | 9.95 | 5 | 1,920 | 11 | 42,020 | PF06749 | Family | Protein of unknown function (DUF1218) | Cell wall |
| **SBP** | 7.23E-05 | 1.31E-02 | 7.72 | 6 | 1,920 | 17 | 42,020 | PF03110 | Domain | SBP domain | Transcription factor |
| **Na_Ca_ex** | 7.23E-05 | 1.31E-02 | 7.72 | 6 | 1,920 | 17 | 42,020 | PF01699 | Family | Sodium/calcium exchanger protein | Transporter |
| **Pec_lyase_N** | 9.53E-05 | 1.52E-02 | 21.89 | 3 | 1,920 | 3 | 42,020 | PF04431 | Family | Pectate lyase, N-terminus | Cell wall |
| **GSDH** | 9.53E-05 | 1.52E-02 | 21.89 | 3 | 1,920 | 3 | 42,020 | PF07995 | Domain | Glucose/Sorbosone dehydrogenase | Enzyme |
| **V-SNARE** | 9.53E-05 | 1.52E-02 | 21.89 | 3 | 1,920 | 3 | 42,020 | PF05008 | Family | Vesicle transport v-SNARE protein N-terminus | Transporter |
| **Pec_lyase_C** | 1.20E-04 | 1.79E-02 | 9.12 | 5 | 1,920 | 12 | 42,020 | PF00544 | Domain | Pectate lyase | Cell wall |
| **zf-C2H2_6** | 1.21E-04 | 1.79E-02 | 3.82 | 11 | 1,920 | 63 | 42,020 | PF13912 | Domain | C2H2-type zinc finger | Transcription factor |

| ***Pfam short name*** | ***Hgeo. p-value*** | ***BH FDR q-value*** | ***Obs./ expect*** | ***Subset has…*** | ***in:*** | ***Bkgnd. has…*** | ***in:*** | ***Pfam accn.*** | ***Pfam type*** | ***Pfam long name*** | ***Note*** |
|---|---|---|---|---|---|---|---|---|---|---|---|
| **Gtr1_RagA** | 1.63E-04 | 2.33E-02 | 5.67 | 7 | 1,920 | 27 | 42,020 | PF04670 | Domain | Gtr1/RagA G protein conserved region | Signal transduction |
| **DPBB_1** | 2.01E-04 | 2.78E-02 | 6.57 | 6 | 1,920 | 20 | 42,020 | PF03330 | Domain | Lytic transglycolase | Enzyme |
| **Glyco_hydro_42** | 2.62E-04 | 3.51E-02 | 10.94 | 4 | 1,920 | 8 | 42,020 | PF02449 | Domain | Beta-galactosidase | Enzyme |
| **Rer1** | 3.68E-04 | 4.13E-02 | 16.41 | 3 | 1,920 | 4 | 42,020 | PF03248 | Family | Rer1 family | Membrane |
| **Remorin_N** | 3.68E-04 | 4.13E-02 | 16.41 | 3 | 1,920 | 4 | 42,020 | PF03766 | Family | Remorin, N-terminal region | Membrane |
| **Bap31** | 3.68E-04 | 4.13E-02 | 16.41 | 3 | 1,920 | 4 | 42,020 | PF05529 | Family | B-cell receptor-associated protein 31-like | Membrane |
| **Ribosom_S12_S23** | 3.68E-04 | 4.13E-02 | 16.41 | 3 | 1,920 | 4 | 42,020 | PF00164 | Family | Ribosomal protein S12/S23 | Ribosome |
| **PABP** | 3.68E-04 | 4.13E-02 | 16.41 | 3 | 1,920 | 4 | 42,020 | PF00658 | Family | Poly-adenylate binding protein, unique domain | RNA binding |
| **Y_phosphatase2** | 3.68E-04 | 4.13E-02 | 16.41 | 3 | 1,920 | 4 | 42,020 | PF03162 | Domain | Tyrosine phosphatase family | Signal transduction |
| **EF-hand_1** | 4.24E-04 | 4.63E-02 | 2.61 | 16 | 1,920 | 134 | 42,020 | PF00036 | Domain | EF hand | Signal transduction |
| **PAE** | 4.55E-04 | 4.85E-02 | 9.73 | 4 | 1,920 | 9 | 42,020 | PF03283 | Family | Pectinacetylesterase | Cell wall |

**Table S7.** Top 20 most abundant Pfam domains across genes in *Q. lobata* and from selected other plant species as background information. Counts are the number of genes with one or more of each Pfam domain. Proteome counts suggest number of respective genome project-predicted protein-coding genes for each species. **Red bold type** is largest value among the six tree species for each row. *Q. lobata* data is from our annotation via InterproScan version 5.34-73.0 (https://www.ebi.ac.uk/interpro/about/interproscan/). Non-*Q. lobata* data taken from http://pfam.xfam.org/ (Pfam 33.1, released May 2020).

| | **Comparison tree group** | | | | | | **Selected species for comparison** | | | | | |
|---|---|---|---|---|---|---|---|---|---|---|---|---|
| **Species** | *Quercus lobata (valley oak)* | *Eucalyptus grandis* (Flooded gum) | *Juglans regia* (English walnut) | *Populus trichocarpa* (Western balsam poplar) | *Prunus persica* (Peach) | *Theobroma cacao* (Cacao) | *Amborella trichopoda* | *Arabidopsis thaliana* (Mouse-ear cress) | *Solanum lycopersicum* (Tomato) | *Oryza sativa* subsp. *indica* (Rice) | *Vitis vinifera* (Grape) | *Zea mays* (Maize) |
| **Proteome count** | 39373 | 44149 | 45533 | 53333 | 38726 | 40614 | 27369 | 39359 | 34634 | 37383 | 29903 | 99234 |
| **Pfam Domain:** | | | | | | | | | | | | |
| **Protein kinase domain (PF00069)** | 1287 | **1743** | 1396 | 1501 | 1043 | 964 | 446 | 1001 | 717 | 953 | 824 | 2813 |
| **NB-ARC domain (PF00931)** | **1031** | 795 | 421 | 681 | 472 | 294 | 119 | 318 | 238 | 481 | 347 | 257 |
| **Leucine rich repeat (PF13855)** | 851 | **1003** | 654 | 903 | 534 | 530 | 223 | 358 | 311 | 405 | 438 | 568 |
| **Protein tyrosine kinase (PF07714)** | 790 | 1001 | 815 | **1073** | 621 | 596 | 215 | 630 | 363 | 460 | 487 | 1140 |
| **Leucine rich repeat N-terminal domain (PF08263)** | **679** | 614 | 473 | 535 | 362 | 379 | 136 | 282 | 266 | 344 | 233 | 466 |
| **PPR repeat family (PF13041)** | **674** | 534 | 562 | 632 | 561 | 538 | 549 | 449 | 423 | 405 | 505 | 607 |
| **PPR repeat (PF01535)** | **669** | 499 | 512 | 546 | 517 | 493 | 493 | 450 | 391 | 408 | 476 | 600 |
| **F-box domain (PF00646)** | **541** | 210 | 171 | 221 | 278 | 213 | 103 | 654 | 209 | 375 | 99 | 187 |
| **Cytochrome P450 (PF00067)** | 507 | **614** | 408 | 447 | 328 | 345 | 234 | 326 | 309 | 383 | 385 | 413 |
| **Rx N-terminal domain (PF18052)** | **489** | 144 | 125 | 197 | 183 | 156 | 33 | 24 | 63 | 388 | 152 | 190 |
| **Leucine Rich Repeat (PF00560)** | **448** | 414 | 262 | 316 | 149 | 199 | 69 | 149 | 123 | 155 | 154 | 118 |
| **TIR domain (PF01582)** | **415** | 426 | 246 | 264 | 183 | 25 | 25 | 250 | 39 | 0 | 77 | 3 |
| **zinc-binding in reverse transcriptase (PF13966)** | **410** | 2 | 187 | 2 | 29 | 96 | 1 | 25 | 21 | 70 | 12 | 41 |
| **Domain of unknown function (DUF4283) (PF14111)** | **408** | 45 | 187 | 96 | 30 | 98 | 18 | 23 | 39 | 33 | 8 | 9 |
| **D-mannose binding lectin (PF01453)** | **355** | 321 | 191 | 276 | 144 | 130 | 39 | 98 | 79 | 128 | 102 | 96 |
| **Reverse transcriptase-like (PF13456)** | 323 | 21 | **446** | 21 | 92 | 299 | 10 | 67 | 43 | 85 | 7 | 12 |
| **UDP-glucoronosyl and UDP-glucosyl transferase (PF00201)** | **300** | 376 | 190 | 241 | 198 | 170 | 124 | 131 | 153 | 186 | 224 | 182 |
| **Myb-like DNA-binding domain (PF00249)** | 275 | 291 | **454** | 445 | 289 | 277 | 123 | 324 | 246 | 230 | 230 | 512 |
| **RNA recognition motif. (a.k.a. RRM, RBD, or RNP domain) (PF00076)** | 260 | 315 | **512** | 478 | 405 | 415 | 179 | 382 | 250 | 254 | 212 | 1166 |
| **S-locus glycoprotein domain (PF00954)** | 251 | **287** | 148 | 235 | 104 | 109 | 26 | 86 | 53 | 108 | 93 | 84 |
